# Drift and isolation drive genomic erosion and island speciation in a lineage of macaques

**DOI:** 10.1101/2025.09.06.674399

**Authors:** Jiwei Qi, Shuhao Liu, Liye Zhang, Zhixin Zhou, Rui Wang, Ying Shen, Yang Teng, Mingyi Zhang, Gaoming Liu, Chunyan Hu, Xiaochen Wang, Qian Zhao, Dilina Rusitanmu, Yilai You, Zhijin Liu, Xuming Zhou, Jeffrey Rogers, Christian Roos, Ming Li

**Affiliations:** State Key Laboratory of Animal Biodiversity Conservation and Integrated Pest Management, Institute of Zoology, Chinese Academy of Sciences, Beijing 100101, China; College of Life Sciences, Hebei University, Baoding, Hebei 071002, China; Gene Bank of Primates, German Primate Center, Leibniz Institute for Primate Research, 37077 Göttingen, Germany; Guangdong Institute of Zoology, Guangdong Academy of Sciences, Guangzhou, Guangdong Province, China; Department of Reproductive Medicine, NHC Key Laboratory of Healthy Birth and Birth Defect Prevention in Western China First People’s Hospital of Yunnan Province, Kunming, China; College of Life Science, Capital Normal University, Beijing, China; Administration Bureau of Qi’ao–Dangang Islands Provincial Nature Reserve, Zhuhai, Guangdong, China; Human Genome Sequencing Center and Department of Molecular and Human Genetics, Baylor College of Medicine, Houston, TX, USA

**Keywords:** Macaque, Island speciation, Bottleneck, Transgression, Drift, Purging selection

## Abstract

Allopatric speciation, especially on large islands and archipelagos, is a significant driver of evolutionary diversification, as geographic isolation fosters the independent evolution of populations^1,2^. In these isolated populations, lineage sorting and genetic drift dominate, accelerating allele fixation and reducing shared genetic variation^3,4^. Here, we investigated how sea-level transgression during the Early Holocene triggered rapid speciation in large vertebrates by studying macaques isolated on Dangan Island (DGD), located just 30 km from present-day Hong Kong. Whole-genome sequencing revealed that ∼10,000 years of isolation drove the macaques’ evolution into a distinct species, as indicated by pronounced genomic divergence (mean *F*st > = 0.462 vs. mainland), 1.94 million lineage-specific variants, and complete ancestral differentiation with no evidence of post-isolation gene flow. A severe demographic collapse (effective population size, Ne ≈ 40) led to substantial genomic erosion (65.8% loss of genetic diversity). Paradoxically, this also enhanced resilience through drift-mediated genetic triage. Increased homozygosity exposed and purged lethal recessive alleles in lipid metabolism pathways (68% reduction in genetic load), while simultaneously fixing mildly deleterious variants—such as a splice-site mutation in *SKAP2*—thereby generating a form of genomic “burden” alongside rapid immune adaptation via 251 fixed missense mutations. These findings demonstrate that island isolation can drive vertebrate speciation within a few thousand years, with genetic drift playing a dominant role in shaping genomic architecture. Accordingly, conservation strategies should prioritize monitoring loss-of-function (LoF) variants in essential pathways and prescreening for deleterious allele combinations between donors and recipients prior to implementing genetic rescue in small, drift-sensitive populations.

## Introduction

Climatic fluctuation is a key factor in driving speciation^5^. Several taxa, such as lions, bears, lemmings, and birds, have experienced relatively recent speciation events^6–10^. Isolation in interglacial refugia is a primary mechanism that facilitates speciation. The early Holocene period coincided with the final stage of postglacial global warming which was marked by rapid changes in global moisture availability and ecosystem biomass^11,12^. The continued melting of polar ice sheets and glaciers resulted in a rapid rise in sea level^13,14^, causing widespread marine inundation of the upper continental shelves^15^. Offshore land bridges were submerged through transgression, resulting in the isolation of many islands and promoting geographic separation^15^.

Allopatric speciation is a primary mechanism that drives evolutionary diversification^1,2^. This process is especially accelerated in small isolated populations, where genetic lineage sorting and genetic drift tend to dominate, promoting allele fixation and reducing shared genetic variation^3,4^. Genetic drift promotes divergence even in the absence of strong selection^16^. Moreover, a limited effective population size increases the risk that recessive deleterious mutations become homozygous, thereby resulting in inbreeding depression^16–18^. For example, the extinction of wooly mammoths on Wrangel Island may have occurred due to the accumulation of genetic load^18,19^. These processes, coupled with ecological heterogeneity and restricted gene flow, facilitate rapid diversification and render islands natural laboratories for studying speciation dynamics and genomic differentiation.

High-throughput sequencing has been used to examine the evolutionary history of island species^16,18,20,21^, and to trace speciation trajectories and determine the effects of genetic erosion^21–25^. The continental shelf in Southeast Asia is home to numerous primate species endemic to single islands, which is exemplified by those on the Mentawai islands or Cat Ba^26–30^. Understanding the accumulation of deleterious mutations and the erosion of genetic diversity through genomic studies is necessary for informing and guiding effective conservation strategies for these invaluable island endemics.

The genus *Macaca* (macaques) contains at least 23 species^31^. Of these, the rhesus macaque (*Macaca mulatta*) is the most widely distributed nonhuman primate species worldwide^32^. Its native range spans East Asia, the northern part of Southeast Asia, and the Indian subcontinent^33^. Within China, rhesus macaques are primarily found in southern regions, where population genomic studies have identified five distinct genetic lineages^33^. Island-dwelling macaques inhabit Hainan Island (subspecies *brevicaudus*), the islands adjacent to Hong Kong, and the Wanshan Archipelago in the Pearl River Delta, particularly Chuan Island (CD) and Dangan Island (DGD)^34^. Morphological studies suggest that these island populations share close affinity with the Hainan subspecies *brevicaudus*^35^. Mitochondrial evidence supports natural colonization from the mainland during the Quaternary period^34^; however, their nuclear genomic composition has not yet been examined.

In this study, we present whole-genome sequencing data for two insular macaque populations from CD and DGD, which are located approximately 18 km and 30 km from the Chinese mainland, respectively. Genomic analyses suggest that the DGD population represents a distinct evolutionary lineage, possibly the 24th species within the genus *Macaca*, originating from an ancestral stock isolated since the Holocene sea-level rise. We systematically assessed the accumulation of deleterious mutations and evaluated the genetic consequences inherent to long-term isolation. Our results highlight the pivotal role of geographic isolation in driving speciation within a relatively short evolutionary time, and emphasize the importance of understanding genetic erosion, primarily driven by genetic drift, for ensuring the long-term viability of small, isolated populations.

## Results

### Genome sequencing and variant calling

We generated whole-genome sequencing data for 51 individuals, containing 30 newly sequenced individuals and 21 previously generated high-depth genomes (Table S1). This collection included 26 insular macaques (six from CD and 20 from DGD) and 25 representatives from recognized *Macaca mulatta* subspecies: 5 *M. m. mulatta*, 8 *M. m. lasiotis*, 7 *M. m. littoralis*, and 5 *M. m. brevicaudus*. The sequencing achieved an average depth of 33.49 ×. Dataset I (Fig. 1a; Table S1) also included an additional 61 published genomes (5 *M. m. mulatta*, 23 *M. m. lasiotis*, 28 *M. m. littoralis*, 5 *M. m. tcheliensis*) to examine the evolutionary trajectories between continental and insular macaques.

**Fig. 1.**
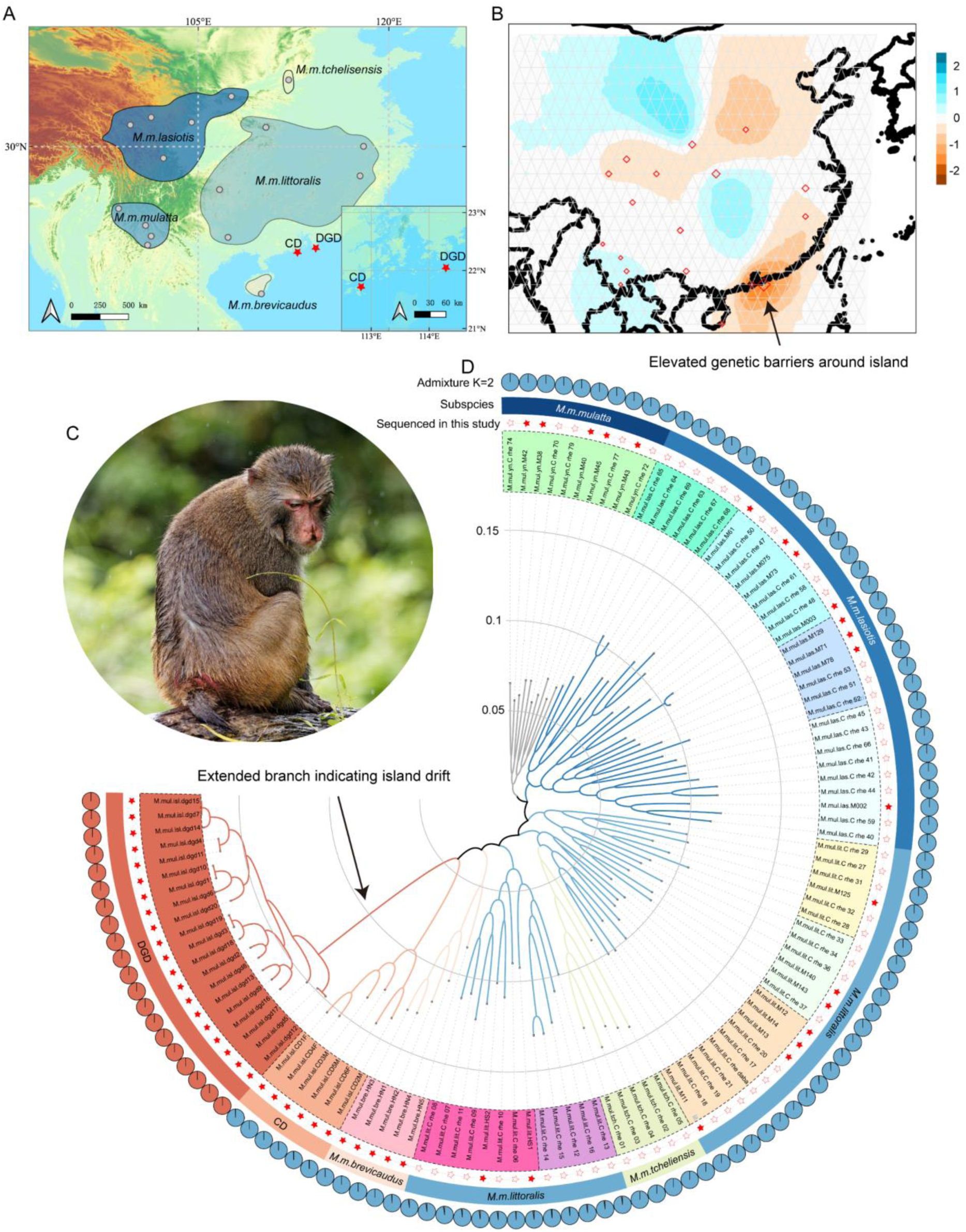
Distribution of *Macaca mulatta* subspecies in China, Effective Migration Surfaces (EEMS), and Phylogenetic Analysis. **(A) Distribution of *Macaca mulatta* subspecies in China.** The islands sampled in this study are marked with a red star: Chuan Island (CD) and Dangan Island (DGD) in the South China Sea. Solid lines indicate hypothesized boundaries within subspecies. **(B) Effective Migration Surfaces (EEMS) analysis.** The red regions show the elevated barriers to gene flow (reduced migration), whereas the blue regions denote lower genetic resistance (facilitated migration). The 30-km-wide strait surrounding the islands (black arrow) exhibits strong migration resistance (red zone) and severely restricts gene flow. **(C) Photograph of a Dangan Island macaque.** This individual shows a markedly shortened tail, which was historically documented as 4.5 inches/11.43 cm in records from 1866^58^, and a congenital cleft lip. **(D) Phylogenomic Circos plot.** The innermost ring shows the phylogenetic tree topology with branch lengths (scale bar indicates genetic distance). Sample labels are colored by geographic origin (matching Table S1). Solid red stars indicate samples newly sequenced or deeply re-sequenced. The fourth ring indicates the subspecies designation. The outer circles shows ancestry proportions at *K* = 2 (red and blue represent ancestral components A and B, respectively), highlighting the distinct red ancestry of DGD.

We established Dataset II by adding ten rhesus macaques of Indian origin, seven additional macaque species, and a baboon outgroup (Table S1) to Dataset I. Sequence alignment to the *M. mulatta* reference genome (Mmul_10, GCA_003339765.3) yielded an average mapping rate of 99.68% and genome coverage of 97.83% (Table S1). Through rigorous variant calling, 71,127,020 autosomal variants in Dataset I and 130,486,137 variants in Dataset II (Table S2) were identified, which were used in subsequent evolutionary analyses.

### Mainland origins, island radiation: Genomic divergence and putative speciation in DGD

We initially gathered genome-wide data from seven species within the genus *Macaca* including ten rhesus macaques of Indian origin (Dataset II) for phylogenetic tree construction, Principal Component Analysis (PCA) analysis, and haplotype network analyses based on mitochondrial DNA (mtDNA). The PCA analysis of autosomal variants revealed distinct phylogeographical differentiation. Specifically, PC1 (17.94%) contrasted the African *M. sylvanus* with Asian macaques (Fig. S1 and S7), whereas PC2 (8.64%) separated the *arctoides*/*sinica* clade from the *mulatta*/*fascicularis* clade. This corroborates earlier phylogenetic findings^36^. Notably, DGD falls within the *mulatta* clade but distant from the Chinese and Indian macaques, similar to the Taiwanese macaque (*M. cyclopis*) and the Japanese macaque (*M. fuscata*). To further determine the phylogenetic position of DGD among macaques, we performed phylogenetic tree reconstruction based on autosomal variants and mtDNA data (Fig. S2). The results indicated that DGD is a derived population from the Chinese mainland lineage that shares a recent common ancestor with CD. This common ancestor also shares a recent but more distant ancestor with the Hainan Island rhesus macaque subspecies *brevicaudus*. Importantly, the autosomal variants results demonstrated a significantly long branch for DGD, suggesting the accumulation of numerous population-specific mutations (Fig. S2). In addition, haplotype network analyses based on mtDNA revealed a geographic clustering that was consistent with CytB and Cox1 gene distributions. This indicates that DGD originated from the rhesus macaque in eastern China (subspecies *littoralis*) and is closely related to *M. m. brevicaudus* and CD (Fig. S3). Moreover, we identified 11 DGD-specific new derived mutation sites among all 13 mtDNA protein-coding genes, three of which are nonsynonymous mutations (Cytb: M306V; Nad1: V113A; Nad4: P429S).

The autosomal variants and mtDNA sequences clarified the evolutionary position of DGD as a population derived from the Chinese continent and sharing a common ancestor with *M. m. brevicaudus* and CD. We next modified our study framework to include only dataset I (comprising island and Chinese mainland populations) to further explore the population characteristics of DGD and its evolutionary footprints. We performed an Admixture analysis, phylogenetic tree reconstruction, PCA analyses, and fineSTRUCTURE to further examine the evolutionary relationships. In addition, we also calculated *F*st and analyzed shared alleles to assess the degree of differentiation and validate common ancestry. Using Estimated Effective Migration Surfaces (EEMS) we tested for deviations from isolation by distance (See Methods). The Admixture analysis identifyed two distinct genetically homogeneous populations. DGD was separated from all other populations, as indicated by the cross-validation errors when K = 2 (Fig. 1c, Fig. S4, and S5). Notably, at K = 2, we observed complete differentiation between DGD (red) and all other components (blue) (Fig. 1c, Fig. S4, and S5).

Furthermore, the phylogenetic tree and PCA based on Dataset I yielded similar results to those based on Dataset II. PC1 distinctly positioned DGD on the left side with a resolution of 6.32%, whereas the continental populations, including *M.m. brevicaudus* and the CD population, were distinctly different (Fig. 2a). The fineSTRUCTURE coancestry matrix (Fig. S6) displayed clear population structuring between DGD and the remaining populations, which suggests lower levels of shared coancestry between the groups.

**Fig. 2.**
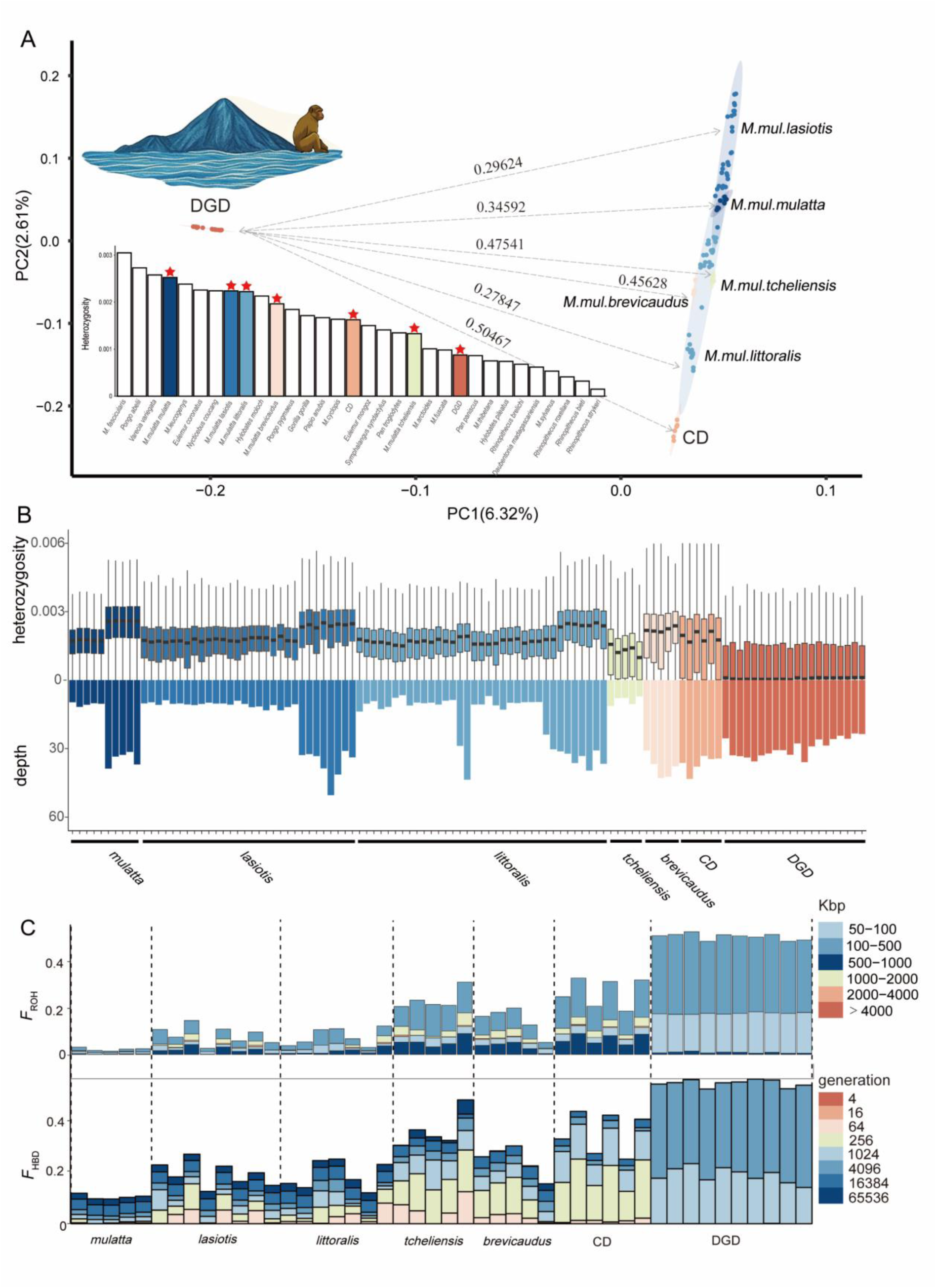
Genomic Landscape of Dangan Island Macaques. **(A) Principal Component Analysis (PCA).** PCA reveals pronounced genetic divergence, with the Dangan Island (DGD) population clustering separately (left) from the mainland and other island populations (right) along PC1. Dashed lines indicate pairwise *F*st values that quantify this divergence. The embedded comparative analysis suggests that DGD’s heterozygosity falls within the low level of primate species surveyed, indicating severe loss of genetic diversity. **(B) heterozygosity plots.** Individual-level variation shows consistently low heterozygosity in DGD macaques (upper panel), and is not correlated with sequencing depth (lower panel). **(C) Genomic inbreeding**. Genomic inbreeding metrics confirm extreme inbreeding in DGD: observational-based approaches, PLINK analysis (upper) revealed abundant short ROH segments (50–500 Kbp), whereas model-based approaches, such as the RZooRoH temporal decomposition (lower), indicate intensive HBD segments <2 cM (recent ancestors traced back approximately 1,024 to 4,096 generations).

To evaluate the genetic divergence of the DGD population more deeply, we calculated the gene-wise fixation index *F*st^37^. This revealed a pronounced differentiation between DGD and the Chinese mainland rhesus macaques, with pairwise *F*st values ranging from 0.27 to 0.47 (Fig. S7, Table S3). Extended comparisons were made across the *mulatta* species group (including mainland Chinese *M. mulatta* subspecies, Taiwanese *M. cyclopis*, and Japanese *M. fuscata*), which revealed even higher differentiation (*F*st = 0.27–0.71; Fig. S8, Table S3). Hierarchical analysis identified three tiers of genetic separation. Intra-subspecies differentiation within mainland populations averaged *F*st = 0.076 (SE = 0.043), interspecific divergence within the *mulatta* group reached *F*st = 0.233 (SE = 0.074), whereas DGD exhibited markedly greater divergence from all lineages (mean *F*st = 0.462, SE = 0.173). This differentiation significantly exceeded interspecies-level genetic distances (*P* < 0.001, Wilcoxon rank-sum test; Fig. S8), thus indicating exceptional evolutionary distinctiveness of the DGD population. To identify geographic drivers of differentiation, the EEMS model was applied. The results indicated a pronounced genetic barrier that corresponded to the sea-land interface adjacent to the DGD and CD regions. These islands are located 30 and 18 kilometers from the mainland, respectively. For DGD, this ∼30-km marine strait constitutes a major biogeographic barrier, effectively restricting contemporary gene flow and maintaining genetic isolation in DGD and CD through physical dispersal limitations (Fig. S8).

Because substantial genetic differentiation distinguished DGD from the recognized species and continental rhesus macaque populations, we performed formal species delimitation analyses using Bayesian Phylogenetics and Phylogeography frameworks^38^. Three independent datasets were analyzed: (1) 1,000 nuclear DNA (nDNA) loci, (2) mitochondrial genomes (mtDNA), and (3) 500 nuclear coding genes. Two distinct species delimitation algorithms (rj1, rj0) were used under a unified guide tree topology (A10). The analyses converged on identical speciation boundaries, unanimously supporting the recognition of two discrete species (DGD vs. *M. mulatta*) with maximum posterior probability support (pp = 1; Table S4). This consensus emerged consistently across all data partitions (nDNA/mtDNA/coding genes) and algorithmic approaches (rj1/rj0), indicating a robust phylogenetic monophyly of the DGD lineage.

### Extreme diversity loss, runs of homozygosity (ROH) associated with land bridge disappearance, and the consequences of drift

Genome-wide sliding window analyses revealed considerable depletion of heterozygosity across insular populations, with the diminutive DGD population (13.2 km²) showing extreme genomic monomorphism (0.8 SNPs/1000 bp), which is half of level observed among their CD conspecifics (Welch’s t-test, p < 0.001), and just one-third of continental subspecies (*littoralis*, *lasiotis*, and *mulatta*; Welch’s t-test, p < 0.001; Fig. 2b). This genomic heterozygosity puts the DGD population at the periphery of the primate diversity metrics, but the heterozygosity of DGD does exceed that of the endangered golden snub-nosed monkey (*Rhinopithecus roxellana*: 0.4/1000 bp^39,40^) (Fig. 2a). We also quantified nucleotide diversity across all study groups. This independent metric consistently revealed markedly decreased genetic diversity in DGD (Welch’s t-test, p < 0.001; Fig. S9). The strikingly low genetic diversity observed in DGD likely results from a historical population bottleneck, with this depletion being further exacerbated by contemporary demographic constraints, specifically, an insular population persisting at critically low census numbers.

Complementary analyses of linkage disequilibrium (LD) patterns, which also reflect demographic history, revealed slower LD decay rates for the DGD and CD populations compared with the continental groups (Fig. S10). Moreover, the DGD population shows a narrow but extended distribution of Tajima’s D values (Fig. S11), which suggests localized signals of purifying selection and demographic contraction across different genomic regions. Importantly, the site frequency spectrum (SFS) exhibits a characteristically flattened profile (Fig. S29), marked by a depletion of low-frequency deleterious mutations and the accumulation of intermediate-frequency variants. This genomic signature confirms the dominance of genetic drift over selection in shaping the evolutionary trajectory of the DGD population.

Isolated insular populations experience elevated inbreeding risks, as evidenced by extended contiguous homozygous regions (Identical-by-descent IBD segments). These genomic signatures are quantifiable biomarkers for assessing inbreeding^26,41^. Here, we estimated the fraction of the genome within IBD segments (*F*_IBD_) using two software approaches: (i) *F*_ROH_ estimated from runs of homozygosity (ROHs) with observational-based approaches, and (ii) *F*_HBD_ estimated from homozygous-by-descent (HBD) segments from model-based approaches RZooRoH. A strong concordance was observed between *F*_ROH_ and *F*_HBD_ estimates across the insular populations (DGD: *F*_ROH_ = 0.48–0.54 vs *F*_HBD_ = 0.48–0.53; CD: *F*_ROH_ = 0.25–0.42 vs *F*_HBD_ = 0.18–0.30). Both metrics, particularly in DGD, were more than twice as high as their mainland counterparts (Fig. 2C). Notably, PLINK-based ROH analysis and RZooRoH HBD modeling revealed an exceptionally high prevalence of short genomic segments in DGDs: >100,000 counts of ROH fragments <500 kb (PLINK) and ∼20% of the genome comprising HBD segments <2 cM (RZooRoH) (Fig. S12). HBD analyses revealed that the recent ancestors of the DGD population may be traced back approximately 1,024 to 4,096 generations (R_k_ = 1,024 − 4,096) (Fig. 2C, Fig. S13 and S14), roughly corresponding to the island isolation chronology (∼9,500 yr BP; generation time: 11 years). To validate the ROH observations, we randomly visualized 20 Mb of HBDs on chromosome 19, which confirmed the number of extremely short HBD fragments in DGD compared with other populations (Fig. S15). The detection of a large number of inbred IBD fragments reflects mating range contraction instantly during marine transgression events ∼9,500 yr BP, which eliminated terrestrial connectivity^13^.

Population bottlenecks, such as those observed in insular macaque populations following land-bridge disappearance, may compromise selection efficacy. Therefore, we examined the ratio of heterozygosity between zero-fold and four-fold degenerate sites (where zero-fold sites exhibit exclusively nonsynonymous mutations, and four-fold sites permit only synonymous mutations). This metric is theoretically increased in small populations because of increased frequencies of deleterious alleles under strong genetic drift and attenuated selection pressure. Confirming this prediction, we identified a negative correlation between the zero-fold/four-fold heterozygosity ratio and neutral heterozygosity. Smaller populations exhibited reduced neutral heterozygosity coupled with elevated zero-fold heterozygosity (Fig. 3B). Notably, DGD represents one of the most extreme cases, and demonstrates substantially amplified levels of putatively deleterious zero-fold heterozygosity. To validate this pattern, we used SnpEff’s variant classification to group high + moderate impact variants (presumably more deleterious) for comparison with low-impact variants. The results showed consistent findings (Fig. S16). Our analysis revealed that stochastic processes supersede natural selection in shaping the DGD lineage, with genetic drift emerging as the dominant evolutionary force following sequential founder effects, and a demographic bottleneck that dramatically depleted standing genetic variation (Fig. S17). Specifically, we observed 1,940,089 fixed variants in DGD (Fig. S17), which is an exceptionally high number that reflects its severe demographic history. These fixed variants caused missense mutations in 2,353 genes (Table S5), with 251 genes acquiring population-specific missense mutations (Table S6). This resulted in potential functional changes in immune-related genes (GO:0002253: activation of immune response, *P. value* = 0.0015; GO:0045087: innate immune response, *P. value* = 0.0009; GO:0051607: defense response to virus, *P. value* = 0.0009), whereas pathways, such as Tyrosine metabolism (mcc00380, *P. value* = 0.0432), Cytoskeleton in muscle cells (mcc04820, *P. value* = 0.0045), and Homologous recombination (mcc03440, *P. value* = 0.0041) were also enriched (Fig. S18 and Table S7).

**Fig. 3.**
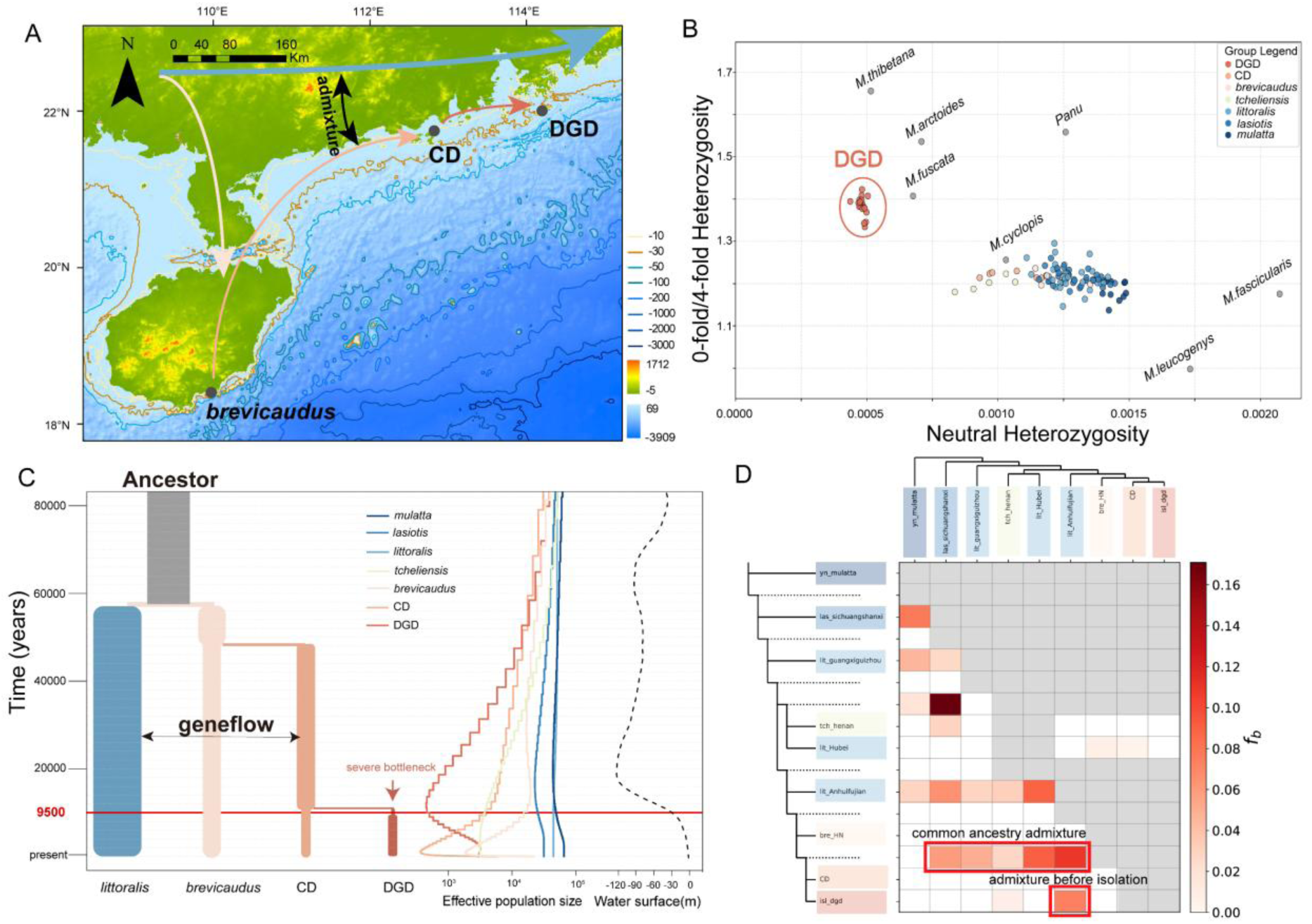
Speciation Scenario and Extreme Drift-Driven Divergence of Dangan Island Macaques. **(A) Paleogeographic divergence trajectory.** Following the split of *M. m. littoralis*, the ancestral *M. m. brevicaudus* (Hainan subspecies) population expanded eastward along the coastline of China. The isolation of Dangan Island (DGD) commenced ∼10 kya during marine transgression events (see also Fig. S40), initiating the independent evolutionary trajectory. The arrows indicate hypothesized migration routes. **(B) Relaxation of purifying selection.** Negative correlation between neutral diversity (proxy for effective population size, Ne) and heterozygosity ratio at zero-versus four-fold degenerate sites. Increased ratios in small populations indicate increased frequencies of deleterious nonsynonymous mutations during strong genetic drift. DGD (red circle) shows extreme outlier status. **(C) Coalescent simulation of speciation scenario.** Left: Best-supported ABC coalescent model (fastsimcoal2) for DGD divergence, which indicates a severe founding bottleneck. Middle: SMC++ reconstruction confirms ancestral Ne ≈ 40 during the isolation period, coinciding with −30 m paleo-sea-level (dash curve; NOAA Paleoclimate Data; see also Fig. S40). **(D) Gene flow analysis.** *f*-branch analysis (Dsuite) detected a significant admixture (*P* < 0.001) between ancestral CD/DGD branches and mainland *littoralis* (red signal). For DGD, integrated evidence from *f3* statistics, TreeMix topology, and coalescent modeling demonstrated pre-isolation continental gene flow, which occurred before island colonization and marine transgression.

### DGD: 9,500 years of insular isolation and population bottlenecks

To reconstruct demographic trajectories, a pairwise sequentially Markovian coalescent (PSMC) analysis was performed to track changes in effective population size (Ne) over time. Genomic comparisons between the mainland and island populations revealed moderately congruent demographic histories, which were characterized by a pronounced collapse in Ne beginning approximately 700,000 yr BP and coincided with the onset of the Naynayxungla Glaciation (NG) (Fig. S19). Recognizing the known limitations of PSMC in reconstructing recent demography, in which parameter sensitivity can generate artifactual peaks^25^, we implemented systematic parameter substitutions that substantially altered the inferred population trajectories. Subsequent analyses revealed moderate Ne recovery in continental populations during the interglacial period separating the NG and Penultimate Glaciation (PG), followed by accelerated contraction postdating the PG (Fig. S19). Because PSMC exhibits reduced power to estimate recent Ne changes (i.e., <20 ka BP ^42^), demographic reconstructions were also performed with SMC++, which is better suited to infer recent demographic histories^43^. SMC++ analyses also supported the demographic expansion between the NG and PG and revealed that population contractions began at the end of the PG (Fig. S20). Furthermore, we identified a dramatic population bottleneck in DGD around 10,000 yr BP, with a Ne ∼ 40 (Fig. S20). To further assess the authenticity of this bottleneck, we used StairwayPlot2 based on the SFS of 20 DGD individuals. The results indicated that the same severe bottleneck occurred around 9,500 yr BP, with a Ne also estimated to be near 40 (Fig. S21). This coincided with the period of dramatic sea-level change and the disappearance of the land bridge. Nonetheless, in the absence of human assistance, the DGD population subsequently rebounded to nearly 2,000 based on local ranger observations, despite the drastic reduction in genetic variation.

Historical records indicate the presence of macaques on DGD since 1819, which supports the idea that the population of the island is natural and not artificially introduced^35,44^. To infer the speciation scenario of the DGD macaque and assess potential secondary contact with continental populations, we conducted demographic inference using approximate Bayesian computation (ABC), coalescent-based simulations. After the first round of model comparisons, a phylogenetic relationship consistent with the whole-genome ML phylogenetic tree was determined as optimal (Fig. 3A, S22, and Table S8). This model supports the divergence of the Hainan Island population (*M. m. brevicaudus*) from the mainland population in southern China (*M. m. littoralis*), followed by eastward dispersal and the subsequent divergence of ancestral CD and DGD populations into distinct lineages (Fig. 3A, S22, and Table S8).

Determining whether island populations experienced secondary contact with mainland populations is important to understand island speciation, as genetic exchange can result in genomic homogenization and impede lineage differentiation^3^. To examine this in the present study, an integrated gene flow analysis was performed using TreeMix, *D*-statistics, *f*-branch, and outgroup *f3* statistics. TreeMix analysis with the OptM method revealed two migration edges as optimal (Fig. S23), which indicated no significant secondary contact between contemporary DGD and mainland populations (Fig. S24). Although a potential mixing event was inferred between ancestral DGD populations and *littoralis* on the continent at edges=3, the minimal migration weight (weight = 0.05) shows that if this gene flow occurred, it was extremely limited. Moreover, outgroup *f3* statistics revealed no significant admixture into or out of the DGD population (Z < −2; Fig. S25A, S25B). However, *D*-statistics and *f*-branch analysis, the latter disentangling correlated *f4*-ratio results to assign gene flow to specific internal branches^45^, revealed significant gene flow signals. Low-level gene flow (1.5%; |Z| ≥ 3) was observed between DGD and the *littoralis* subspecies from Fujian and Anhui populations on the southeast coast of China (Fig. S25C), and evidence of admixture (7.2%) was identified between the ancestral branches of CD/DGD and the mainland *littoralis* subspecies (Fig. 3D). Taken together, these results indicate that genetic exchange occurred between the ancestors of the CD/DGD and mainland *littoralis* subspecies before isolation from the mainland. Importantly, after the disappearance of the land bridge, the DGD population experienced no detectable admixture with any other populations, which represents a case of complete insular isolation, and suggests speciation (Fig. 3A, D). This conclusion is further supported by ABC simulations with *fastsimcoal2* (round 2; Fig. S22, Table S9), which models divergence scenarios with gene flow and consistently identified the ancestral CD/DGD-*littoralis* exchange as the optimal fit. By incorporating historical demographic dynamics (round 3), we simulated speciation scenarios comodeling bottlenecks and gene flow (Fig. S22). Under the best-fit model, DGD underwent a severe genetic bottleneck postdivergence from CD with no subsequent gene flow (Fig. 3C, S22, Table S10). Partitioning the genome into 50 segments for SFS simulations based on this optimal scenario yielded 95% confidence intervals for isolation timing: 915–1,291 generations (Table S11; generation time: 11 years). This estimate is consistent with the geological isolation of DGD Island from the mainland during the sea-level rise at ≈9,500 yr BP (Fig. 3C). Consequently, our analyses reject recent human-mediated establishment (past 500 years), and instead, identify DGD as a distinct lineage isolated naturally for ∼1,000 generations.

### A combination of genetic drift and purging of deleterious mutations has resulted in a reduction of genetic load in DGD

Theory suggests that such a small population is likely to have accumulated deleterious mutations through genetic drift^46^. To explore the genetic history of this isolated insular population, the loss of genetic diversity was first assessed in each of the founder populations. We found that the three-step founder effects in DGD resulted in a sequential loss of genetic diversity, including 14.1% from *littoralis* to *brevicaudus*, 18.6% from *brevicaudus* to CD, and 51.1% from CD to DGD (Fig. S26A). Sampling effects and genetic drift collectively resulted in a total loss of 65.8% in genetic diversity from the mainland population *littoralis* to the insular DGD population across all chromosomes.

Based on loss-of-function (LoF) mutations (identified as transcript ablation, splice donor, splice acceptor, stop gained, frameshift, insertion, deletion, or splice region variants), relative genetic load (as measured by Rxy) differed between the ancestral populations (Fig. 4A). Although the insular DGD population exhibited the highest inbreeding levels (Fig. 2C), it showed a significantly reduced LoF load compared with all ancestral populations. For the derived missense mutations, whose effects are less clear but are often assumed to be more mildly deleterious^46^, load was also significantly reduced, except in DGD versus CD, in which a slight increase was observed. Furthermore, the mainland population maintained a higher proportion of LoF alleles in the heterozygous state compared with the DGD population, whereas many were converted to the homozygous state in DGD (Fig. 4B). This suggests that many of these mutations are primarily deleterious in the homozygous state, which is consistent with the theoretical predictions^47^. Significantly fewer LoF alleles within ROH were also observed compared with heterozygous genomic regions (Figure S27). This difference was 68% smaller in the DGD island population, indicating that inbreeding may have facilitated purging of a substantial proportion of severely deleterious, recessive LoF alleles through exposure in the homozygous state.

**Fig. 4.**
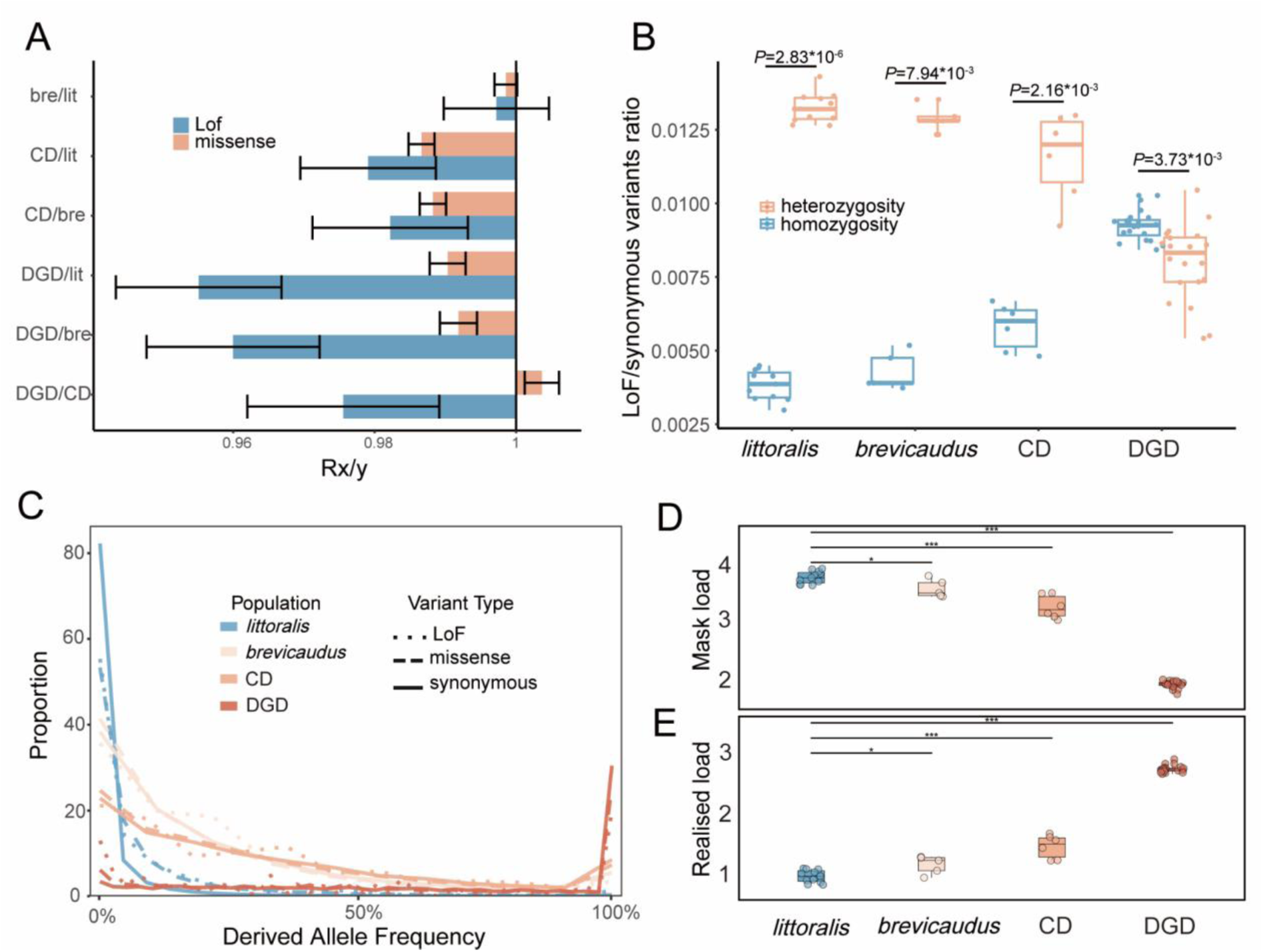
Genomic Erosion in the Dagan Island Macaque. **(A) Rx/y ratio analysis.** Relative number of derived alleles at the LOF (red) and missense (blue) sites that frequently occur in one population, but not in another. The error bars represent ± 2 SDs. **(B) Purifying selection efficacy in the homozygous tracts.** The LoF-to-synonymous variant ratios in the homozygous (blue) vs. non-homozygous (red) genomic regions. Horizontal bars: population means, P-values from the Kolmogorov-Smirnov tests. **(C) Derived allele frequency (DAF) spectra. The** DAF distributions for damaging (LoF/missense; solid line and dashed line) and neutral (synonymous; dotted line) mutations. DGD shows a rightward shift, which indicates the drift-driven fixation of deleterious alleles. **(D) Masked genetic load.** The sum of the GERP scores (>4) for deleterious heterozygous alleles normalized per individual. (one-way ANOVA test; *** p < 0.001, * p < 0.05) **(E) Realized genetic load.** The sum of the GERP scores (>4) for the deleterious homozygous alleles normalized per individual. (One-way ANOVA test; *** p < 0.001, * p < 0.05).

To determine the deleterious effect of these sites, we estimated individual masked and realized loads using autosomal polymorphism data and genomic evolutionary rate profiling (GERP) scores. All of the island groups (compared with mainland *littoralis*) showed pronounced masked-to-realized load transitions under elevated inbreeding (Fig. 4D and 4E), which is consistent with previous observations^22^. Specifically, in DGD, widespread heterozygote-to-homozygote conversion caused a substantial decrease in masked load, but significant increase in realized load (mask load decreased by 52.8%, one-way ANOVA test, *p* < 0.001; realized load increased by 190.6%, one-way ANOVA test, *p* < 0.001) (Fig. 4D and 4E). The realized load actually exceeds the masked load (mask/realized = 1.7348/2.9815). This indicates that island bottlenecks accelerate the purging process of genetic load through recessive allele exposure to selection. Importantly, depressed deleteriousness at homozygous loci (Fig. S28) indicates strong purifying selection against highly deleterious homozygous variants in surviving individuals. Moreover, the DGD mutations are very sparse (Fig. S26B and S26C); however, these mutations, particularly the singleton mutations, have outrageous Ts/Tv ratios (transition-to-transversion = 0.25), which indicates that they underwent strong purifying selection^48^.

We observed that under the combination of drift and purifying selection, the bottlenecked DGD population exhibits extensive fixation of synonymous (30%), missense (26%), and LoF variants (20%) (Fig. 4C). This is accompanied by a significant rightward shift in the derived allele frequency spectrum and a high proportion of fixed derived variants (Fig S29). These results indicate that within a system governed by genetic drift, the fixation of mutations appears to occur randomly on average.

To characterize the functional impact of LoF mutations, we generated two gene sets. Geneset I: Standing LoF variation in ancestral population and absent in DGD, which identified sites never homozygous in ancestral populations (*littoralis*, *brevicaudus*, CD) with moderate allele frequencies that were completely purged in DGD. We propose that these mutations caused homozygous lethality during inbreeding, driving elimination through purifying selection of DGD’s population bottleneck. Geneset II: Standing LoF variation in ancestral population and drift-fixed in DGD, which likely underwent fixation through a prolonged genetic drift and avoided purifying selection, which potentially compromised long-term population fitness. The analysis of 1,042 genes in Geneset I (Table S12) revealed strong enrichment for lipid metabolism (GO:0016042: lipid catabolic process, *P. value* = 0.0001; GO:0006631: fatty acid metabolic process, *P. value* = 0.0050; GO:0006635: fatty acid beta-oxidation, *P. value* = 0.0025; mcc00760: fatty acid degradation, *P. value* = 0.0065) and energy-related functions (GO:0019752: carboxylic acid metabolic process, *P. value* = 0.0191; mcc00760: Nicotinate and nicotinamide metabolism, *P. value* = 0.0445) (Fig. 5B, Table S13). Strong selection of lipid metabolism genes likely occurred ∼9,500 yr BP during post-isolation food scarcity. During cold stress, increased thermoregulatory demands induced lipolysis and circulating FFAs^49,50^. A caloric deficit with impaired lipid metabolism would cause fatal ketosis/dyslipidemia, as fatty acid-derived ketone bodies (e.g., β-hydroxybutyrate) serve as starvation energy sources^51^. Key enzymes in these pathways (e.g., *AKT1*, *GCDH*, *ACADL*, *CPT1C*, *HSD17B4*, and *BDH2*) were enriched in Geneset I. Homozygous LoF mutations disrupting these functions may have induced lethal metabolic failure in DGD.

**Fig. 5.**
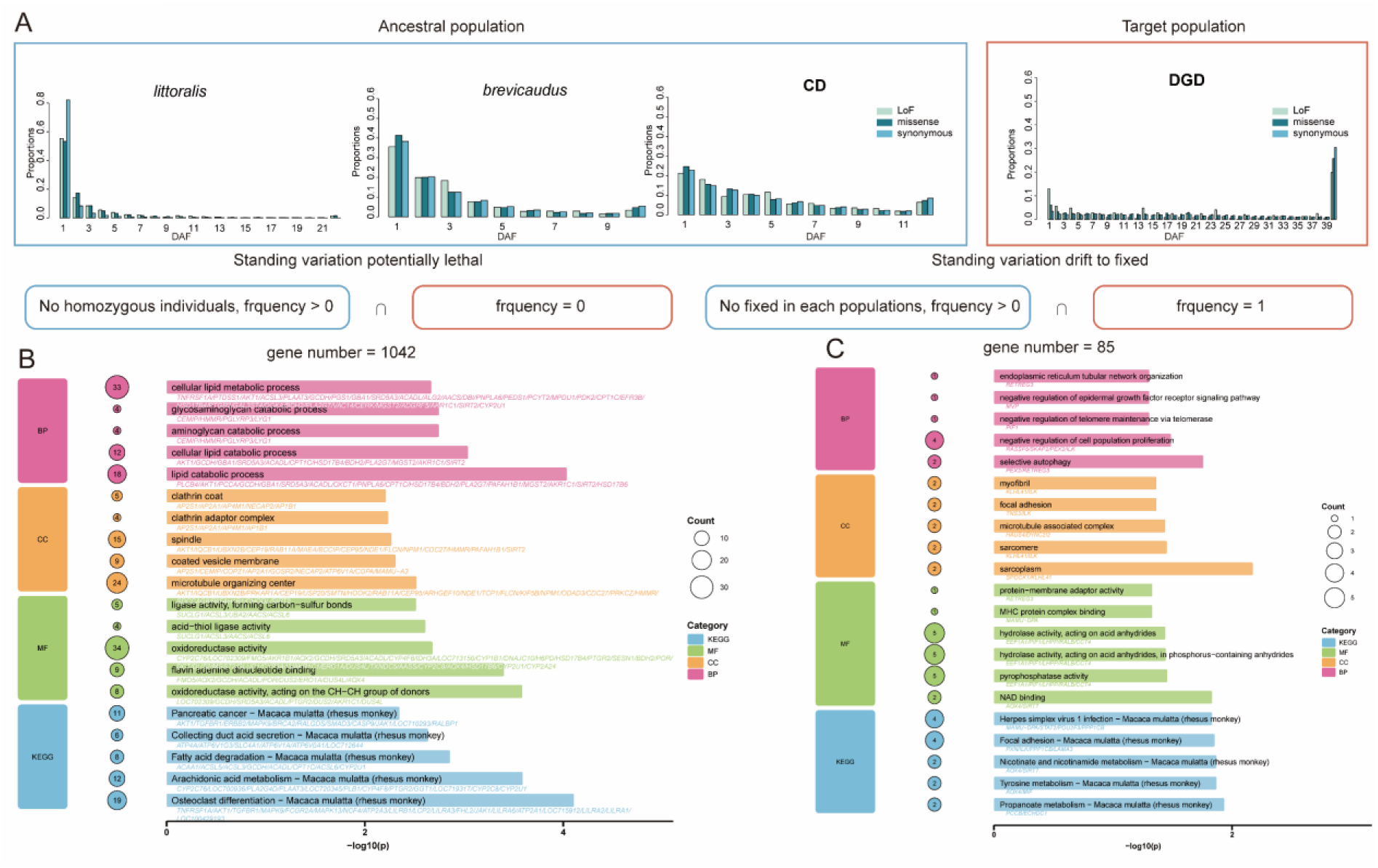
Functional Impact of Genetic Drift on LoF Variation. **(A) Derived allele frequency (DAF) spectra.** The ancestral populations (*littoralis*, *brevicaudus*, CD) versus derived DGD population. The pronounced fixation peak (DAF = 1) in DGD reflects genome-wide deleterious allele fixation through genetic drift. **(B)** Functional enrichment of the purged deleterious variation (Geneset I: 1,042 genes). The LoF variants eliminated in DGD despite moderate ancestral frequencies (0.01 ≤ MAF ≤ 0.5). The recessive lethal mutations purged during inbreeding-induced homozygosity. **(C) Enrichment of drift-fixed deleterious variation (Geneset II: 85 genes).** Ancestral LoF variants fixed in DGD via unopposed genetic drift. These mutations escaped purifying selection, which potentially compromised long-term adaptive potential.

Functional enrichment of Geneset II (85 gene; Fig. 5C, Table S14-15), which may affect DGD health in the long-term, included cell adhesion and cytoskeletal organization (GO:0005925: Focal adhesion, *P. value* = 0.0432; GO:0030017: Sarcomere, *P. value* = 0.0354), metabolic processes (GO:0051287: NAD binding, *P. value* = 0.0148; mcc00640: Propanoate metabolism, *P. value* = 0.0116; mcc00350: Tyrosine metabolism, *P. value* = 0.0135), and negative regulation of cell proliferation (GO:0008285: Negative regulation of cell proliferation, *P. value* = 0.0326; GO:0061912: Selective autophagy, *P. value* = 0.0175). To overcome the limitations in distinguishing ancestral alleles using the Indian-origin Mmul_10 reference (which may contain deleterious variants or misrepresent ancestral states^46^), the genomic data from seven macaque species (Methods) were integrated for robust ancestral/derived allele assignment. After filtering misidentified LoF variants (e.g., double codon mutations [Fig. S30] or high ancestral population frequency suggesting low deleteriousness [Fig. S31]), 34 high-confidence derived deleterious fixed LoF variants were identified that affected 31 genes (Table S16). Applying stringent evolutionary criteria (1) low ancestral frequencies in *M. mulatta* populations, and (2) low frequencies and ancestral homozygosity across seven macaque species, seven putatively ultra-deleterious fixed LoFs (*ZNF790, PPP1CB, ELOA, NDE1, SKAP2, TMEM235, ATP2C2*; Fig. S31–S37) were identified. Although transcriptomic evidence for the DGD population is currently lacking, the LoF of these genes, particularly the splice acceptor mutation in *SKAP2* (containing a *de novo* mutation that has been fixed [Fig. S36]), deserves attention in subsequent conservation practices. Because of their potential to disrupt key biological pathways, we recommend prioritizing these genes during future population monitoring and adaptive management strategies.

## Discussion

### A new macaque species from a near-shore island: genomic evidence of 9,500 years isolation 30 km from a present-day mega-city

During the Late Pleistocene, Chuan Island (CD) and Dangan Island (DGD) formed part of the hill systems in mainland southern China (Fig. S40)^32^. Early Holocene sea-level rise submerged low-lying valleys, transforming hilltops into isolated islands^13,52^. This markedly accelerated over time, increasing from 16.4 ± 6.1 mm/yr at 10,500 yr BP to 33.0 ± 7.1 mm/yr at 9,500 yr BP, before decreasing to 8.8 ± 1.9 mm/yr by 8,500 yr BP (Fig. S40D)^13^. This geographic isolation restricted habitats, promoted intensive inbreeding (*F*_ROH_ = 0.48 − 0.54; *F*_HBD_ = 0.48 − 0.53; Fig. 2C), and likely initiated allopatric divergence on the island (Fig. S41). Currently, DGD hosts ∼70 terrestrial vertebrate species^53^, and the surrounding waters form a strong genetic barrier for macaques.

CD and DGD are China’s earliest documented isolated macaque populations^44^. CD shows definitive genetic signatures of recent isolation, including abundant long ROH segments indicative of recent inbreeding (Fig. 2C) and minimal divergence from mainland *M. mulatta* (*F*st = 0.13; Fig. S7). Our species delimitation analyses did not consistently support CD as distinct (Table S4), which contrasts with the deeper divergence of DGD.

The issue of species delimitation has remained complex with much uncertainty^54^. Based on the Phylogenetic Species Concept (PSC)^54–57^, we propose that the DGD macaque represents a distinct species as evidenced by 1) monophyly in genome-scale phylogenies (Fig. 1C, S2) and Admixture’s complete ancestral differentiation at K = 2 (Fig. 1C); 2) 1.94 million population-specific variants (Fig. S17) affecting 251 genes with derived missense mutations (Table S6) and three mitochondrial nonsynonymous substitutions; and 3) divergence that was produced by >9,500 years of geographical isolation, corroborated by ABC coalescent modeling (915–1,291 generations), with no subsequent gene flow (Fig. S22–25). The extreme genetic distinctiveness (*F*st = 0.462; significantly exceeding recognized macaque species thresholds of 0.13–0.34, P < 0.001; Fig. S8) transcends preliminary morphological classifications. Swinhoe (1867) described this population from North Lena (now known as DGD) as *Macaca sancti-johannis*^58^. The holotype, skin and skull of a juvenile female, are stored in the Natural History Museum of the United Kingdom (NHMUK) in London^59^.

### Genomic consequences and drift-driven rapid speciation

Island populations of large vertebrates often face increased long-term extinction risks compared with their mainland counterparts^60^. These risks are further intensified on small islands when combined with anthropogenic disturbances and habitat degradation^61^. DGD macaque population exhibits remarkable resilience. Despite its limited area (13.2 km²), the island supports over 1,300 macaques, which are sustained by extensive primary forests and abundant freshwater resources^44,62^ that form a rare and valuable natural habitat.

This population endured a severe historical bottleneck, with demographic and coalescent modeling indicating an effective population size (Ne) of approximately 40 (SMC++/StairwayPlot2; Fig. 3C, S20–S21) and only 193–277 (ABC coalescent simulations; Table S11) founders surviving the initial isolation. These constraints intensified genetic drift and reduced the efficacy of purifying selection^63–65^. In such small populations, the efficacy of selection against weakly deleterious mutator alleles is reduced, leading to elevated germline mutation rates—a prediction consistent with the drift-barrier hypothesis that genetic drift impairs the optimization of DNA repair mechanisms^66^. In keeping with this, we observed that the zero-to four-fold heterozygosity ratio increasing from ∼1.2 to ∼1.4 (Fig. 3B) as did the proportion of deleterious variants (high + moderate impact) from ∼0.66 to ∼0.76 (Fig. S16). These effects also resulted in the equalized allele frequencies across loci (SFS; Fig. S27), The simultaneous dispersion to both sides of Tajima’s D (experiencing bottlenecks, elevating ancestral allele frequencies → positive Tajima’s D; followed by recent expansion, excess low-frequency alleles → negative Tajima’s D; Fig. S11). The fixation of 251 missense mutations, predominantly in immune-related genes (Tables S6–S7; Fig. S18), highlights potential functional adaptations shaped primarily by rapid evolutionary processes. Moreover, the pronounced accumulation of population-specific fixed variants underscores the dominant influence of genetic drift, in which accelerated fixation rates drove marked genetic divergence. These results empirically validate Kimura’s neutral theory predictions under severe bottlenecks^68^, establishing genetic drift as the architect of the accelerated divergence of DGD^69^.

Prior studies have shown that genetic drift can drive rapid speciation in isolated vertebrates. For example, the Norwegian lemming diverged from western Siberian populations shortly before the Last Glacial Maximum, without gene flow^6^. Similarly, the Saimaa ringed seal shows profound divergence (>60,000 years) in genetic, behavioral, and ecomorphological traits, including adaptive dentition and tongue morphology^70^. Demographic reconstructions confirm repeated bottlenecks and recent isolation as the primary forces, rather than selection, and underpin the formation of distinct Evolutionarily Significant Units (ESUs)^71^.

### Conservation strategy for the DGD macaque

The DGD macaque population indicates that drift-purging equilibrium can promote long-term survival^22,25,72^. Our results suggest that the DGD population, despite losing 65.8% of its ancestral genetic diversity through serial founder effects (Fig. S26A), achieved a decrease in LoF genetic load (Fig. 4A). This counterintuitive outcome results from two synergistic mechanisms: (1) recessive allele exposure through increased homozygosity (Fig. 4B), which eliminates lethal variants while reducing LoF load by 68% within homozygosity runs (Fig. S27), and (2) purifying selection against highly deleterious variants in key metabolic pathways that are required for survival under insular constraints (Fig. 5B). Concurrently, purifying selection acted against *de novo* mutations (singleton Ts/Tv = 0.25), depressing deleteriousness at homozygous loci. Therefore, genetic drift facilitated survival by unmasking recessive liabilities for elimination.

Functional genomic analyses reveal a triage mechanism governing the fate of LoF variants. On one hand, purifying selection eliminated LOF mutations in 1,042 genes associated with lipid/energy metabolism pathways (Fig. 5B), including critical enzymes, such as CPT1C and ACADL, which likely prevented lethal ketosis during postglacial resource scarcity^49,50^. This adaptive purging targeted mutations that were incompatible with thermoregulatory and starvation-response demands, particularly in fatty acid β-oxidation^49–51^. Conversely, genetic drift fixed 85 LoF genes affecting cell adhesion and metabolism (Fig. 5C), including seven ultra-deleterious variants, such as a splice-acceptor mutation in SKAP2 (Fig. S36). These low-frequency and relatively moderate deleterious mutations accumulated stochastically in the insular population, potentially compromising long-term fitness^22^. This dichotomy mirrors broader debates about LoF effects, which may persist neutrally or rarely confer adaptation^73^. Our multispecies ancestral state reconstruction confirmed 34 high-confidence-derived LoF fixations (Fig. S32–S38), which highlight the role of drift in shaping genomic “legacy costs.”

In conclusion, despite its extremely small population size, the current genetic status of the DGD insular population has defied conventional expectations of inbreeding vulnerability. Gene function predicts vulnerability, necessitating prioritized monitoring of fixed LoF alleles in essential pathways (e.g., *SKAP2*), whereas genetic rescue efforts must determine the complementarity of donor-recipient deleterious alleles to avoid reintroducing purged lethal mutations. The DGD case illustrates how isolation can promote resilience through genomic austerity: purging lethal recessives, while also accumulating tolerable deleterious legacies. Future studies should develop integrated models of deleterious variation landscapes and test for parallel purging-fixation patterns in analogous insular populations. Genetic rescue strategies must account not only for diversity loss, but also for the functional spectrum of segregating deleterious variation.

### Brief Material & Methods

We generated whole-genome sequencing data for 51 macaques, including 30 newly sequenced wild individuals and 21 previously collected samples for which we increased sequencing depth (Table S1). This dataset included 26 insular macaques (6 from CD and 20 from DGD) and 25 mainland representatives of four recognized *Macaca mulatta* subspecies: 5 *M. m. mulatta*, 8 *M. m. lasiotis*, 7 *M. m. littoralis*, and 5 *M. m. brevicaudus*. All samples were sequenced using Illumina NovaSeq 6000 (150-bp paired-end), achieving an average depth of 33.5×.

We defined two datasets for analysis. Dataset I (Fig. 1a; Table S1) comprised the 51 genomes above plus 61 published rhesus macaque genomes (5 *M. m. mulatta*, 23 *M. m. lasiotis*, 28 *M. m. littoralis*, 5 *M. m. tcheliensis*) to explore population structure and divergence between continental and insular groups. Dataset II expanded this panel by including additional macaque species and a *Papio* outgroup for interspecific analyses (e.g., gene flow and phylogenetics).

Variants were called using GATK v4.2 following best practices^74–75^, filtered stringently, and annotated with SnpEff v4.3^75^. Kinship was inferred using KING v2.1.3^76^. Population structure was assessed via PCA (VCF2PCACluster^77^), ADMIXTURE v1.3^78^, and phylogenetic trees built using TreeBest and RAxML v8.2.12^79^, visualized with iTOL v6^80^. Mitogenomes were assembled using NOVOPlasty v4.3^81^, aligned with MAFFT v7^82^, and analyzed in PopART v1.7^83^.

Haplotypes were phased with Beagle v5.4^84^ and SHAPEIT v1r6887^85^; local recombination rates were estimated using FastEPRR v2.0^86^. Haplotype-based structure was inferred using fineSTRUCTURE v4.1.1^87^, and effective migration surfaces estimated with EEMS^88^. Genetic diversity and linkage disequilibrium were assessed with ANGSD v0.938^89^, VCFtools v0.1.15^90^, and PopLDdecay v3.42^91^. Runs of homozygosity and inbreeding were evaluated using PLINK v2.0^92^ and RZooRoH v0.3^93^. Species delimitation analyses were performed using BPP v4.6.2^38^. Genetic load and selection efficacy were evaluated via derived allele frequencies, SnpEff annotations^75^, Rx/y ratio^94^, and GERP++ scores mapped from hg38 using LiftOver^95,96^. Stairway Plot v2^41^ and PSMC v0.6.5^42^ were used to infer population history; SMC++ v1.15.2^43^ further resolved recent demography. Demographic modeling was conducted with fastsimcoal2 v2.8^97^. Gene flow was assessed using TreeMix v1.13^98^, ADMIXtools^99^, and Dsuite^45^. Functional enrichment of candidate gene sets was performed using clusterProfiler v4.6.0^100^.

## Acknowledgements

The work was supported by the National Key Research and Development Projects of the Ministry of Science and Technology of China (2023YFC3304003), The Sino-German Mobility Programme (Grant No. M-0084), State Key Laboratory of Animal Biodiversity Conservation and Integrated Pest Management (Grant No. SKLA2504), the China Scholarship Council Innovative Talent Programme (No.2022-2260). Yunnan Fundamental Research Kunming Medical University Projects (202501AY070001-119) and High-level talent introduction project of Yunnan Provincial Health Commission(2024-KHRCBZ-A01) and We thank to Dietmar Zinner for his valuable comments on the article.

## Author Contributions

M.L., J.Q. and C.R. conceived and designed the analyses. J.Q., S.L., L.Z. and R.W. conducted analyses. J.Q., Z.Z., Y.T., M.Z., Y.S. and Y.Y. conducted the sampling. G.L., C.H., W.X., Q.Z. and D.R. provided help for data analysis. Z.L., X.Z., J.R. and C.R. provided revision comments. J.Q., M.L., J.R. and C.R. wrote and edited the manuscript.

## Data Availability Statement

The raw sequence data reported in this paper have been deposited in the Genome Sequence Archive in National Genomics Data Center, China National Center for Bioinformation / Beijing Institute of Genomics, Chinese Academy of Sciences (GSA: PRJCA027357) that are publicly accessible at https://ngdc.cncb.ac.cn/gsa.

## Supplementary Information

Figure S1-Figure S41; Table S1-Table S18

