## supplementary file for "Drift and isolation drive genomic erosion and island speciation in a lineage of macaques"

### Material & Methods

#### Sample collection and genome sequencing.

Blood samples were collected from 30 wild-born rhesus macaques from three populations: Daba, Hubei (n = 4), CD (n = 6), and DGD (n = 20) (Fig. 1A, Table S1). Whole blood (5 ml/individual) was drawn from the femoral vein into EDTA-anti-coagulant tubes and immediately stored at −80°C until DNA extraction. All procedures adhered to the Good Experimental Practices guidelines of the Institute of Zoology, Chinese Academy of Sciences under oversight by the Institutional Animal Care and Use Committee. These blood samples were supplemented with samples from 21 previously studied[1] continental macaques for enhanced sequencing depth. All 51 samples were subject to whole-genome sequencing at Annoroad Gene Technology (Beijing) using the Illumina NovaSeq 6,000 platform, which generated 150-bp paired-end reads with a mean depth of 33.49 × (Table S1). Raw sequencing data are available under accession PRJCA027357 at the Genome Sequence Archive (https://ngdc.cncb.ac.cn/gsa) of China’s National Genomics Data Center.

#### Dataset generation.

For comparative analysis, we integrated sequence data from 61 additional individuals representing continental Chinese macaque populations from our prior study (PRJNA345528)[1] to form Dataset I (112 individuals total). Dataset II was generated by adding 10 Indian-origin macaques (PRJNA251548), one individual from each of seven congeneric species (*M. cyclopis* [SRR14843533], *M. fuscata* [SRR14843532], *M. fascicularis* [SRR8194878], *M. thibetana* [CRR1146560], *M. leucogenys* [SRR14843520] *M. arctoides* [SRR14843525], *M. sylvanus* [SRR14843527]), and a baboon outgroup [SRR8285766] to Dataset I.

**Variant calling and filtering.**

The raw FASTQ reads obtained from the Illumina platform were end-trimmed using the default settings of Trim Galore for Illumina (https://github.com/FelixKrueger/TrimGalore). The processed reads were aligned to the rhesus macaque (*Macaca mulatta*) reference genome (assembly GCF_003339765.3) using the Burrows-Wheeler Aligner (v0.7.17 bwa mem)[2] with default parameters. The alignment files were processed using SAMtools v1.9[3] for sorting, followed by duplicate removal using Picard MarkDuplicates v2.20.2 (http://broadinstitute.github.io/picard/) to generate indexed BAM files. Variant calling was done for each sample using GATK v4.2[4] HaplotypeCaller to generate GVCFs, followed by joint genotyping using CombineGVCFs and GenotypeGVCFs. Insertions and deletions (InDels) were excluded from the downstream analysis. Genotypes exhibiting extreme coverage depths were masked. Autosomes required masking if coverage was below 0.5 × or above 2 × the sample average, whereas separate criteria were applied to the X chromosome. The Y chromosome was excluded entirely. Subsequent SNP filtering was done using GATK VariantFiltration with stringent thresholds (QD < 2.0; QUAL < 30.0; SOR > 3.0; FS > 60.0; MQ < 40.0;

MQRankSum < −12.5; ReadPosRankSum < −8.0). In addition, heterozygous genotypes with minor allele support below 0.25 were masked. Finally, repetitive regions within the reference genome were identified and masked using the snpable pipeline (k-mer size = 100 bp). Additional filters were used to remove SNPs with missingness ≥10%, quality < 20 (phred-scale), and nonbiallelic sites. The final datasets contained 71,127,020 autosomal SNPs in Dataset I and 130,486,137 in Dataset II (Table S2). All variants were annotated through SnpEff v4.3[5] using the Mmul_10 GFF3 annotation from the NCBI (Table S2).

**Inference of kinship in the population.**

To examine the kinship coefficient within a population, we estimated the relatedness of the samples in each population using KING v2.1.3[6] with default parameters. An estimated kinship coefficient range >0.354, [0.177, 0.354], [0.0884, 0.177], and [0.0442, 0.0884] corresponds to duplicate sample/monozygotic twins with $1^{st}$-degree, 2nd-degree, and 3rd-degree relationships, respectively. We excluded the C_rhe_54 sample from the following population structure analysis as it had a close relationship with C_rhe_51 (kinship coefficient of 0.4348) (Table S17).

**Phylogeny, population structure, and PCA.**

Phylogenetic trees based on autosomal SNPs were calculated with NJ (for Dataset I) and ML (for both Datasets) algorithms using *Papio anubis* as the outgroup. The NJ tree was generated with TreeBest software (http://treesoft.sourceforge.net/treebest.shtml) with 1,000 bootstrap replicates. The ML tree was reconstructed in RAxML v8[7] using the GTR + CAT substitution model and 1000 bootstrap replicates. iTOL v6[8] was used to visualize the phylogenetic trees. The PCA was conducted using VCF2PCACluster v1.41[9]. Admixture v1.3[10] was used to examine individual ancestry proportions, with coancestry clusters ranging from k = 2 to k = 9. The analysis was run 40 times for each k. The best run was selected based on the likelihood of each run. The best value of the coancestry cluster (k) was estimated using a cross-validation procedure in the Admixture software.

**Phylogeny of mitogenomes.**

NOVOPlasty v4.3[11] was used to *de novo* assemble the mitochondrial genomes (mitogenomes) of all samples. They were annotated with MitoZ v2.4[12]. Cox1 and Cytb were extracted and aligned using MAFFT v7[13]. Indels and poorly aligned positions were removed by Gblocks v0.91b[14]. A haplotype network was generated using the Median-Joining method in PopART v1.72b[15]. An ML tree was reconstructed using RAxML v8 with a GTRGAMMA substitution model and 1,000 bootstrap replicates.

**Phasing genomic VCF file.**

Dataset I contains all macaque VCFs for the high-quality genomic SNPs, which were phased using Beagle v1399[16] with default settings. For each sample, phase informative reads (PIRs) were extracted from the BAM file of the corresponding genome sequences using the extractPIRs tool (implemented in the SHAPEIT package) with parameters "--base-quality 20 --read-quality 30." By combining VCF files that had been processed by Beagle and PIRs files, these VCF files were rephased, and high-quality haplotypes were constructed across genomes using SHAPEIT v1r68[17].

This VCF was split into two files, continental and island, and two separate recombination maps were calculated, along with the distribution of windowed population recombination rate (ρ) for the mainland and island macaque population, which was estimated using FastEPRR v2.0[18] with a 100-Kb sliding window and a 50-Kb step (Fig. S39).

**fineSTRUCTURE.**

The VCF file was reformatted separately using the impute2chromopainter.pl script for each chromosome-level scaffold. Recombination rate maps were generated using the makeuniformrecfile.pl script, assuming a constant rate of recombination per base. The fineSTRUCTURE program[19] was run in automatic linked mode, using the default settings. The results were plotted using the provided R functions scripts.

**Genetic Differentiation and Effective Migration Surfaces (EEMS) Analysis.**

Genome-wide genetic differentiation was quantified using Weir and Cockerham's $F$st. Pairwise $F$st values were calculated using VCFtools v0.1.15[20] with 50 kb sliding windows advanced in 25 kb steps. This generated a genome-wide pairwise $F$st matrix for all study groups that incorporated both differentiation classifications and population structure inferences. To test for deviations from isolation-by-distance expectations, migration and diversity patterns were modeled using the Effective Migration Surfaces (EEMS) framework[21]. Geographic ranges were defined according to the macaque distribution data. Genetic dissimilarity matrices were calculated using the EEMS' bed2diffs_v1 program. For each analysis, 9 million MCMC iterations were executed with a thinning interval of 9,999. Convergence was assessed three times through parallel, independent runs. The results were visualized in R using the reemsplots2 package, with Figure 1B integrating population sampling locations.

**Coalescent-based species delimitation.**

Coalescent-based species delimitation was assessed using Bayesian Phylogenetics and Phylogeography (BPP v4.6.2)[22], which uses the multispecies coalescent model to compare alternative species-delimitation hypotheses in a Bayesian framework, while accounting for incomplete lineage sorting resulting from ancestral polymorphisms and gene tree-species tree discordance. A10 analyses (species delimitation using a user-specified guide tree) were used with the species tree (mulatta, DGD) for DGD and (mulatta, CD) for CD as the guide tree. Because of the computational intensity, BPP analyses were performed on three datasets: 1) 1,000 nuclear loci of 1,000 bp each, 2) 500 randomly selected gene loci, and 3) the complete mitogenomes treated as a single locus. Genotypes were converted to sequences using the Bcftools consensus module, which represents heterozygous sites with IUPAC ambiguity codes. The 1,000 nuclear loci were selected by these filtering criteria to satisfy BPP assumptions: 1) locus length between 500–1,000 nucleotides to ensure no intra-locus recombination, 2) no missing data per locus, and 3) each locus on a separate scaffold to ensure sufficient physical distance for free recombination between loci. Mitochondrial loci were treated as a single locus following BPP recommendations, as mitochondrial genomes generally do not undergo recombination. The JC69 substitution model was used, as it is adequate for closely related species, and equal probabilities were used for the prior species model. The analyses were run with both algorithms 0 and 1 using different values of ε, α, and m, respectively (Table S4). For each species tree model, inverse-gamma priors: θ ~ IG(3, 0.003) were assigned for all θs (quantifying within-species genetic diversity) and τ ~ IG(3, 0.0004) for the root age τ0. Diffuse priors (a = 3) were used, with b adjusted for reasonable means. The θ prior (mean heterozygosity 0.0015/kb) was based on the calculated genetic diversity in our samples. The τ prior (mean sequence divergence 0.0002/kb) was based on MEGA7 (Table S18). Each analysis consisted of 200,000 MCMC iterations after a 10,000-iteration burn-in, with sampling every two iterations (yielding 100,000 samples). Convergence was assessed by comparing the results between runs.

**Genetic diversity and Inbreeding.**

Individual heterozygosity was estimated using ANGSD v0.938[23] with the quality filtering parameters "-minQ 20 -minmapq 30" to exclude low-quality bases and reads. Nucleotide diversity (π) was calculated in 50 kb sliding windows advanced in 25 kb steps using VCFtools v0.1.15[20]. The comparative primate genetic diversity statistics were obtained from published data[24]. Inter-group diversity comparisons were done using Welch's two-sample t-tests. Linkage disequilibrium coefficients ($r^2$) were calculated with PopLDdecay v3.42[25]. Tajima's D was assessed using VCFtools v0.1.15 with the same slide window parameters. SFS with ANGSD (-minMapQ 30 -minQ 20 -doCounts 1 -GL 1 -doSaf1 -maxIter 100) were determined.

IBD segments were detected using two complementary approaches: genome-wide ROH were identified using PLINK v2.0[26] with sliding-window parameters requiring ≥1 SNP per 50 kb (--homozyg --homozyg-kb 100 --homozyg-snp 50 --

homozyg-density 50 --homozyg-window-snp 50 --homozyg-window-het 1 --homozyg-window-missing 5), where segments were considered broken upon encountering >1 heterozygous or >5 missing calls per window. Concurrently, RZooRoH v0.3[27] was used to characterize individual inbreeding histories through Hidden Markov Models (HMMs)[28], which estimate ancestral inbreeding contributions across generational timescales and classify HBD segments into distinct age cohorts that correspond to approximately 4, 16, 64, 256, 1,024, 4,096, 16,384, and 65,536 previous generations, plus a non-HBD category. To quantify autozygosity, three related inbreeding coefficients were calculated: the true genomic IBD fraction ($F_{\mathrm{IBD}}$), its empirical estimate from ROH ($F_{\mathrm{ROH}}$ = [observed homozygous SNPs - expected homozygous SNPs]/[total called SNPs - expected homozygous SNPs]), and the model-based HBD estimate ($F_{\mathrm{HBD}}$ = total autozygosity/callable genome length).

**Ratio of heterozygosity zero- to four-fold degenerate sites and high + moderate to low-impact variants.**

To evaluate selection efficacy directly, two ratios were calculated: (1) heterozygosity at zero- versus four-fold degenerate sites within the coding transcripts, in which mutations would always or never change the encoded amino acid, respectively; and (2) high/moderate-impact versus low-impact variants. Both ratios are expected to be increased in small populations because of the elevated frequency of deleterious alleles under strong genetic drift. These measures were derived from heterozygous loci to mitigate reference genome bias, while accounting for ancestral/derived allele status. Variant classification used SnpEff annotations, with high/moderate-impact variants defined as exon loss, 3' UTR truncation + exon loss, 5' UTR truncation + exon loss, coding sequence disruption, regulatory region ablation, missense variants, splice donor/acceptor/region alterations, stop-gained, frameshifts, or indels (insertions/deletions), whereas low-impact variants included synonymous, 5' UTR, premature start gain, initiator codon, start loss, and transcript retention events.

**Demographic reconstruction.**

Long-term demographic history was determined using the PSMC v0.6.5[29] model, which was applied to high-depth sequencing data and implemented three parameter configurations (-N25 -t15 -r5 -p “4+25*2+4+6,” -N25 -t15 -r5 -p “2+2+25*2+4+6,” and -N25 -t15 -r5 -p “1+1+1+1+25*2+4+6”) with a 11-year generation time [24,30] to mitigate false-positive signals[31]. Mutation rates ($\mu = 2.46 \times 10^{-8}$/bp/generation; 1.758 $\times 10^{-9}$/bp/years) were estimated in r8s v1.81[32] using 4-fold degenerate sites and calibrated with three soft-bound fossil constraints: *Papio anubis*-*Macaca sylvanus* (7.07 ± 0.37 Mya), *M. sylvanus*-*M. arctoides* (3.69 ± 0.65 Mya), and *M. arctoides*-*M. fascicularis* (3.62 ± 0.56 Mya)[33,34]. For recent demographic inference, SMC++ v1.15.2[35,36] was applied with parameters -time-points 1e2,1e6 -spline cubic -knots 35 -regularization-penalty 4 across all chromosomes. Paleoclimatic context was integrated using million-year-resolution atmospheric and sea-level records (NOAA Paleoclimatology Dataset ID: noaarecon-11932). Population bottleneck validation was done using a StairwayPlot v2[37] analysis of folded SFS derived from 20 DGD individuals processed through ANGSD described above.

**Simulation of evolutionary scenarios.**

ABC coalescent simulations in fastsimcoal2 v2.8[38] were used to infer speciation scenarios for the DGD macaque, assess potential secondary contacts, and estimate divergence times from other populations. To mitigate ancestral allele state bias, folded multidimensional site frequency spectra (multi-SFS) were generated using easySFS (https://github.com/isaacovercast/easySFS#easysfs) from whole-genome data. Our three-phase analytical framework consisted of: (1) an initial evaluation of 10 divergence models for the DGD island lineage, including scenarios with a ghost population diverging directly from mainland *M. m. littoralis*; (2) secondary contact analysis incorporating gene flow events informed by *D*-statistics, *f*-branch, *f3*-statistics, and TreeMix, which tested postdivergence migration along southern Chinese coasts; and (3) demographic integration of SMC++ inferred population bottlenecks to determine land-sea isolation effects, specifically distinguishing prebottleneck mainland mixing from postisolation dispersal mechanisms. For each model, 100 independent fastsimcoal2 runs were executed with 500,000 coalescent simulations per likelihood estimation (-n500,000), 60 conditional maximization cycles (-L60), and a minimum 100-count SFS bins (-C100), to select optimal parameters through maximum likelihood optimization and compare models via Akaike information criterion. Parameter uncertainty was quantified through non-parametric equipartition by partitioning the genome into 50 contiguous blocks of equal length, re-estimating SFS per block using easySFS, and performing 100 fastsimcoal2 optimizations per block using whole-genome maximum likelihood parameters as initial values. The final parameter distributions were derived from the block-specific maximum likelihood estimates, with standard errors computed across all 50 replicates.

**Gene flow analysis.**

To determine whether island populations experienced secondary contact with their mainland counterparts, we conducted integrated gene flow analysis using TreeMix v1.13[39], *D*-statistics[40], *f*-branch tests[41], and outgroup *f3*-statistics[40]. For TreeMix reconstruction, we generated maximum-likelihood trees with 1,000-SNP blocks (-k 1,000) using baboon as the outgroup root, and iteratively adding migration events (-m 1–10) over 20 replicates to ensure convergence. The resulting topologies were visualized using TreeMix's built-in R functions, with optimal migration edges determined using the OptM R package[42]. The Evanno method and Δm metric were used until 99.8% of inter-population variance was explained. *D*-statistics and *f3*-statistics were implemented in ADMIXtools v5.1[40] to evaluate scenarios: *D*(DGD, CD; X, *P. anubis*) for admixed populations (X), with significant gene flow indicated by positive D-statistics and |Z-scores| ≥ 3. Outgroup-*f3* analysis consisted of two configurations: *f3*(X; DGD, *P. anubis*) assessing gene flow from DGD to X populations (out island), and *f3*(DGD; X, *P. anubis*) testing introgression into DGD (in island), where $f3 < 0$ with $Z < -2$ indicated a significant admixture. Finally, we performed *f*-branch tests in Dsuite[41] to localize gene flow to specific internal branches, and visualized the outputs through dtools.py scripts.

**Genetic Load and the Purging of Deleterious Mutations**.

To compare patterns of deleterious burden, the genetic load was first estimated using the Rx/y ratio[43], which indicates the relative frequency of LoF and missense mutations in one population relative to another. Using the results of SnpEff annotation, "LoF" variants (defined as transcript ablation, splice donor, splice acceptor, stop gained, frameshift, insertion, deletion, or splice region variants), "Missense" variants, and "Synonymous" variants for Rx/y ratio (or called $R_{A,B}$) comparisons were extracted. The relative number of derived alleles for each comparison was calculated as follows:

$$R_{A,B}(\mathrm{C}) = \frac{L_{A,B}(C)}{L_{B,A}(C)} ,$$

Where $L_{A,B}$ (C) was determined by the following formula:

$$L_{A,B}(\mathrm{C}) = \frac{\sum_{i\in C} f_i^A(1-f_i^B)}{\sum_{i\in I} f_j^A(1-f_j^B)} .$$

At each site i, when $L_{A,B}(\mathrm{C})$ was estimated, the derived allele frequency in population A is defined as $f_i^A = d_i^A/n_i^A$, where $d_i^A$ is the number of derived alleles identified in population A and $n_i^A$ is the total number of alleles. The $f_i^B$ is similarly defined in population B. For the above formula, C and I indicate the category of protein-coding sites (LOF or missense) and a set of intergenic sites, respectively. To estimate the variance of $R_{A,B}(\mathrm{C})$, a 100-block jackknife was performed at the sites of the C set.

To determine whether the homozygous tract LOF rate was decreased in each group, the number of homozygous and non-homozygous LOF sites present in at least one individual was counted. The Kolmogorov-Smirnov test was used to assess the significance of the difference between relative rates of LOF variants in the homozygous and non-homozygous sites in the genome.

Based on comparisons to seven macaque species, ancestral/derived alleles were inferred at each locus. The 100 vertebrate species multiZ alignment (including Mmul_10) was downloaded from UCSC (http://hgdownload.soe.ucsc.edu/goldenPath/hg38/multiz100way/maf/), and GERP scores were transferred from hg38 to Mmul_10 using LiftOver[44,45]. Individual masked load was estimated as the sum of the GERP scores for deleterious derived alleles in heterozygous genotypes divided by called genotypes. The realized load was the sum of the GERP scores for homozygous deleterious derived alleles divided by all called sites. A GERP score threshold of > 4 indicated potentially deleterious mutations[46,47], as these loci are highly conserved (indicating strong selection constraints), whereas scores < 4 indicated neutral loci (more evolutionarily flexible, representing ~92% of the genome).

Purging recessive deleterious variants (i.e., LoF alleles) is expected to produce distinct signatures in homozygous tracts (runs of homozygosity; ROH) versus non-homozygous tracts[48]. Recessive LoF variants affecting viability or early survival should be less frequent in homozygous tracts, where they are exposed to purifying selection, compared with other genomic regions. To test this, we quantified LoF variant sites in homozygous and heterozygous genome portions. LoF rates were normalized using synonymous homozygous variant rates from corresponding regions (SnpEff output) to control for inter-individual homozygosity differences. Significant differences in relative LoF rates between homozygous and non-homozygous genome portions were determined using a paired *t*-test in R.

Finally, we constructed derived allele frequency spectra (DAF) for deleterious variants. To characterize purifying selection strength across populations, SFS were generated for distinct deleterious allele classes using SnpEff functional annotations: “LoF” (high-impact), “missense” (moderate-impact), and “synonymous” (low-impact) variants. Transition/transversion (ts/tv) ratios were calculated using bcftools v1.7[49] stats on singleton-containing VCF files to assess purifying selection against *de novo* mutations.

**Characterizing fixed and purged LoF standing variation in DGD.**

To evaluate the functional effect of LoF standing variation in the DGD population, two distinct gene sets were defined based on contrasting evolutionary trajectories:

Geneset I: Ancestral Standing LoF Variants Purged in DGD

This set consists of LoF variants present as standing variations in ancestral populations (*M. m. littoralis*, *M. m. brevicaudus*, CD) at moderate allele frequencies ($0.01 \leq MAF \leq 0.30$), but eliminated in DGD. These sites were never observed in the homozygous state in ancestral populations. We propose that these mutations caused recessive embryonic or early postnatal lethality when made homozygous during DGD’s period of extreme inbreeding. Their elimination likely resulted from intense purifying selection during the population bottleneck, in which deleterious alleles were exposed and removed through reduced fitness of homozygous carriers.

Geneset II: Ancestral Standing LoF Variants Fixed in DGD

This set contained ancestral LoF polymorphisms that became fixed in DGD through prolonged genetic drift. Although segregating neutrally or under weak selection in ancestral populations, these variants avoided purifying selection during demographic contraction of DGD. Their fixation represents a substantial genetic load that may compromise long-term adaptive potential by reducing population fitness by accumulating deleterious alleles[48,50,51].

Using the Integrative Genomics Viewer (IGV v2.7)[52], BAM files were visualized from continental and island populations and manually validated LoF variants, and functional enrichment analysis was performed on both gene sets containing fixed versus purged LoF variants using clusterProfiler v4.6.0 (R package)[53].

### Supplementary Figures

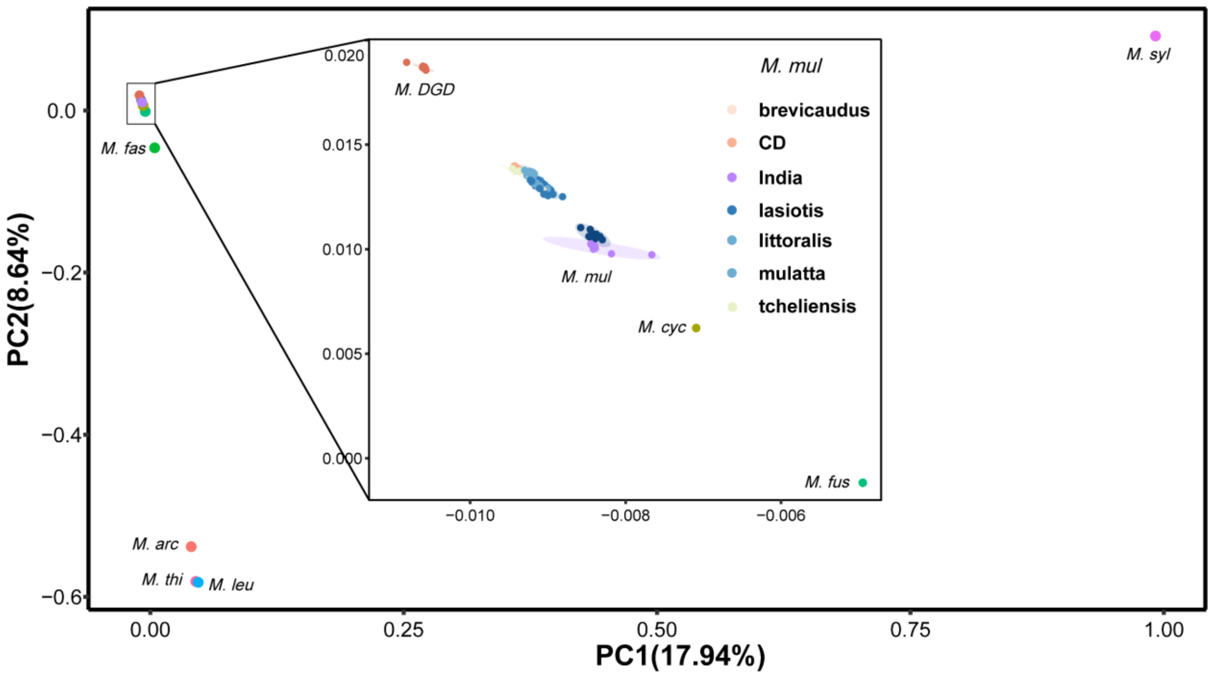


**Supplementary Figure 1.** PCA analysis of macaques. *M. mulatta* is enlarged inside the figure. Different subspecies and populations are represented by different colors. *M. mul* is *Macaca mulatta*; *M. syl* is *Macaca sylvanus*; *M. leu* is *Macaca leucogenys*; *M. thi* is *Macaca thibetana*; *M. arc* is *Macaca arctoides*; *M. fas* is *Macaca fascicularis*; *M. cyc* is *Macaca cyclopis*; *M. fus* is *Macaca fuscata*;

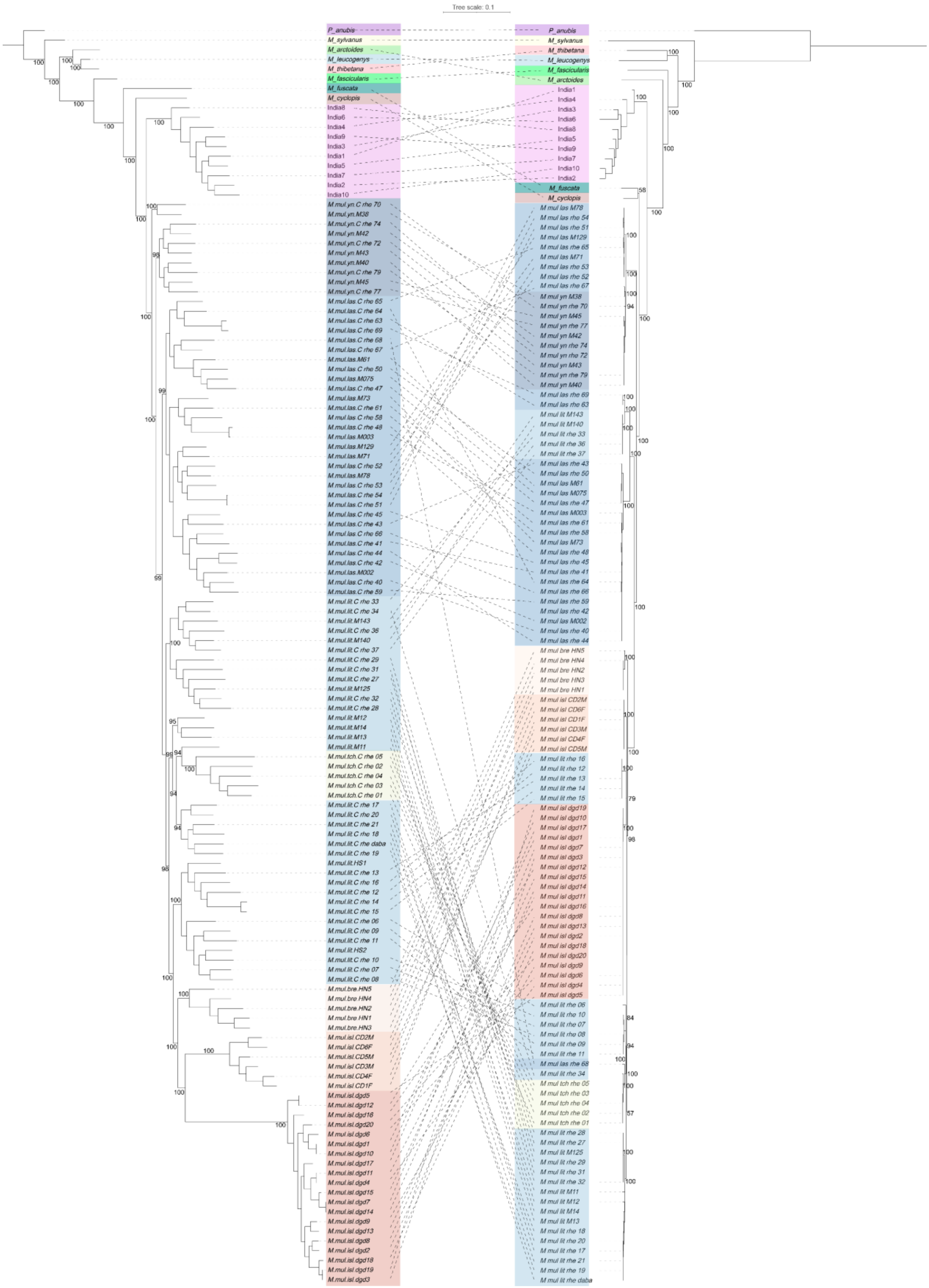
Tree scale: 0.1
P_anubis
M_sylvanus
M_arctoides
M_leucogenys
M_thibetana
M_fascicularis
M_fuscata
M_cyclopis
India8
India6
India4
India9
India3
India1
India5
India7
India2
India10
M.mul.yn.C rhe 70
M.mul.yn.M38
M.mul.yn.C rhe 74
M.mul.yn.M42
M.mul.yn.C rhe 72
M.mul.yn.M43
M.mul.yn.M40
M.mul.yn.C rhe 79
M.mul.yn.M45
M.mul.yn.C rhe 77
M.mul.las.C rhe 65
M.mul.las.C rhe 64
M.mul.las.C rhe 63
M.mul.las.C rhe 69
M.mul.las.C rhe 68
M.mul.las.C rhe 67
M.mul.las.M61
M.mul.las.C rhe 50
M.mul.las.M075
M.mul.las.C rhe 47
M.mul.las.M73
M.mul.las.C rhe 61
M.mul.las.C rhe 58
M.mul.las.C rhe 48
M.mul.las.M003
M.mul.las.M129
M.mul.las.M71
M.mul.las.C rhe 52
M.mul.las.M78
M.mul.las.C rhe 53
M.mul.las.C rhe 54
M.mul.las.C rhe 51
M.mul.las.C rhe 45
M.mul.las.C rhe 43
M.mul.las.C rhe 66
M.mul.las.C rhe 41
M.mul.las.C rhe 44
M.mul.las.C rhe 42
M.mul.las.M002
M.mul.las.C rhe 40
M.mul.las.C rhe 59
M.mul.lit.C rhe 33
M.mul.lit.C rhe 34
M.mul.lit.M143
M.mul.lit.C rhe 36
M.mul.lit.M140
M.mul.lit.C rhe 37
M.mul.lit.C rhe 29
M.mul.lit.C rhe 31
M.mul.lit.C rhe 27
M.mul.lit.M125
M.mul.lit.C rhe 32
M.mul.lit.C rhe 28
M.mul.lit.M12
M.mul.lit.M14
M.mul.lit.M13
M.mul.lit.M11
M.mul.tch.C rhe 05
M.mul.tch.C rhe 02
M.mul.tch.C rhe 04
M.mul.tch.C rhe 03
M.mul.tch.C rhe 01
M.mul.lit.C rhe 17
M.mul.lit.C rhe 20
M.mul.lit.C rhe 21
M.mul.lit.C rhe 18
M.mul.lit.C rhe daba
M.mul.lit.C rhe 19
M.mul.lit.HS1
M.mul.lit.C rhe 13
M.mul.lit.C rhe 16
M.mul.lit.C rhe 12
M.mul.lit.C rhe 14
M.mul.lit.C rhe 15
M.mul.lit.C rhe 06
M.mul.lit.C rhe 09
M.mul.lit.C rhe 11
M.mul.lit.HS2
M.mul.lit.C rhe 10
M.mul.lit.C rhe 07
M.mul.lit.C rhe 08
M.mul.bre.HN5
M.mul.bre.HN4
M.mul.bre.HN2
M.mul.bre.HN1
M.mul.bre.HN3
M.mul.isl.CD2M
M.mul.isl.CD6F
M.mul.isl.CD5M
M.mul.isl.CD3M
M.mul.isl.CD4F
M.mul.isl.CD1F
M.mul.isl.dgd5
M.mul.isl.dgd12
M.mul.isl.dgd16
M.mul.isl.dgd20
M.mul.isl.dgd6
M.mul.isl.dgd1
M.mul.isl.dgd10
M.mul.isl.dgd17
M.mul.isl.dgd11
M.mul.isl.dgd4
M.mul.isl.dgd15
M.mul.isl.dgd7
M.mul.isl.dgd14
M.mul.isl.dgd9
M.mul.isl.dgd13
M.mul.isl.dgd8
M.mul.isl.dgd2
M.mul.isl.dgd18
M.mul.isl.dgd19
M.mul.isl.dgd3
P_anubis
M_sylvanus
M_thibetana
M_leucogenys
M_fascicularis
M_arctoides
India1
India4
India3
India6
India8
India5
India9
India7
India10
India2
M_fuscata
M_cyclopis
M mul las M78
M mul las rhe 54
M mul las rhe 51
M mul las M129
M mul las rhe 65
M mul las M71
M mul las rhe 53
M mul las rhe 52
M mul las rhe 67
M mul yn M38
M mul yn rhe 70
M mul yn M45
M mul yn rhe 77
M mul yn M42
M mul yn rhe 74
M mul yn rhe 72
M mul yn M43
M mul yn rhe 79
M mul yn M40
M mul las rhe 69
M mul las rhe 63
M mul lit M143
M mul lit M140
M mul lit rhe 33
M mul lit rhe 36
M mul lit rhe 37
M mul las rhe 43
M mul las rhe 50
M mul las M61
M mul las M075
M mul las rhe 47
M mul las M003
M mul las rhe 61
M mul las rhe 58
M mul las M73
M mul las rhe 48
M mul las rhe 45
M mul las rhe 41
M mul las rhe 64
M mul las rhe 66
M mul las rhe 59
M mul las rhe 42
M mul las M002
M mul las rhe 40
M mul las rhe 44
M mul bre HN5
M mul bre HN4
M mul bre HN2
M mul bre HN3
M mul bre HN1
M mul isl CD2M
M mul isl CD6F
M mul isl CD1F
M mul isl CD3M
M mul isl CD4F
M mul isl CD5M
M mul lit rhe 16
M mul lit rhe 12
M mul lit rhe 13
M mul lit rhe 14
M mul lit rhe 15
M mul isl dgd19
M mul isl dgd10
M mul isl dgd17
M mul isl dgd1
M mul isl dgd7
M mul isl dgd3
M mul isl dgd12
M mul isl dgd15
M mul isl dgd14
M mul isl dgd11
M mul isl dgd16
M mul isl dgd8
M mul isl dgd13
M mul isl dgd2
M mul isl dgd18
M mul isl dgd20
M mul isl dgd9
M mul isl dgd6
M mul isl dgd4
M mul isl dgd5
M mul lit rhe 06
M mul lit rhe 10
M mul lit rhe 07
M mul lit rhe 08
M mul lit rhe 09
M mul lit rhe 11
M mul las rhe 68
M mul lit rhe 34
M mul tch rhe 05
M mul tch rhe 03
M mul tch rhe 04
M mul tch rhe 02
M mul tch rhe 01
M mul lit rhe 28
M mul lit rhe 27
M mul lit M125
M mul lit rhe 29
M mul lit rhe 31
M mul lit rhe 32
M mul lit M11
M mul lit M12
M mul lit M14
M mul lit M13
M mul lit rhe 18
M mul lit rhe 20
M mul lit rhe 17
M mul lit rhe 21
M mul lit rhe 19
M mul lit rhe daba

**Supplementary Figure 2.** ML phylogenetic tree of macaques based on autosomal SNP (left) and mtDNA (right). Tree has been rooted with outgroup *Papio anubis*. Bootstrap values of the main nodes are shown at the branch.

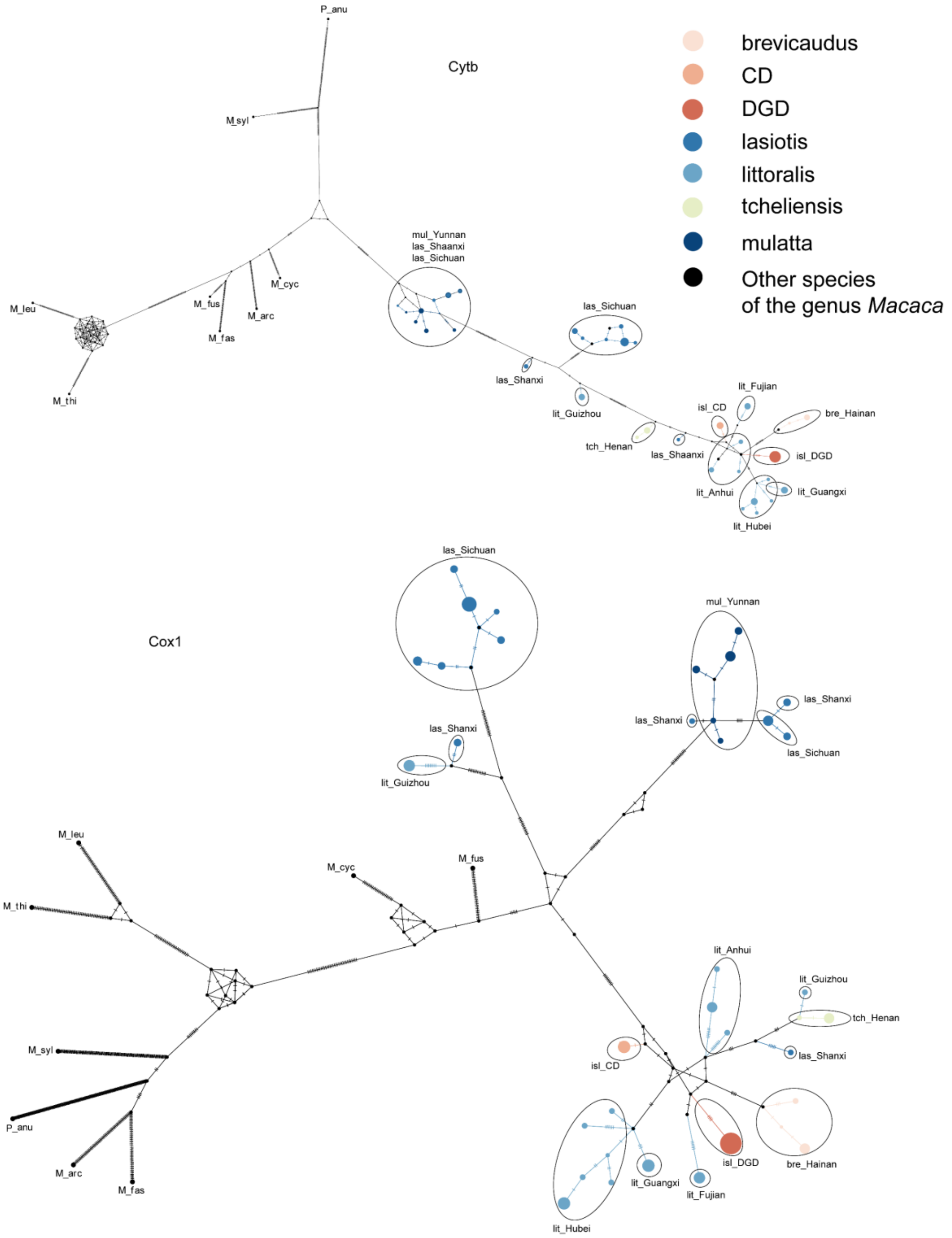

**Supplementary Figure 3.** Haplotype network analyses based on extracted mitochondrial Cytb (top) and Cox1 (bottom) sequences. The Cytb network suggests that DGD derived from *M. m. littoralis* from the southeastern mainland of China. *M. syl* is *Macaca sylvanus*; *M. leu* is *Macaca leucogenys*; *M. thi* is *Macaca thibetana*; *M. arc* is *Macaca arctoides*; *M. fas* is *Macaca fascicularis*; *M. cyc* is *Macaca cyclopis*; *M. fus* is *Macaca fuscata*;

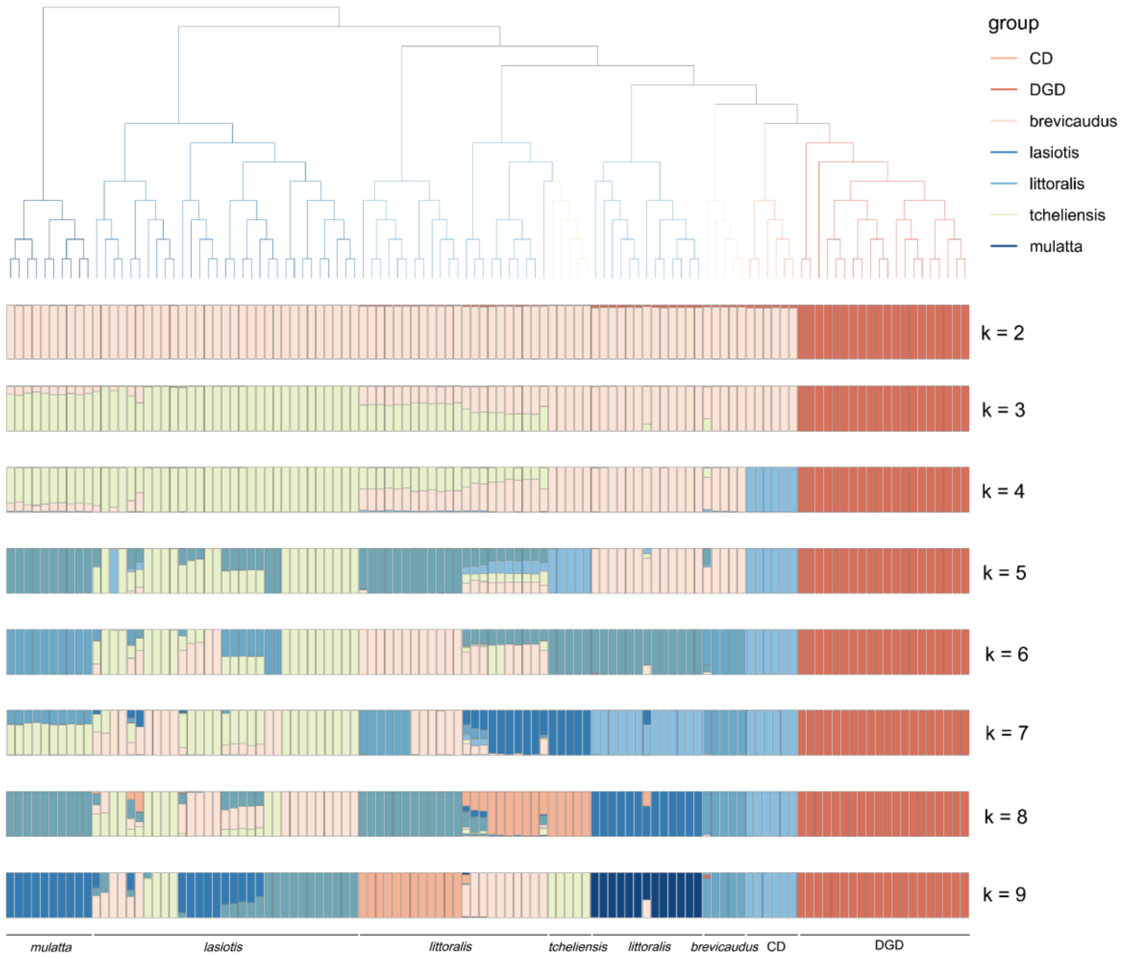


**Supplementary Figure 4.** ADMIXTURE analyses and phylogenetic ML tree reconstruction for dataset I with 112 individuals. At K=2, DGD forms a separate ancestral component (red) from all other *M. mulatta* populations (pink).

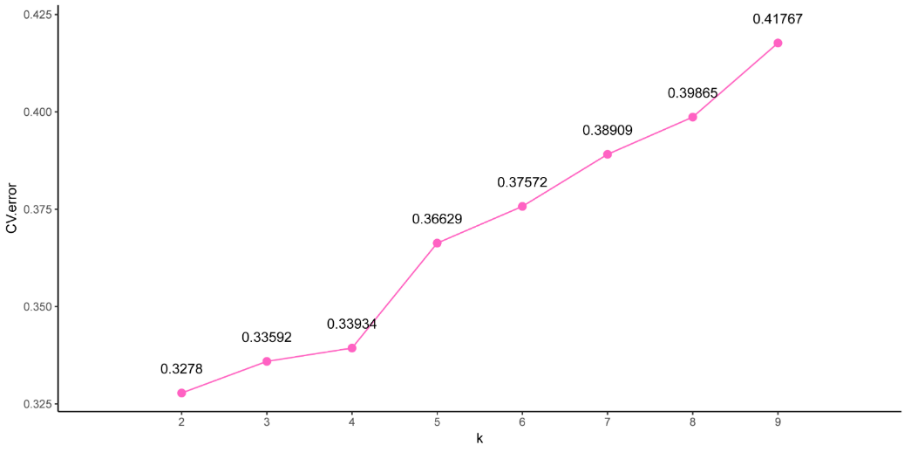


**Supplementary Figure 5.** Cross-validation (CV) error for different number of K in the ADMIXTURE analysis. Minimum of estimated CV error on K= 2 suggests the most suitable number of ancestral populations (see also Fig. S4).

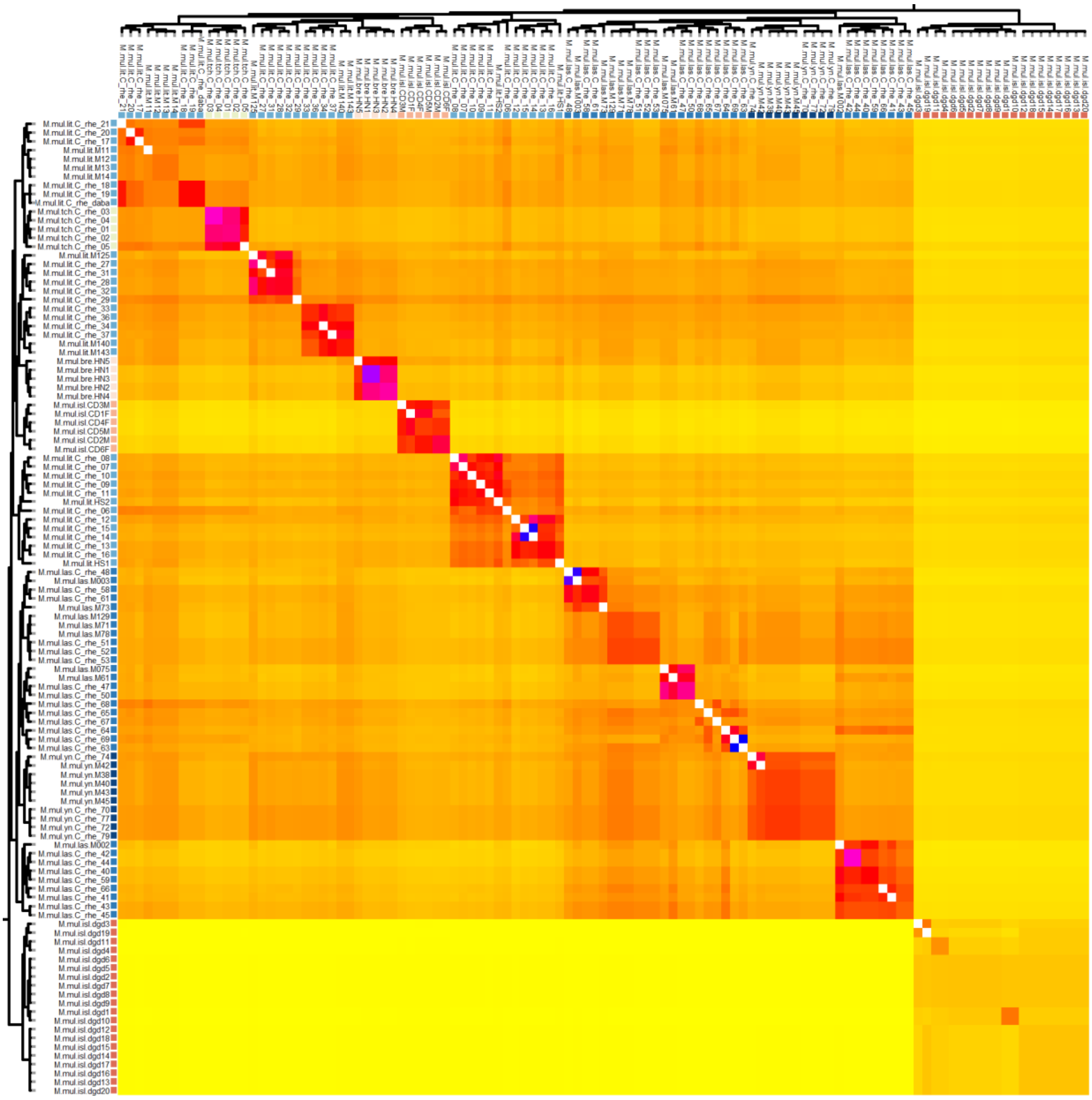

**Supplementary Figure 6.** fineSTRUCTURE coancestry matrix presented as a heatmap where cell color represents estimated shared genetic ancestry from high (blue, red) to low (yellow), run on whole genome SNP data for Dataset I. Sample color coded by population origin.

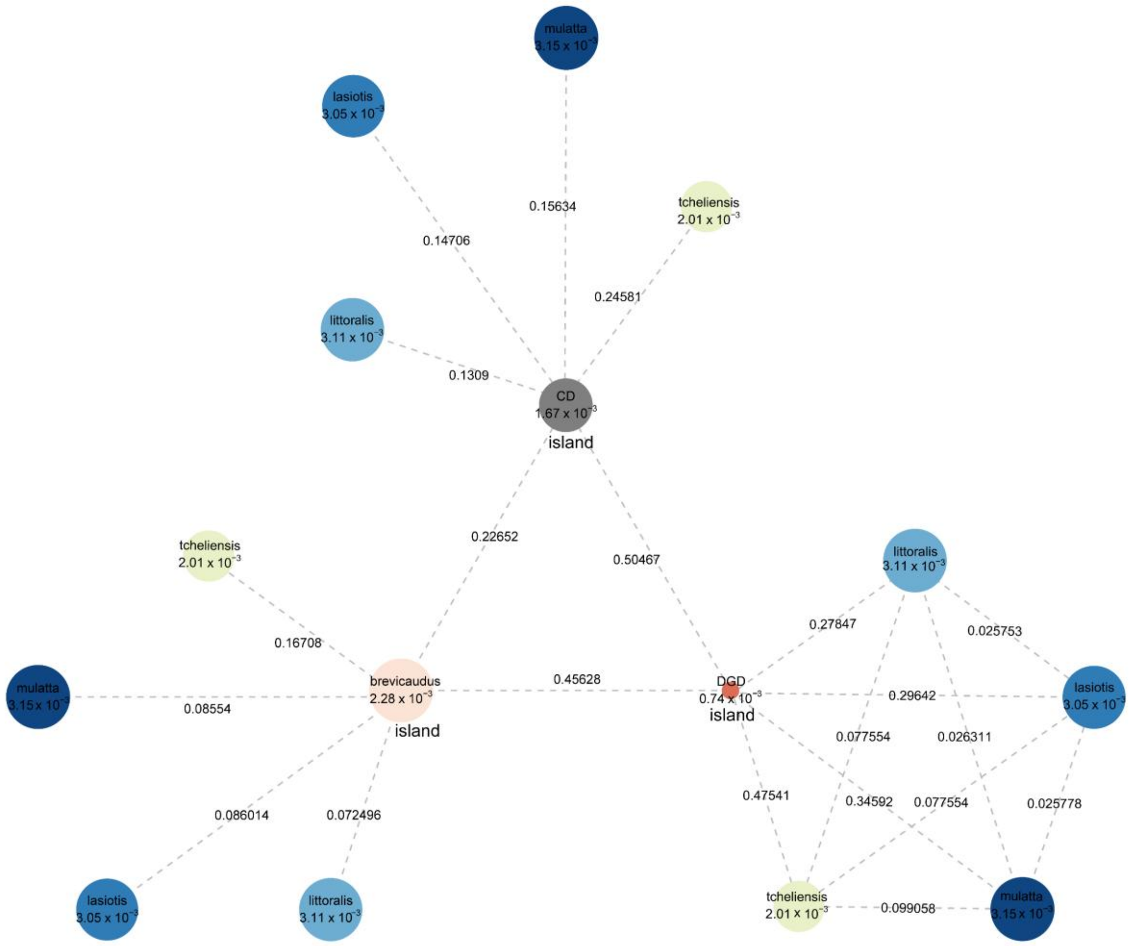


**Supplementary Figure 7.** Visualization of gene-wise fixation index (*F*st) between the island subspecies *M. m. brevicaudus,* DGD and CD versus mainland *M. mulatta* subspecies, with population-specific nucleotide diversity ($\theta\pi$) shown concurrently. Color gradients correspond to distinct taxonomic units (subspecies) and island-associated groups. Circle diameters scale proportionally with genetic diversity levels. Numeric *F*st estimates along interpopulation connectors quantify pairwise differentiation magnitudes.

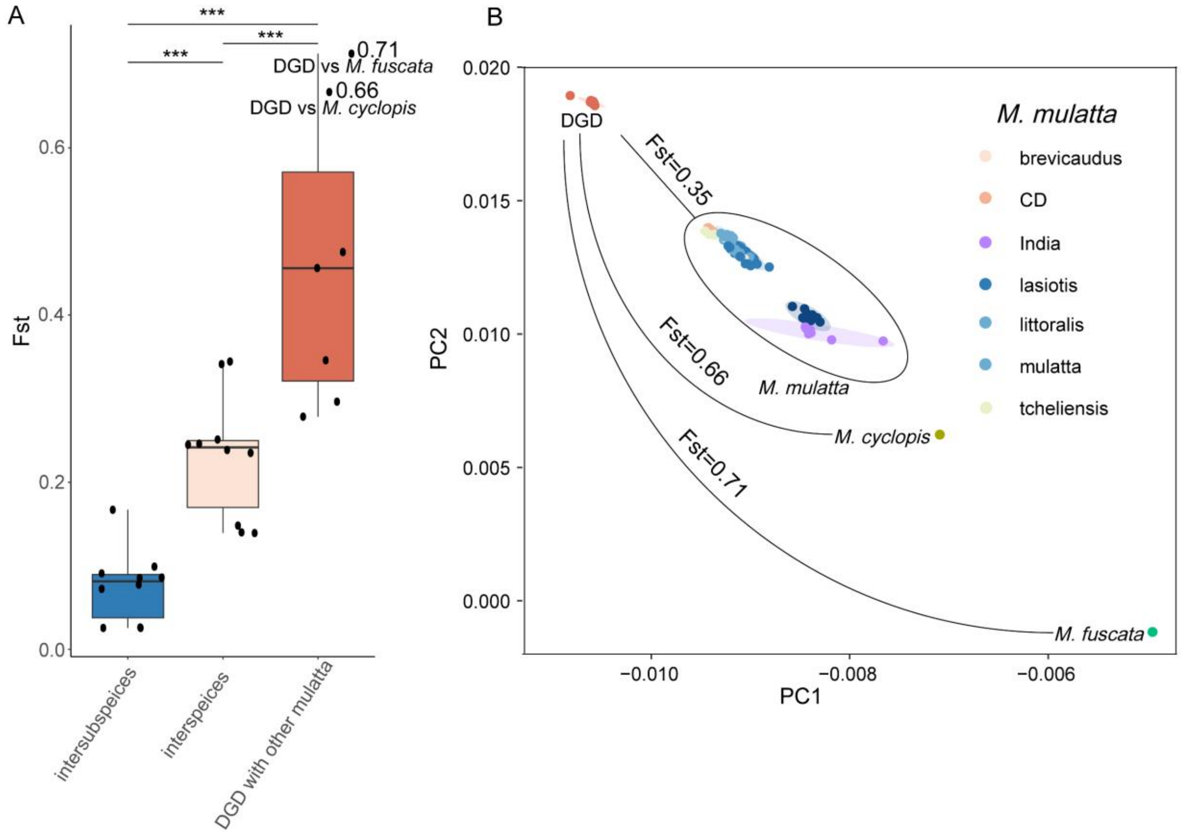


**Supplementary Figure 8.** Comparison of genetic differentiation patterns across taxonomic levels in the *mulatta* species group. **A** gene-wise fixation index (*F*st) at subspecies-level, interspecies-level, and between DGD versus all other lineages. DGD exhibits significantly higher genetic differentiation compared to interspecies-level differentiation (Wilcoxon signed-rank test, ***$P < 0.001$). **B** Principal Component Analysis incorporating pairwise *F*st estimates, demonstrating distinct genetic clustering of DGD relative to recognized species *M. cyclopsis* and *M. fuscata*. The magnitude of differentiation between DGD and these established species parallels interspecific divergence levels.

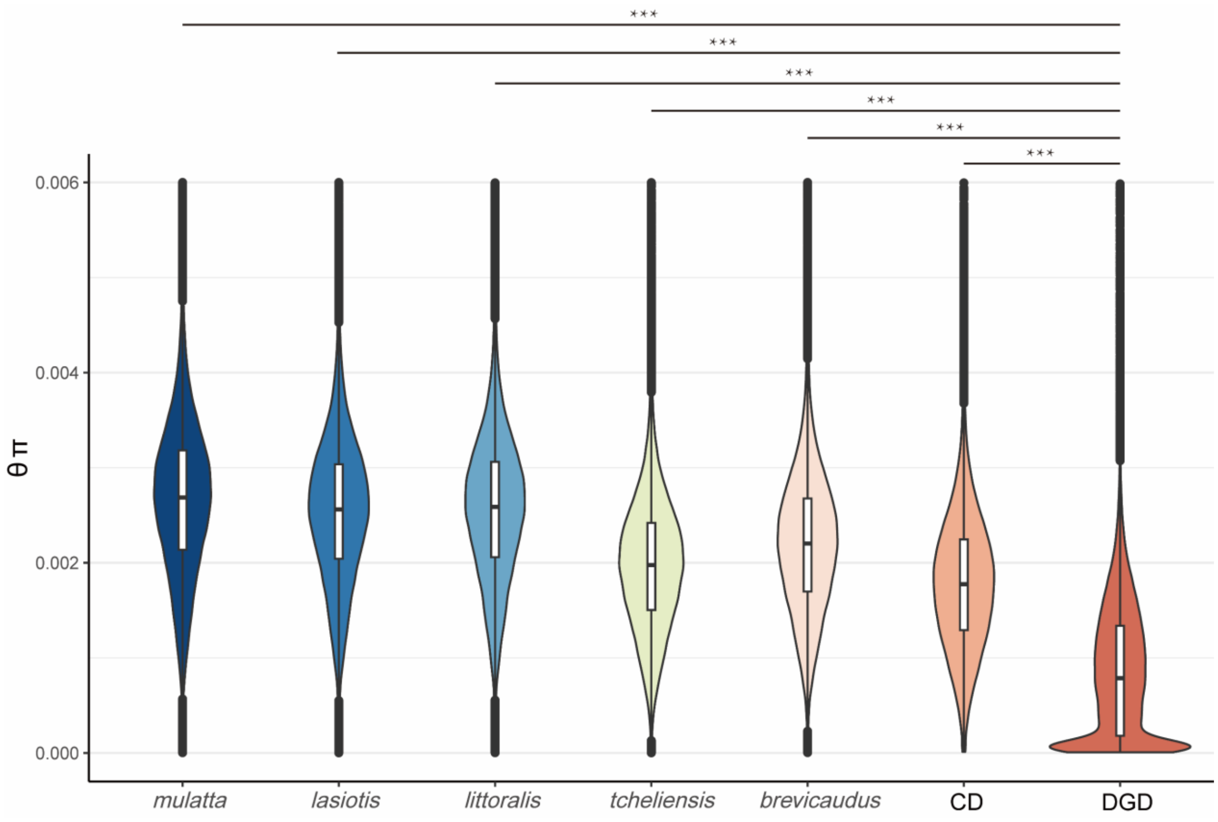


**Supplementary Figure 9.** Nucleotide diversity (θπ) across all studied *M. mulatta* subspecies and populations. The nucleotide diversity (θπ) of DGD is significantly lower than that of all others (Welch's t-test, ***P < 0.001).

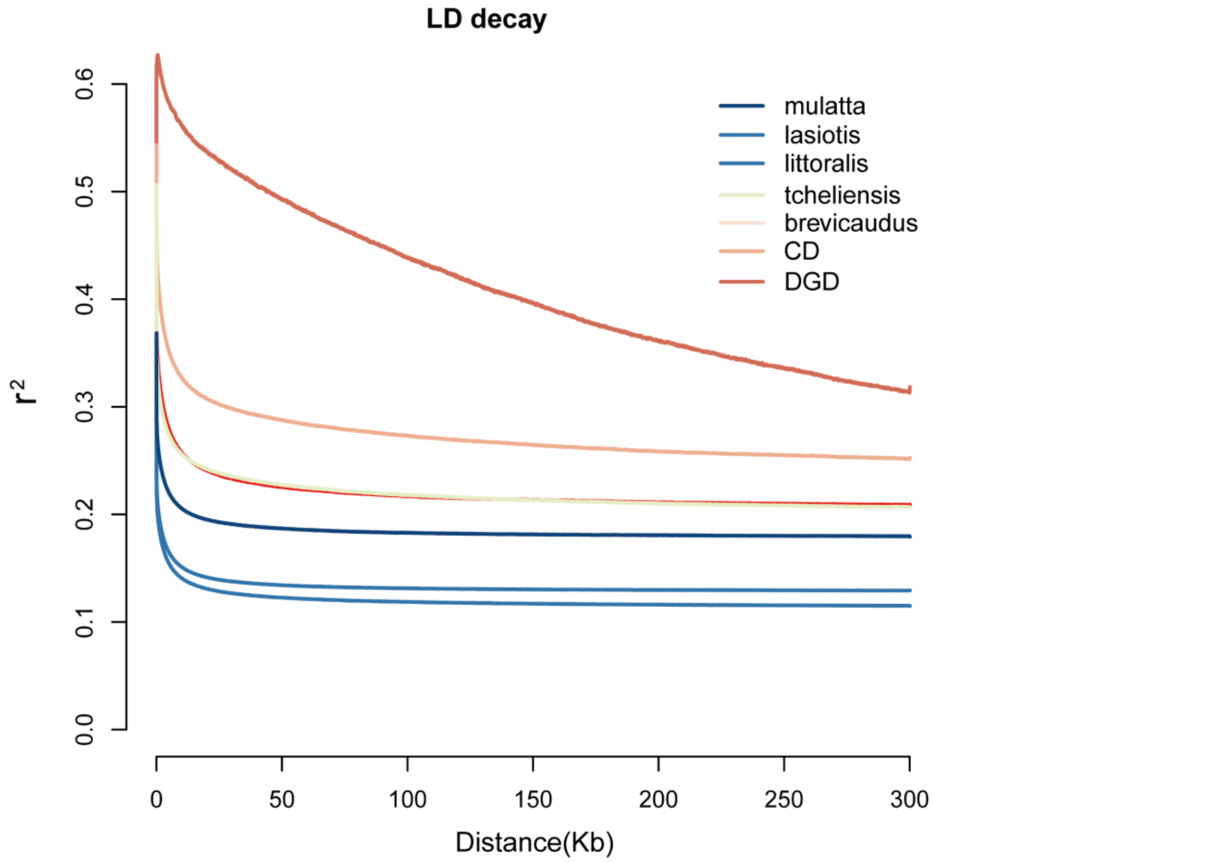


**Supplementary Figure 10.** Decay of linkage disequilibrium (LD) patterns measured with the squared coefficient of correlation ($r^2$) based on the genome-wide SNPs from different populations of all studied *M. mulatta* subspecies and populations.

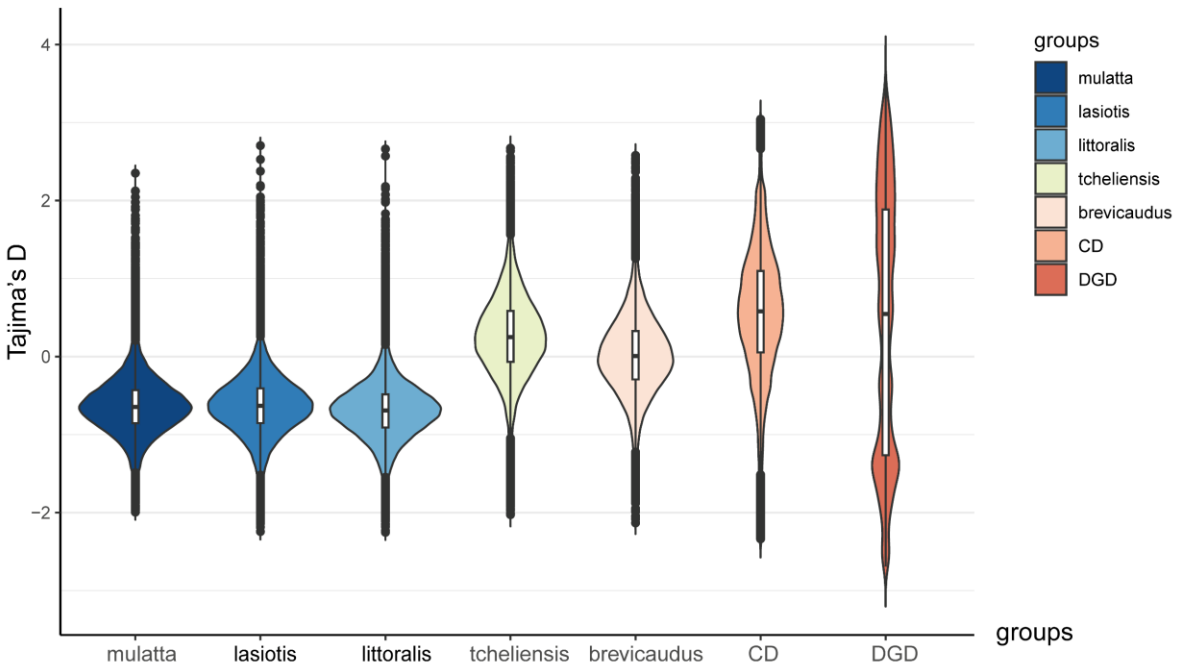


**Supplementary Figure 11.** Genome-wide Tajima's D, calculated in 25 kb steps overlapping a 50 kb sliding window for all studied *M. mulatta* subspecies and populations.

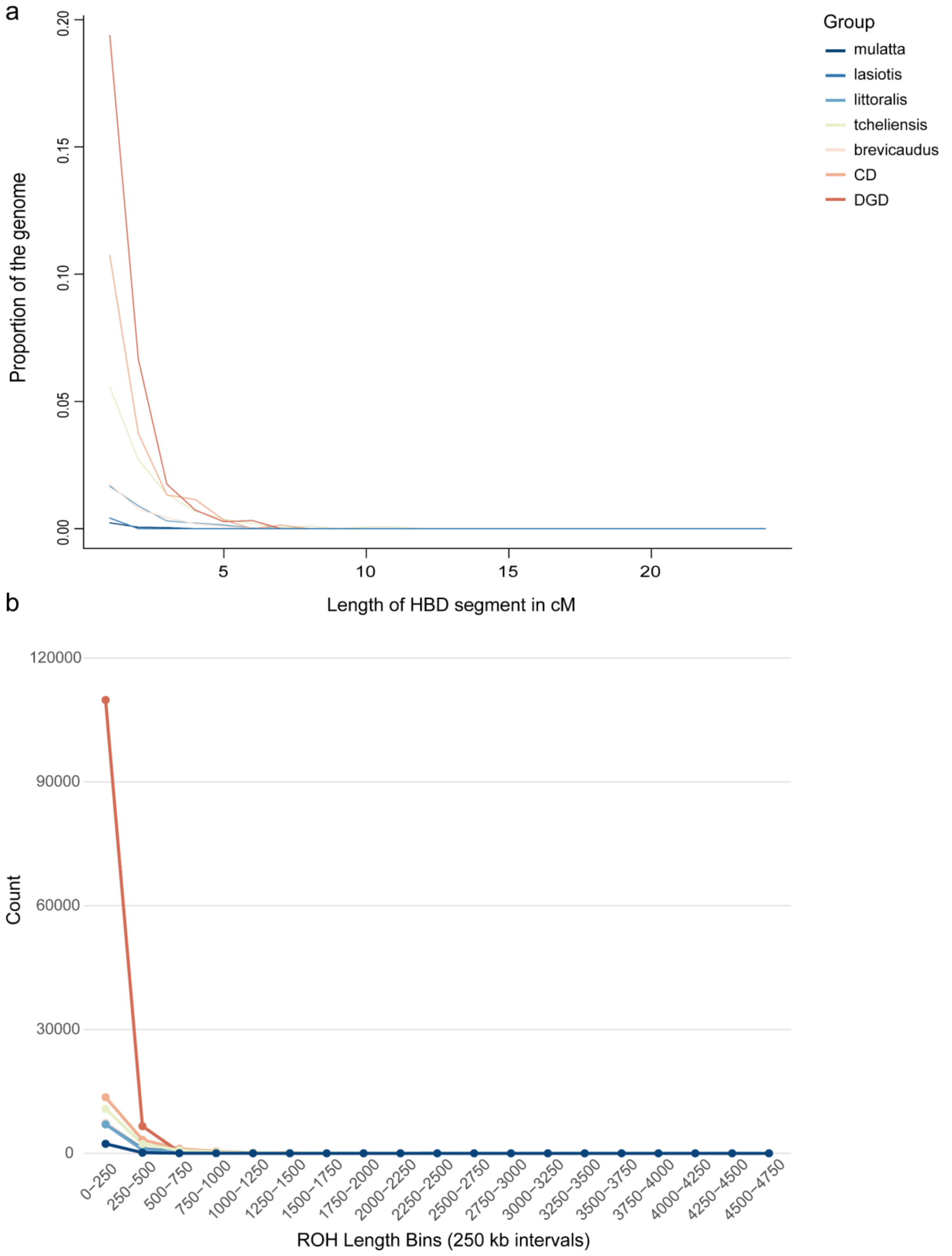


**Supplementary Figure 12.** Comparative analysis of genomic homozygosity patterns: Correlation between Runs of Homozygosity (ROH) fragment length and their genomic number/proportion estimated by RZooRoH (upper panels) versus PLINK (lower panels). The figure shows that both methods detected DGD with a large number of short ROH fragments. PLINK is less than 750 kb in length and RZooRoH is less than 5cm Morgen distance.

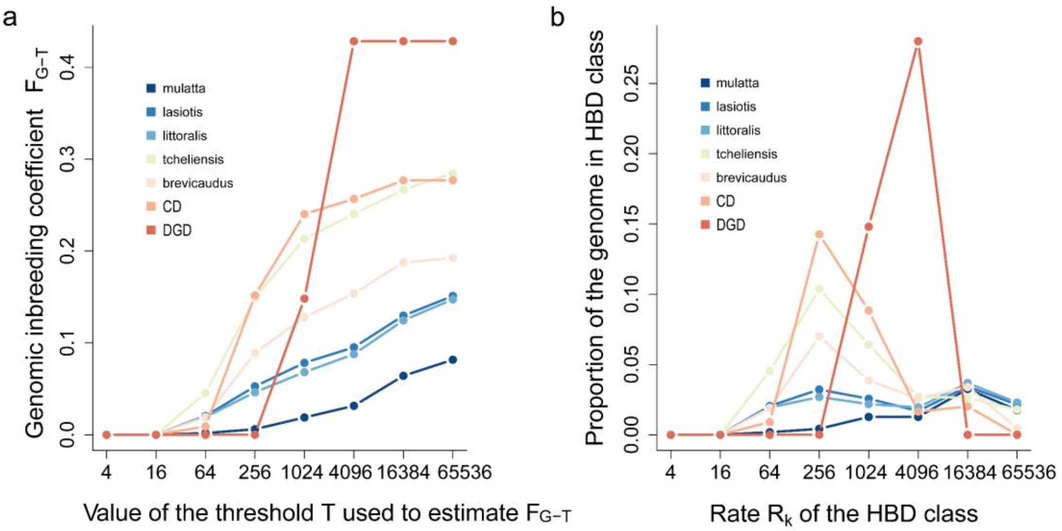


**Supplementary Figure 13.** The average inbreeding coefficients for all studied *M. mulatta* subspecies and populations were estimated using HBD classes and the proportion of the whole genome covered by each HBD class. The figure shows that the HBD fragments observed in the 1024 - 4096 generations have the highest inbreeding coefficients (cumulative) as well as the proportion of the genome (non-cumulative).

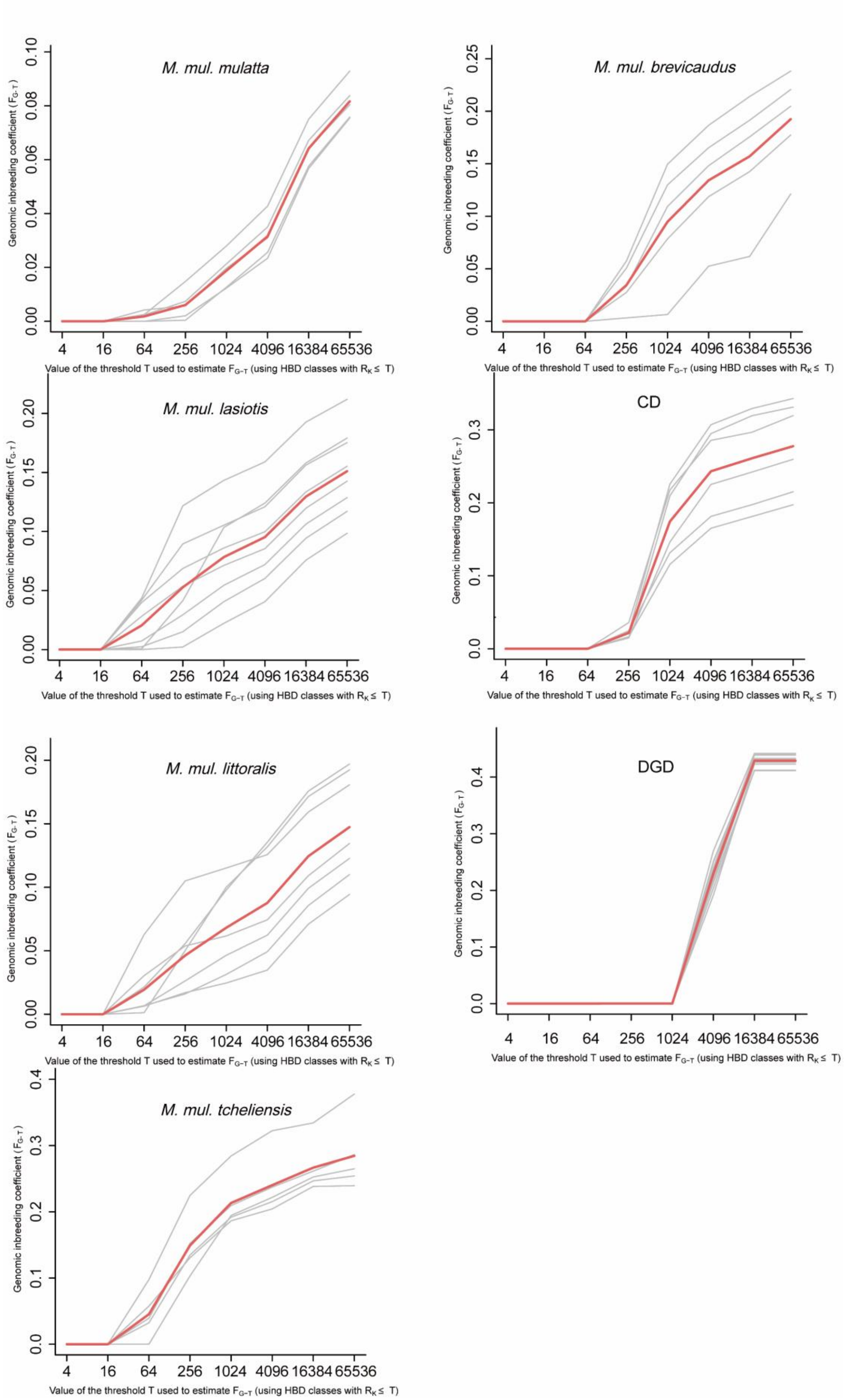

M. mul. mulatta
M. mul. brevicaudus
M. mul. lasiotis
CD
M. mul. littoralis
DGD
M. mul. tcheliensis
Genomic inbreeding coefficient ( $F_{G-T}$ )
Value of the threshold T used to estimate $F_{G-T}$ (using HBD classes with $R_K \leq$ T)
4 16 64 256 1024 4096 16384 65536

**Supplementary Figure 14.** The average inbreeding coefficient was estimated using the HBD class for each of the studied *M. mulatta* subspecies and populations. Shown is the distribution of every individual in each population (grey lines) and mean (red line) (see also Fig. S12). All of the inbred segments in the DGD population originated from very consistent generations, implying that they have experienced the same demographic history.

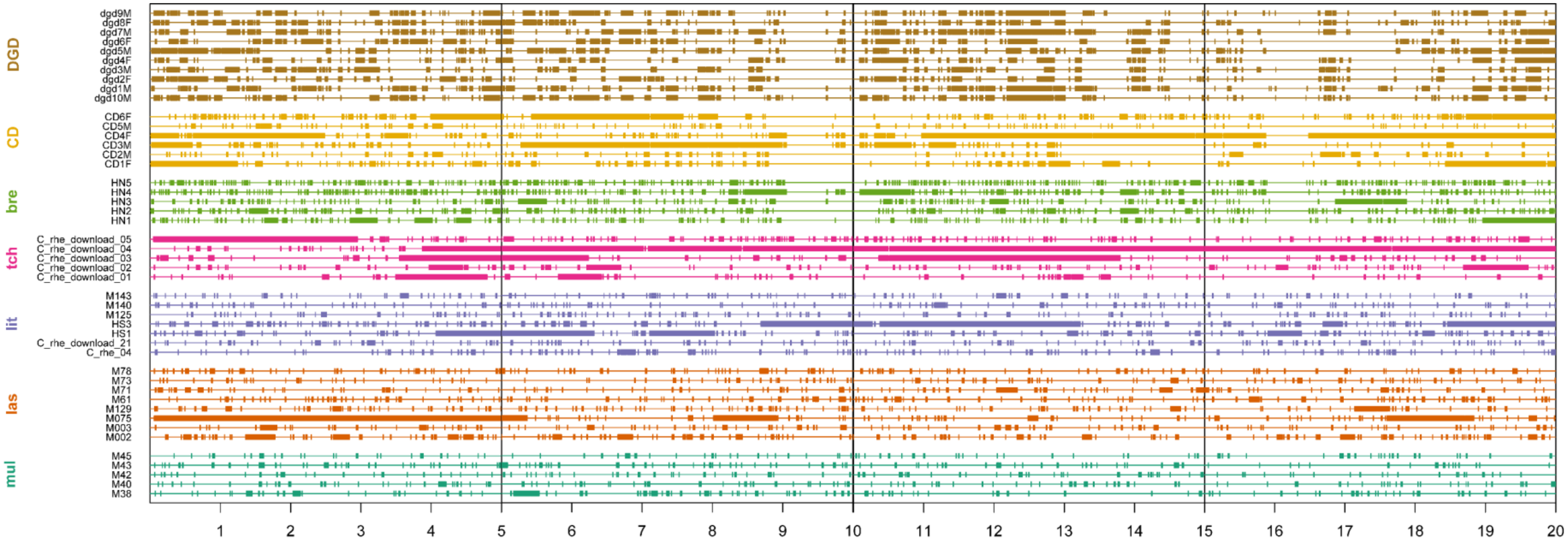


**Supplementary Figure 15.** Local visualization of all HBD fragments on chromosome 19 for all studied *M. mulatta* subspecies and populations. In the 20MB region of chromosome 19 DGD appears to have a large number of consecutively short ROH. While *M. m. tcheliensis* and CD appeared to have longer ROH fragments. This is consistent with other statistics. mul is *Macaca mulatta mulatta*; las is *Macaca mulatta lasiotis*; lit is *Macaca mulatta littoralis*; tch is *Macaca mulatta tcheliensis*; bre is *Macaca mulatta brevicaudus*.

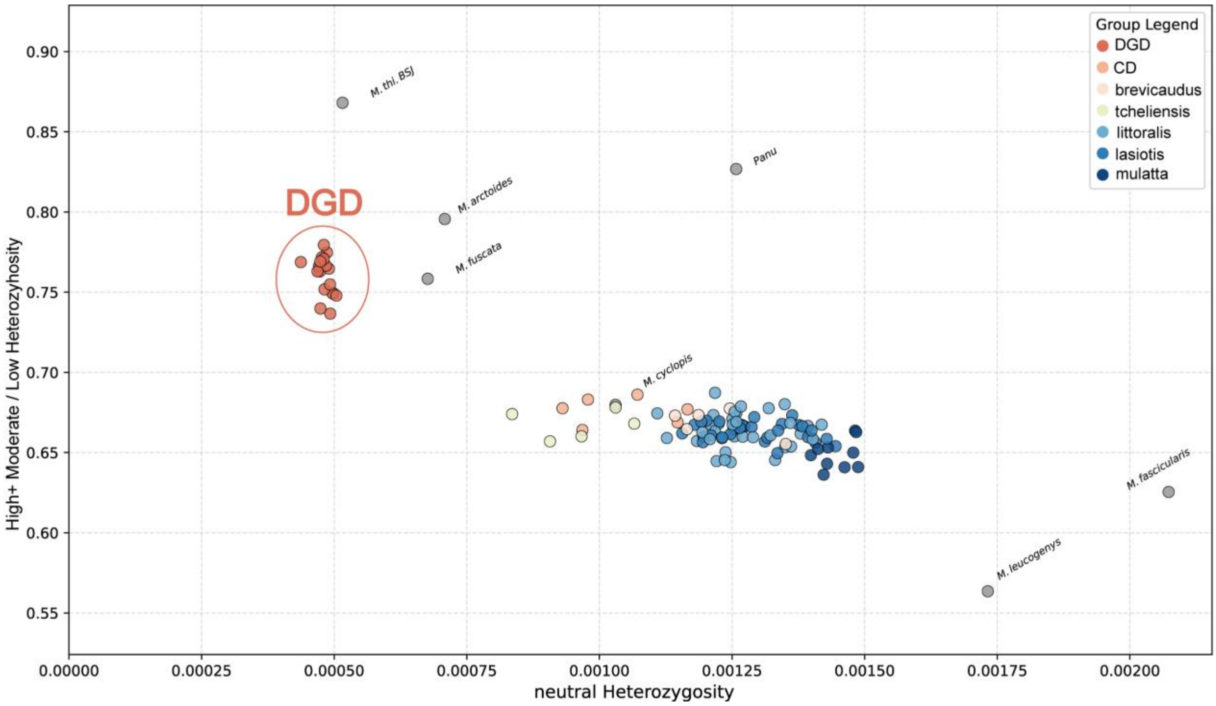


**Supplementary Figure 16.** Negative relationship between neutral diversity (influenced by effective population size) and ratio of heterozygosity at high+moderate impact variants to low-impact variants (influenced by the efficacy of purifying selection) (same as in Fig. 3B).

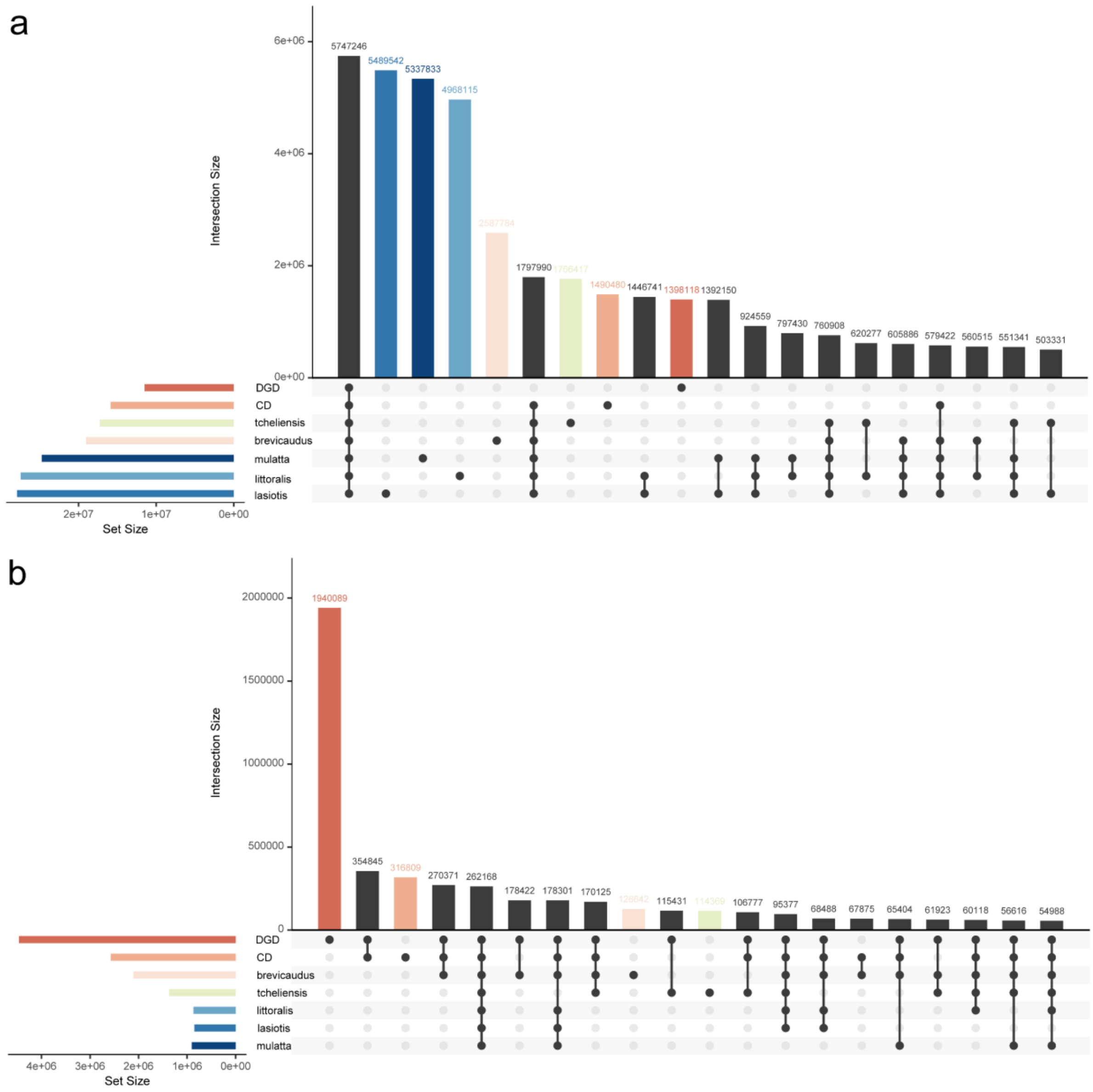


**Supplementary Figure 17.** Upset diagrams showing genetic variation and fixed variant across studied *M. mulatta* subspecies and populations. **A.** Intersection analysis of SNPs across all study groups relative to the Mmul_10 reference genome. Population-specific variant counts reflect sequential founder effects and demographic history. The DGD population (continent-derived) exhibits a markedly reduced total variant count, consistent with its three-phase founder effect and historical demographic bottleneck. **B.** Distribution of shared and fixed SNPs across all study groups. Intersection sizes (y-axis) represent shared variants, while unique variants (x-axis)

are stratified by population (DGD: red; CD: orange; *brevicaudus*: pink; tcheliensis: cyan). Pronounced fixation of lineage-specific SNPs in DGD highlights strong genetic drift effects, with accelerated fixation rates driving substantial divergence in this population. These patterns collectively underscore drift-driven evolution following population isolation and bottleneck events.

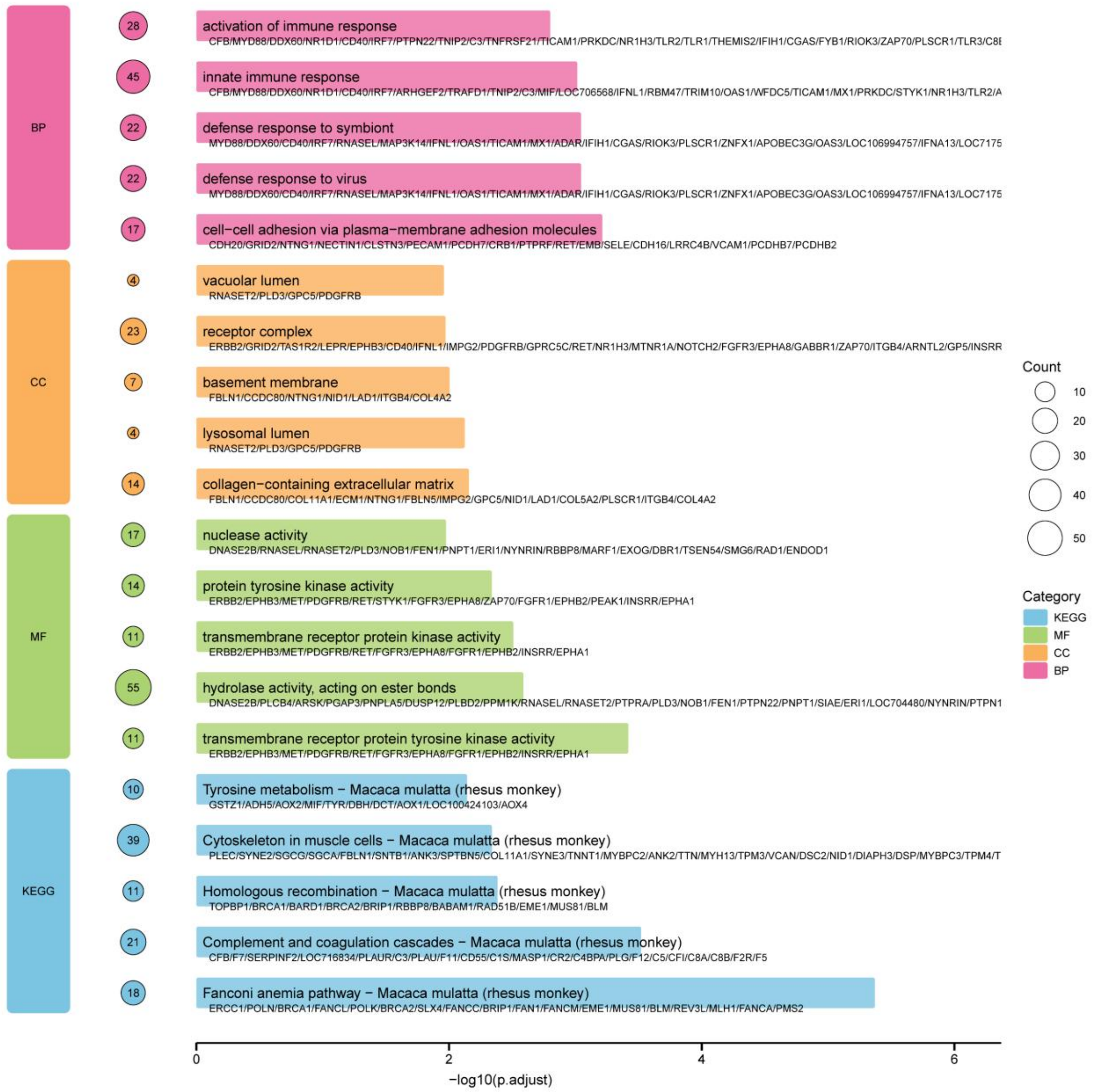


**Supplementary Figure 18.** Enrichment analysis for Gene Ontology (GO) and KEGG pathway of fixed missense gene set. The top 5 significantly enriched GO terms and KEGG terms are displayed.

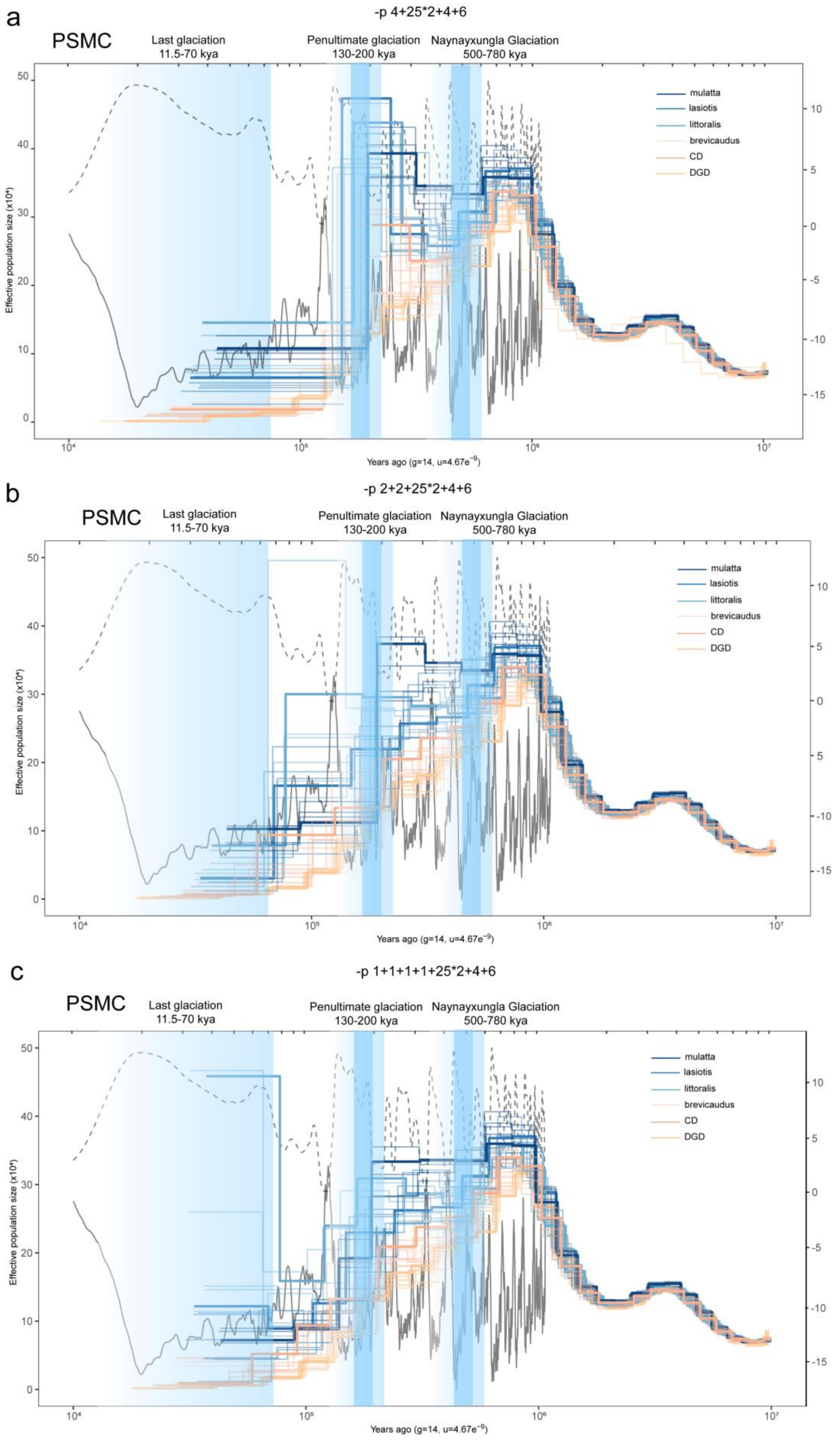
a
-p 4+25*2+4+6
PSMC
Last glaciation
11.5-70 kya
Penultimate glaciation
130-200 kya
Naynayxungla Glaciation
500-780 kya
mulatta
lasiotis
littoralis
brevicaudus
CD
DGD
Effective population size (x10⁴)
Years ago (g=14, u=4.67e⁻⁹)
b
-p 2+2+25*2+4+6
PSMC
Last glaciation
11.5-70 kya
Penultimate glaciation
130-200 kya
Naynayxungla Glaciation
500-780 kya
mulatta
lasiotis
littoralis
brevicaudus
CD
DGD
Effective population size (x10⁴)
Years ago (g=14, u=4.67e⁻⁹)
c
-p 1+1+1+1+25*2+4+6
PSMC
Last glaciation
11.5-70 kya
Penultimate glaciation
130-200 kya
Naynayxungla Glaciation
500-780 kya
mulatta
lasiotis
littoralis
brevicaudus
CD
DGD
Effective population size (x10⁴)
Years ago (g=14, u=4.67e⁻⁹)

**Supplementary Figure 19.** Demographic history of Chinese rhesus macaque subspecies and populations inferred using the PSMC method. We implemented different parameters, which have been labeled above the image (A. -p 4+25*2+4+6 B. -p 2+2+25*2+4+6 C. -p 1+1+1+1+25*2+4+6). Each line represents an individual macaque, colored by its subspecies/population assignment. The black dashed line represents global sea level relative to the present (×10M). The black solid curve represents surface air temperature relative to the present.

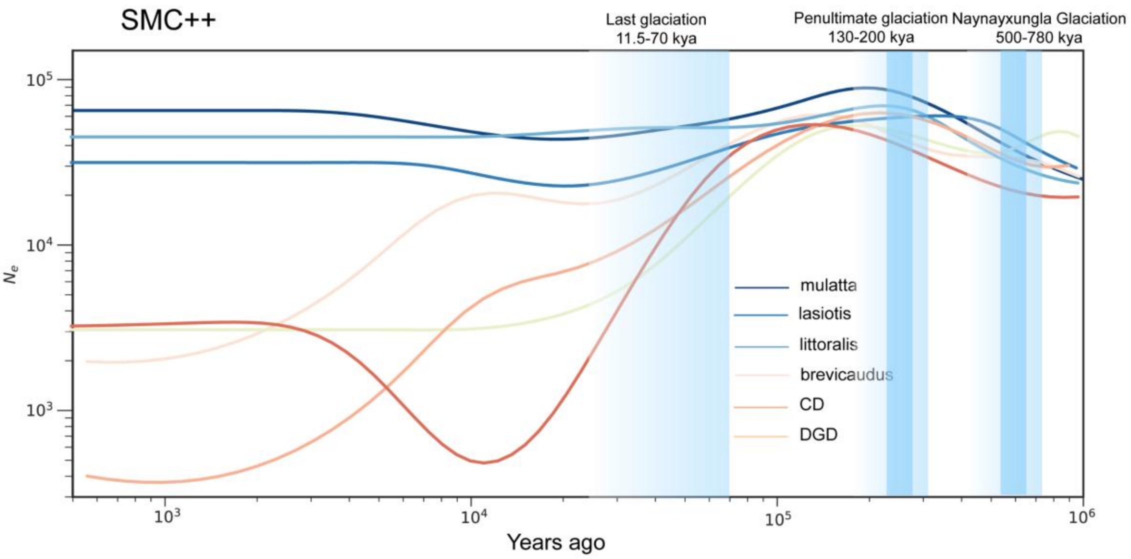


**Supplementary Figure 20.** Changes in effective population size (Ne) for all studied *M. mulatta* subspecies and populations inferred by SMC++. Each colored line represents one studied subspecies or populations.

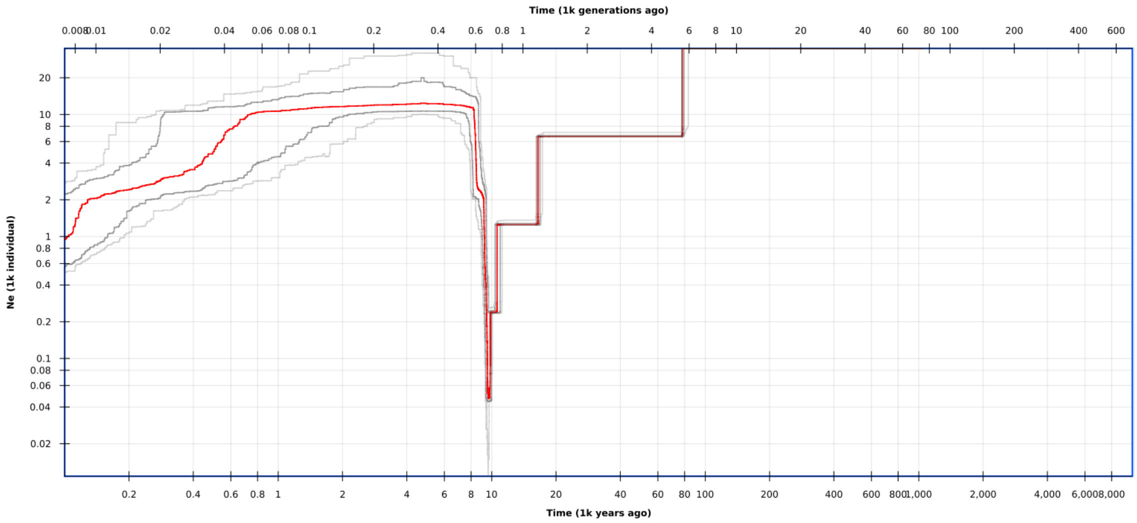


**Supplementary Figure 21.** Inferred demographic history by StairwayPlot2 inference with folded SFSs based on 20 DGD individuals. The solid red line refers to the median of 200 inferences based on subsampling. Dark and light gray lines refer to 75% and 95% confidence intervals of the inference, respectively.

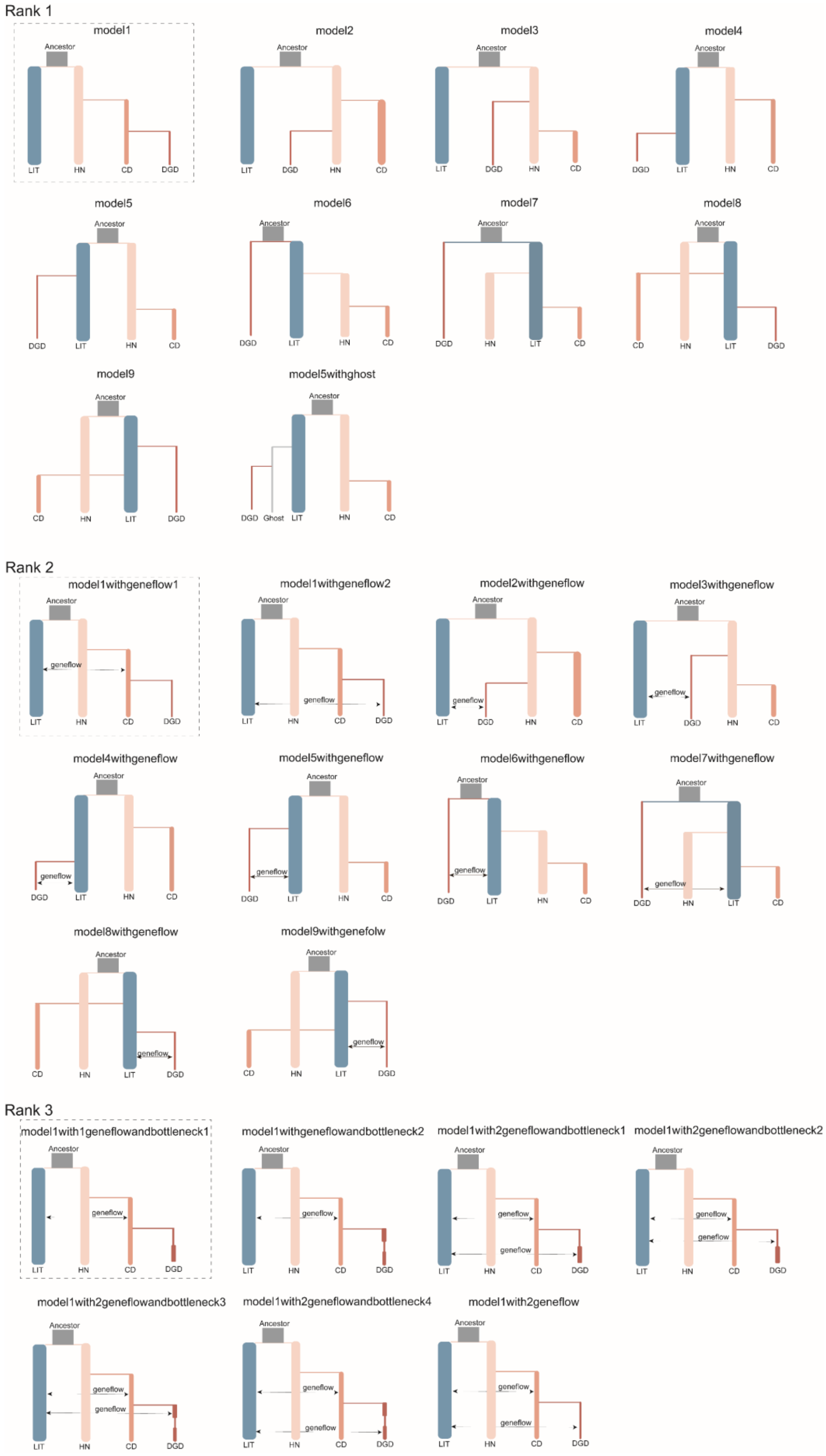
Rank 1
model1
model2
model3
model4
model5
model6
model7
model8
model9
model5withghost
Ancestor
LIT
HN
CD
DGD
Ghost
Rank 2
model1withgeneflow1
model1withgeneflow2
model2withgeneflow
model3withgeneflow
model4withgeneflow
model5withgeneflow
model6withgeneflow
model7withgeneflow
model8withgeneflow
model9withgenefolw
geneflow
Rank 3
model1with1geneflowandbottleneck1
model1withgeneflowandbottleneck2
model1with2geneflowandbottleneck1
model1with2geneflowandbottleneck2
model1with2geneflowandbottleneck3
model1with2geneflowandbottleneck4
model1with2geneflow

**Supplementary Figure 22.** Alternative Demographic Models and Best-Fit Model for Historical Inferences using fastsimcoal2. The figure shows the sequence of demographic model testing conducted across three iterative rounds of coalescent simulations in fastsimcoal2. The dashed box highlights the best-fit model identified in each round. First, we tested fundamental divergence scenarios for DGD, including a "ghost population" model where DGD diverged from mainland *M. m. littoralis* via unsampled intermediates before island isolation. Building upon the best Round 1 model, the second round incorporated potential gene flow events, guided by *D*-statistics, *F*-branch, *f3*-statistics, and TreeMix analyses, to investigate secondary contact along the southern Chinese coast following initial divergence. Finally, extending the best Round 2 model, the third round focused on the population bottleneck associated with land-sea isolation, specifically assessing the likelihood and timing of potential secondary contact for the DGD population: either before the bottleneck (mainland mixing) or after (e.g., via drift or human-mediated dispersal post-isolation). The demographic scenario from the overall best-fit model (Round 3) provides estimates of divergence times and effective population sizes (detailed in Supplementary Tables S8, S9 and S10).

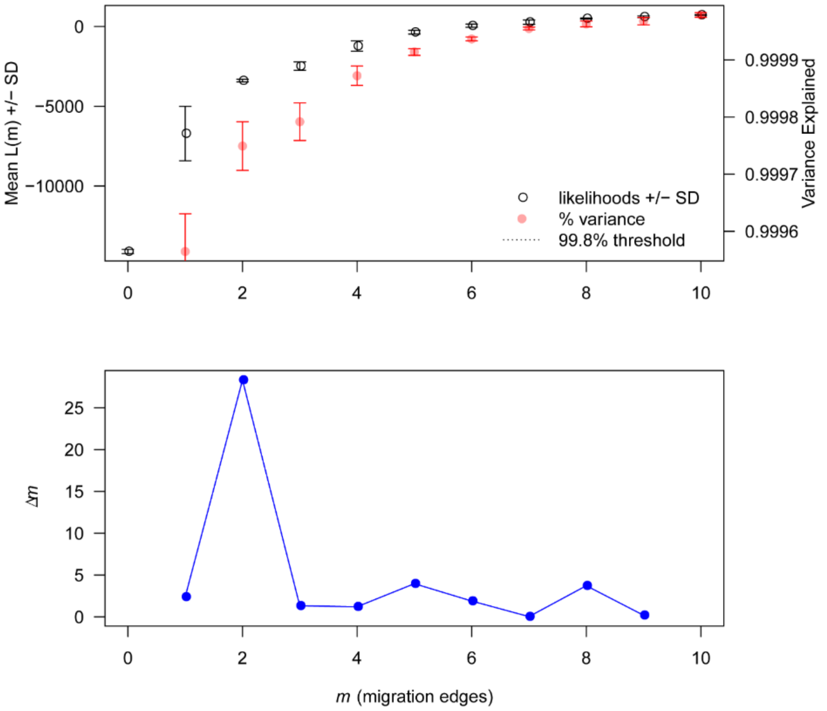


**Supplementary Figure 23.** Selection of the most likely number of migration edges by the OptM R package. Distribution of log likelihood and variance explained of Treemix models with 0 - 10 migration edges. The number of migration edges that explains ~98% of the variance is chosen as the optimal number of edges. **c** Selection of the most likely number of migration edges by plotting *Δm*.

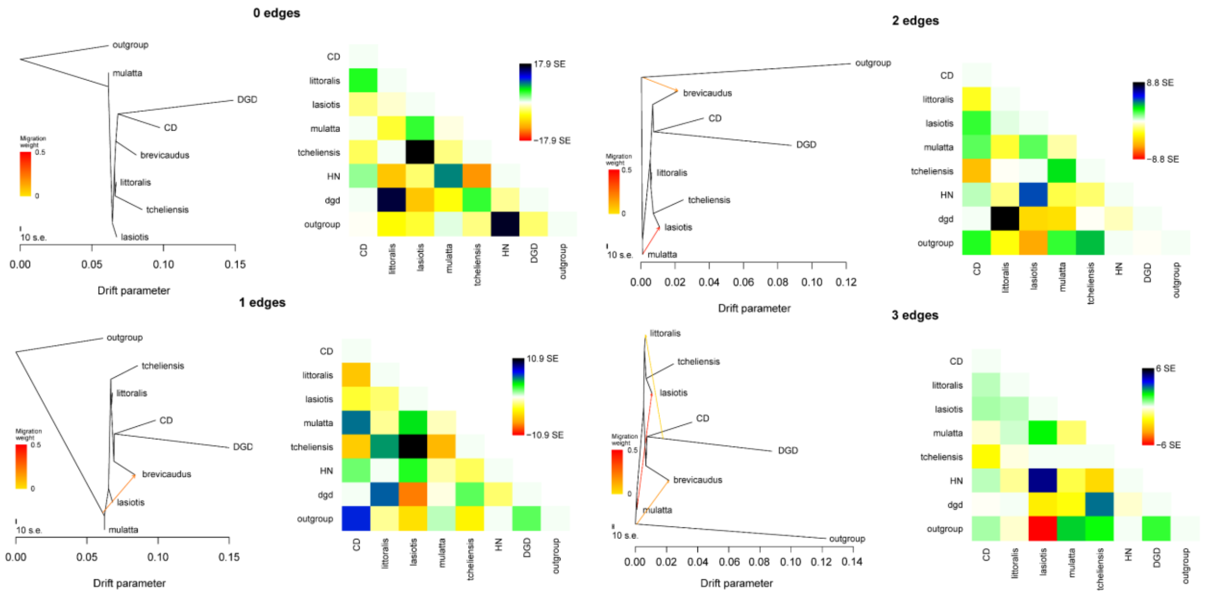


**Supplementary Figure 24.** Inference of population splits and admixtures with TreeMix. Shown are edges from 0 to 4, maximum likelihood (ML) tree assuming 0 - 3 migration edge (left panel) and corresponding model residual fit (right panel). Horizontal branch lengths are proportional to the amount of genetic drift that has occurred on the branch. The arrows in the 4 edges image point to potential mixing events occurring on the continent.

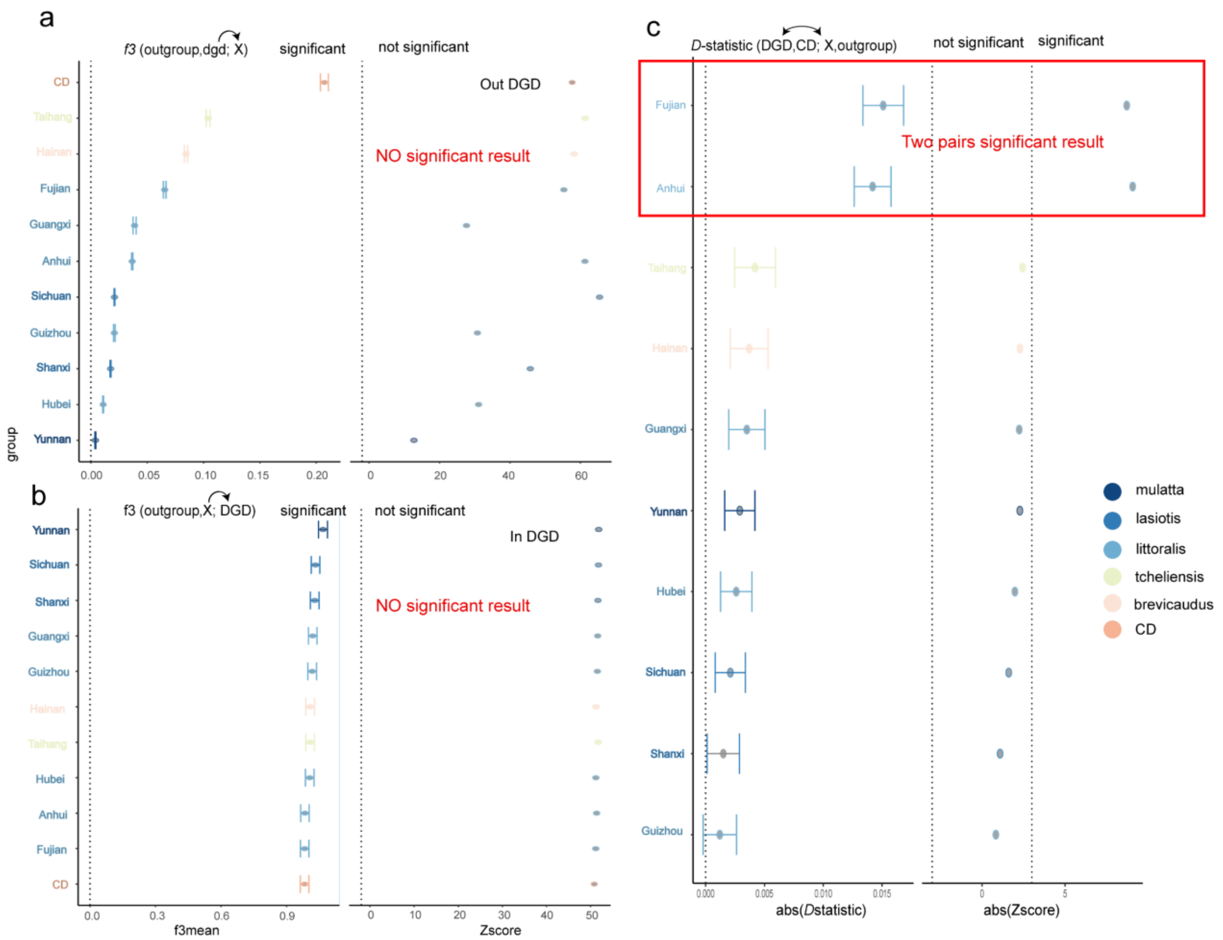


**Supplementary Figure 25.** Results of *f3*-statistics and *D*-statistics.

A. f3-statistics (DGD as source population) Results of *f3*-statistics testing all potential sink populations receiving gene flow from DGD. Negative mean *f3* values (with SD) and Z-scores < -2 indicate gene flow from DGD to sink X.

B. *f3*-statistics (DGD as sink population) Results of f3-statistics testing all potential source populations contributing to DGD. Negative mean f3 values (with SD) and Z-scores < -2 indicate gene flow into DGD from source X.

C. *D*-statistics for admixture testing. *D*-statistics of the form D (DGD, CD; X, outgroup) across studied populations. Values and associated Z-scores are shown. dash line $|Z| \geq 3$ signifies significant admixture between DGD and population X.

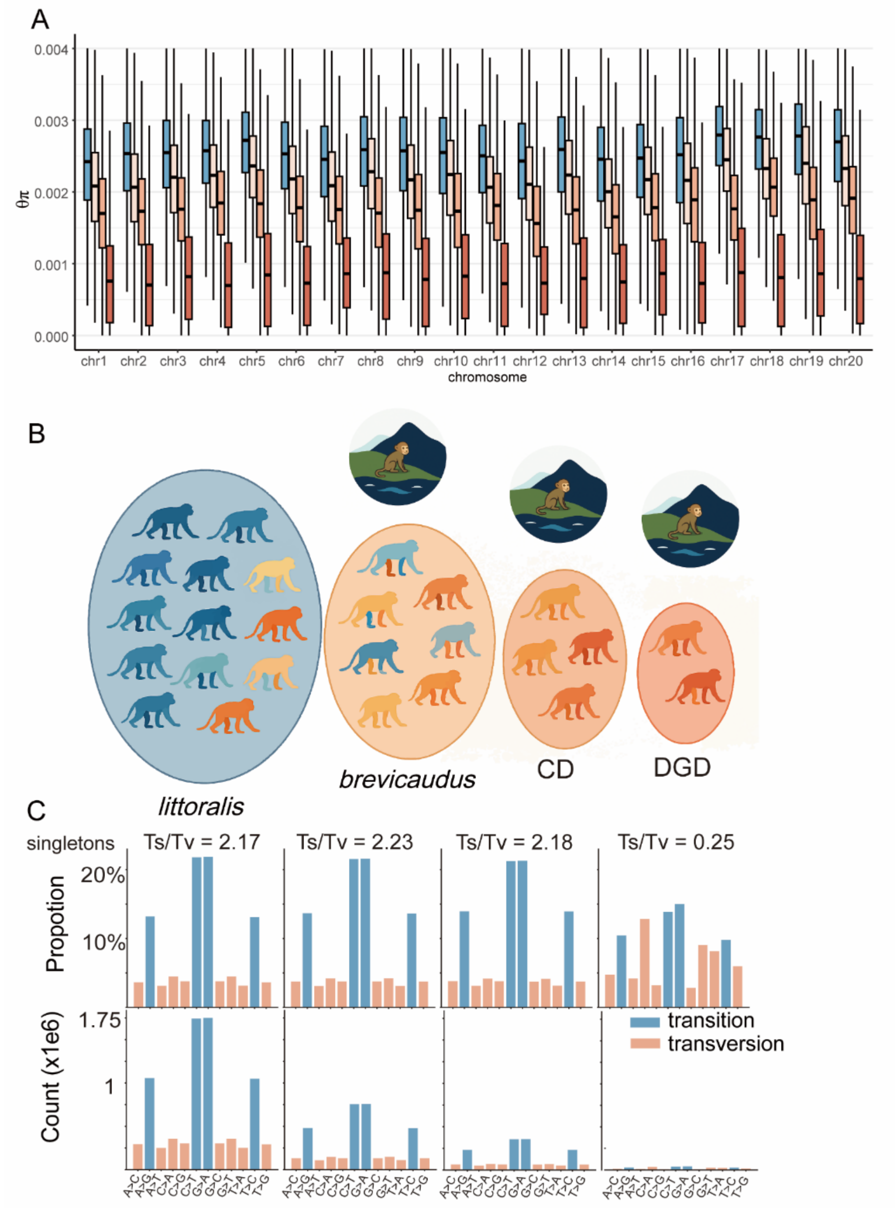


**Supplementary Figure 26.** The erosion of genomic genetic diversity and shifts in mutation spectrum caused by the three-step founder effect. **A.** Chromosome-wide diversity erosion. Genetic diversity (θπ) for all chromosomes using a three-step founder-event cascade. DGD lost 65.8% of its diversity post-isolation from mainland littoralis. **B.** Scheme of genomic erosion through founder events. Cumulative diversity loss (color gradient: blue→orange→red) through serial founder effects. The circles represent effective population size (Ne). **C.** Mutation spectrum alteration. Top: Singletons Transition/Transversion (Ti/Tv) ratio. Bottom: Absolute mutation counts. DGD shows a decrease in overall variation and low Ti/Tv (0.25), which reflect strong purifying selection against *de novo* mutations.

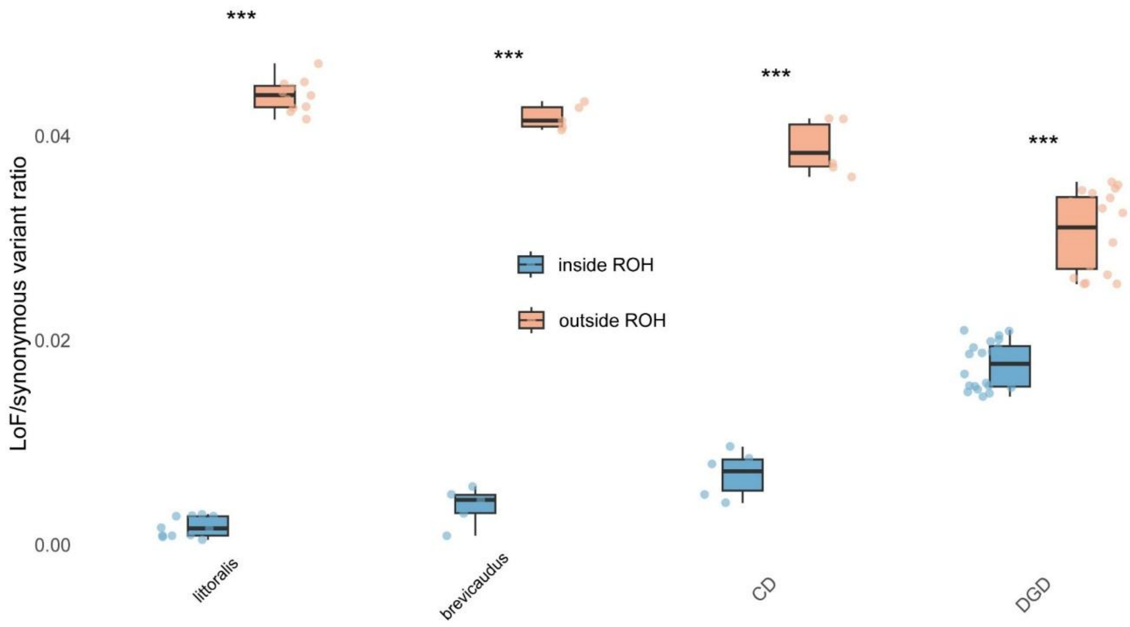


**Supplementary Figure 27.** Rates of LoF variants in runs of homozygosity (ROH). Blue and orange show the rate of LoF variants relative to synonymous variants inside and outside ROH, respectively, for each individual genome. Middle lines represent means (Welch's two-sample t-test; ***P<0.001). There was a significantly lower number of LoF alleles inside ROH compared to heterozygous parts of individual genomes in both populations. And this difference was 68% smaller in the DGD population compare to *M. m. littoralis*, suggesting that inbreeding events may have facilitated the removal of a significant proportion of severely deleterious and recessive LoF alleles through exposure in homozygous state from the DGD population.

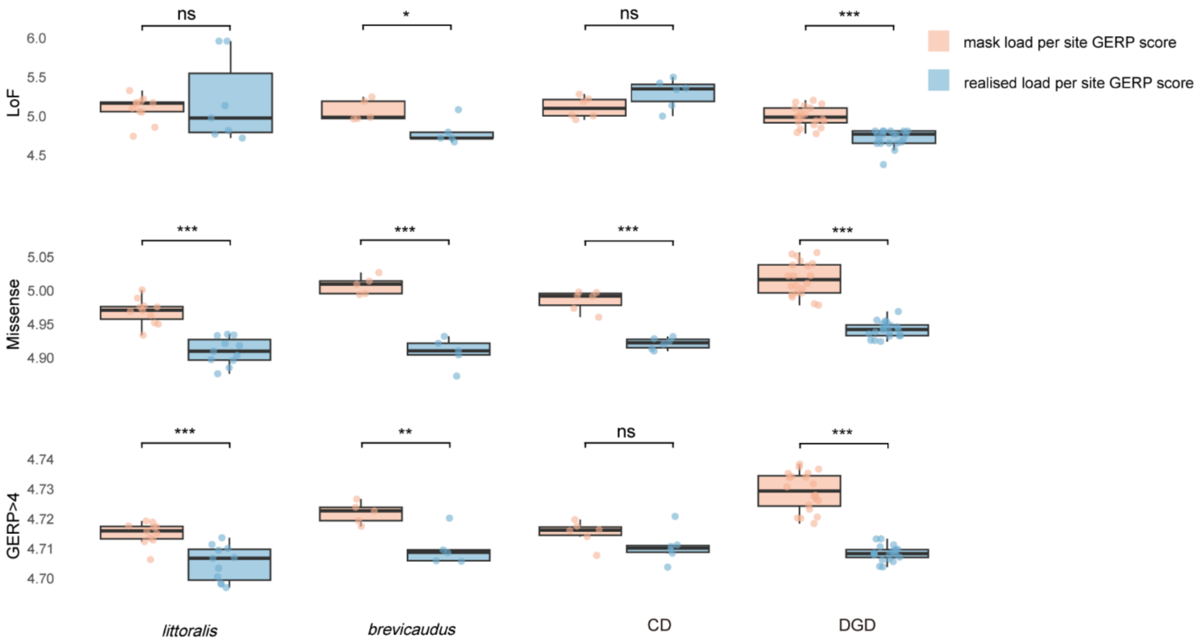


**Supplementary Figure 28.** Mean GERP scores of LoF, missense, and GERP>4 loci across four *M. mulatta* subspecies/populations. Realized load (homozygous state) exhibits significantly lower GERP scores than masked load (heterozygous state), indicating strong purifying selection against highly deleterious homozygous variants in surviving individuals.

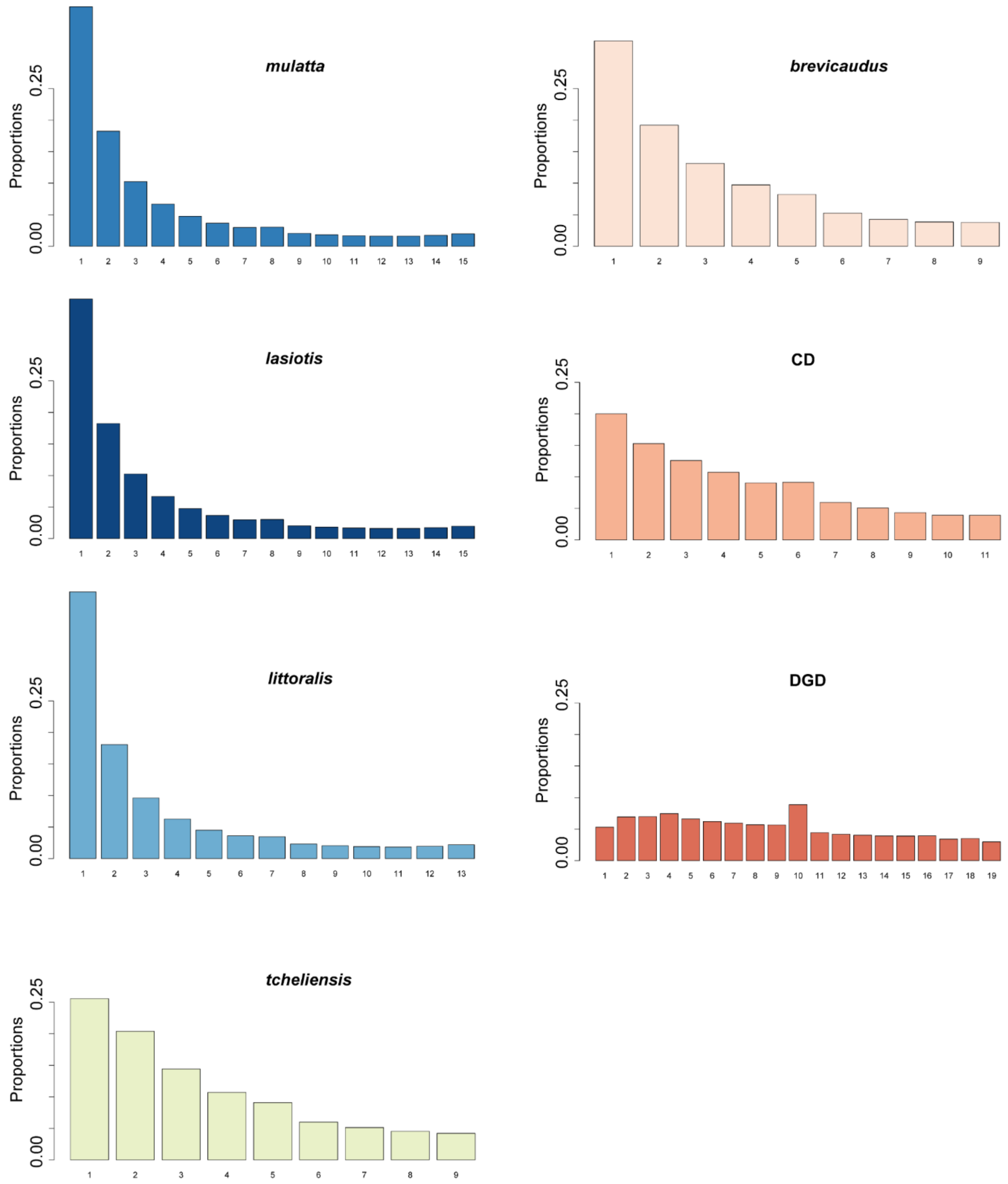


**Supplementary Figure 29.** Shape of the site frequency spectrum (SFS) for all studied *M. mulatta* subspecies and populations. DGD exhibits a nearly flat allele frequency distribution, reflecting attenuated selection efficacy and intensified genetic drift. This pattern manifests as reduced prevalence of low-frequency deleterious alleles alongside increased intermediate-frequency variants dominated by neutral processes.

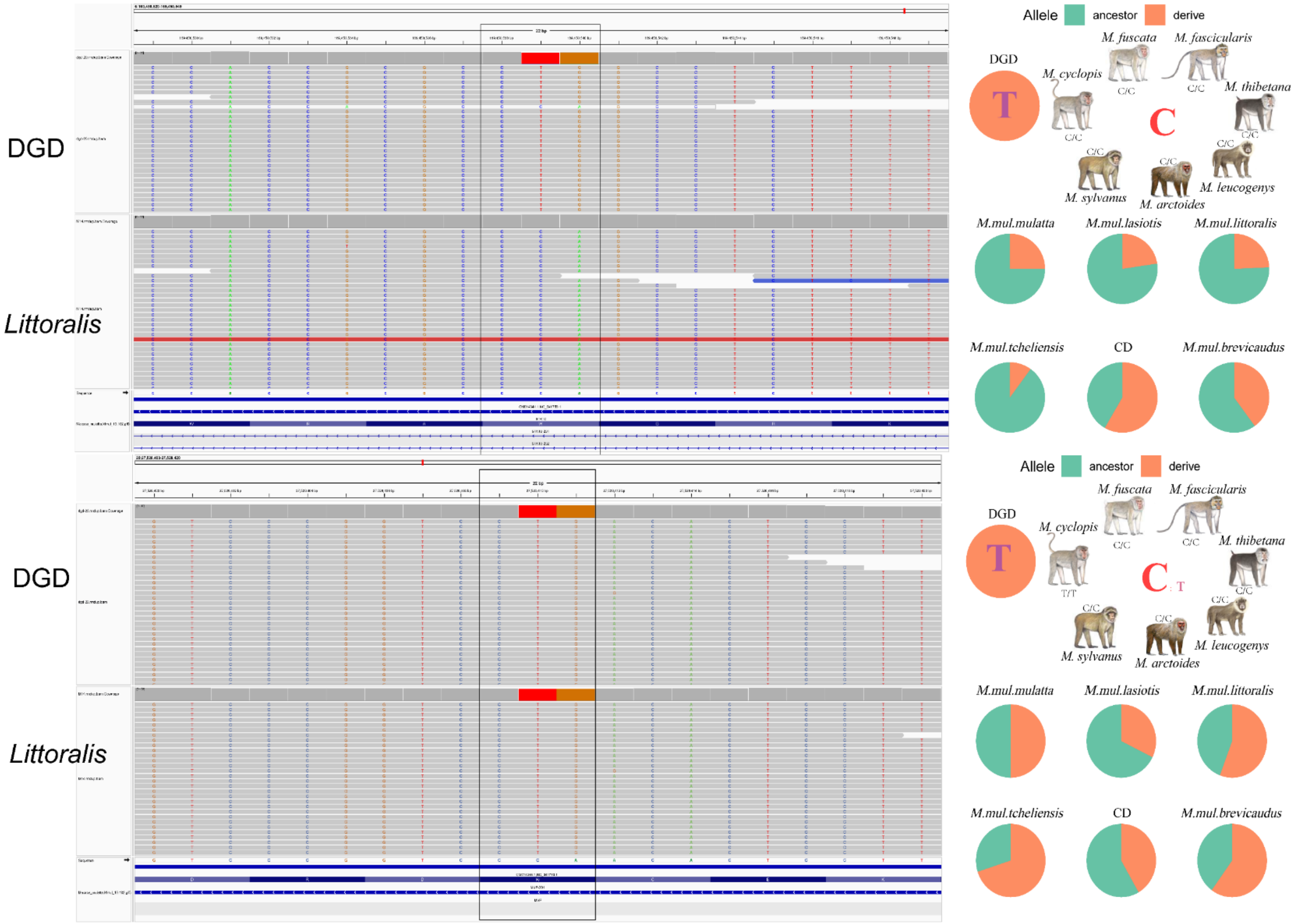
DGD
Littoralis
DGD
Littoralis
Allele
ancestor
derive
M. fuscata
M. fascicularis
M. cyclopis
M. thibetana
M. sylvanus
M. arctoides
M. leucogenys
T
C
M.mul.mulatta
M.mul.lasiotis
M.mul.littoralis
M.mul.tcheliensis
CD
M.mul.brevicaudus
Allele
ancestor
derive
M. fuscata
M. fascicularis
M. cyclopis
M. thibetana
M. sylvanus
M. arctoides
M. leucogenys
T
C
M.mul.mulatta
M.mul.lasiotis
M.mul.littoralis
M.mul.tcheliensis
CD
M.mul.brevicaudus

**Supplementary Figure 30.** Double codon mutations of putative Loss-of-Function Variants. IGV of candidate stop-gain variants at Chr6: 169,450,529-169,450,549 and Chr20: 202,752,840-202,752,860. Both loci exhibit dual in-codon mutations predicted by SnpEff as premature stop codons. However, nucleotide resolution reveals neither site forms a complete stop codon (TAG/TAA/TGA).

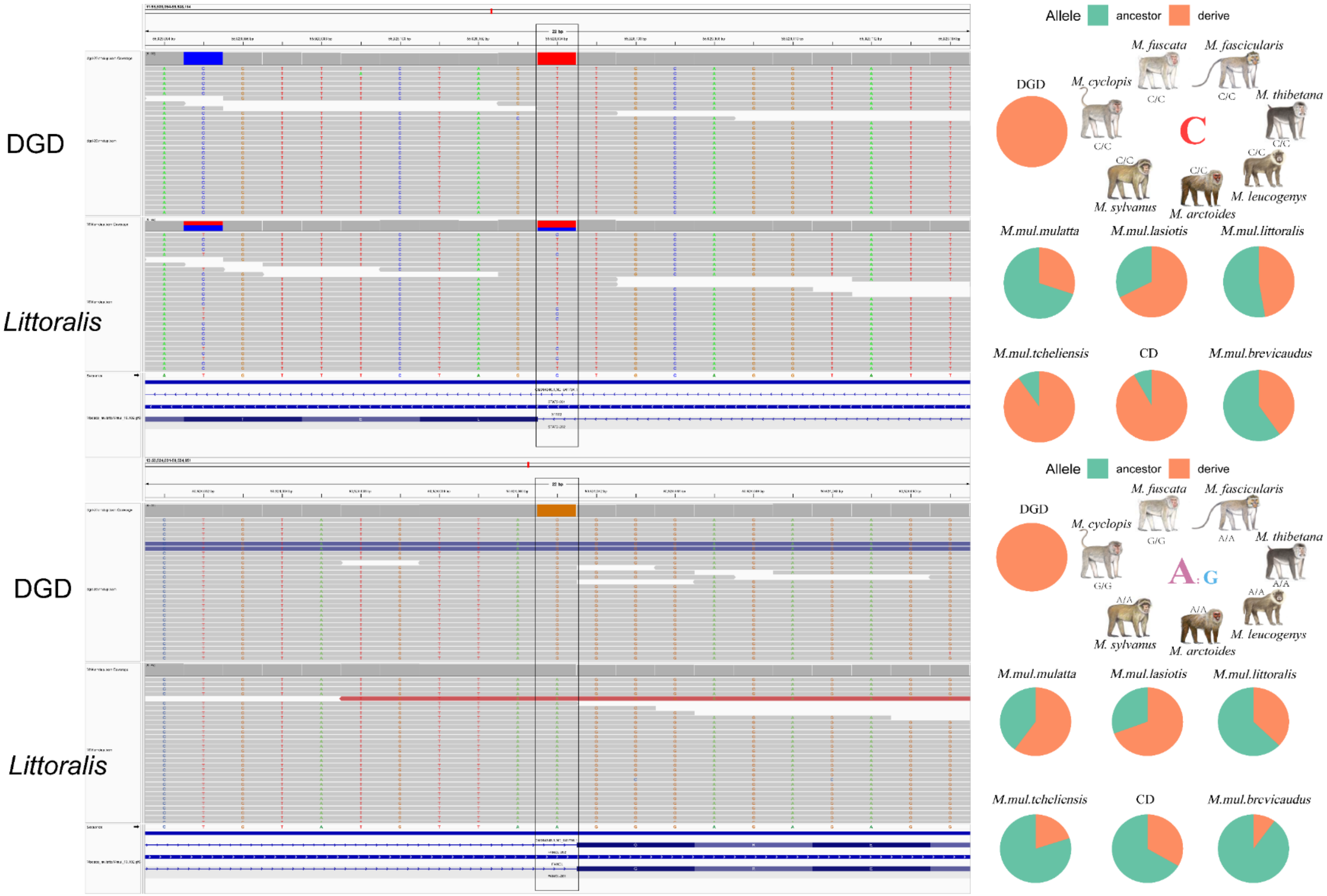


**Supplementary Figure 31.** Putative splice acceptor variants with high ancestral frequency suggest attenuated deleterious effects. IGV of candidate splice acceptor variants at Chr11:55,928,094-55,928,114 and Chr13:50,524,031-50,524,051. Both loci exhibit high ancestral population frequency mutation predicted by SnpEff as splice acceptor variants. However, the elevated ancestral frequency contradicts expected strong negative selection for deleterious LoF variants.

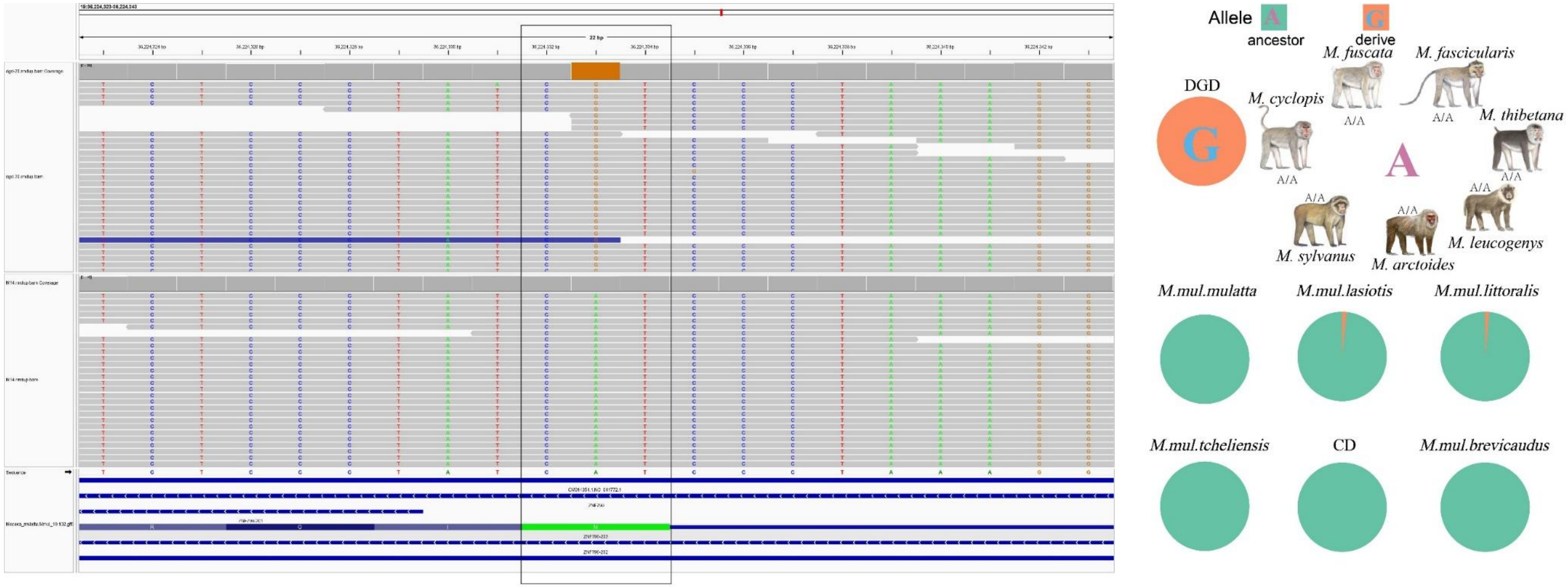


**Supplementary Figure 32.** ZNF790: A putatively ultra-deleterious fixed start lost loss-of-function (LoF) gene. Predicted to enable DNA-binding transcription factor activity with specificity for RNA polymerase II and cis-regulatory region sequence-specific DNA binding [54].

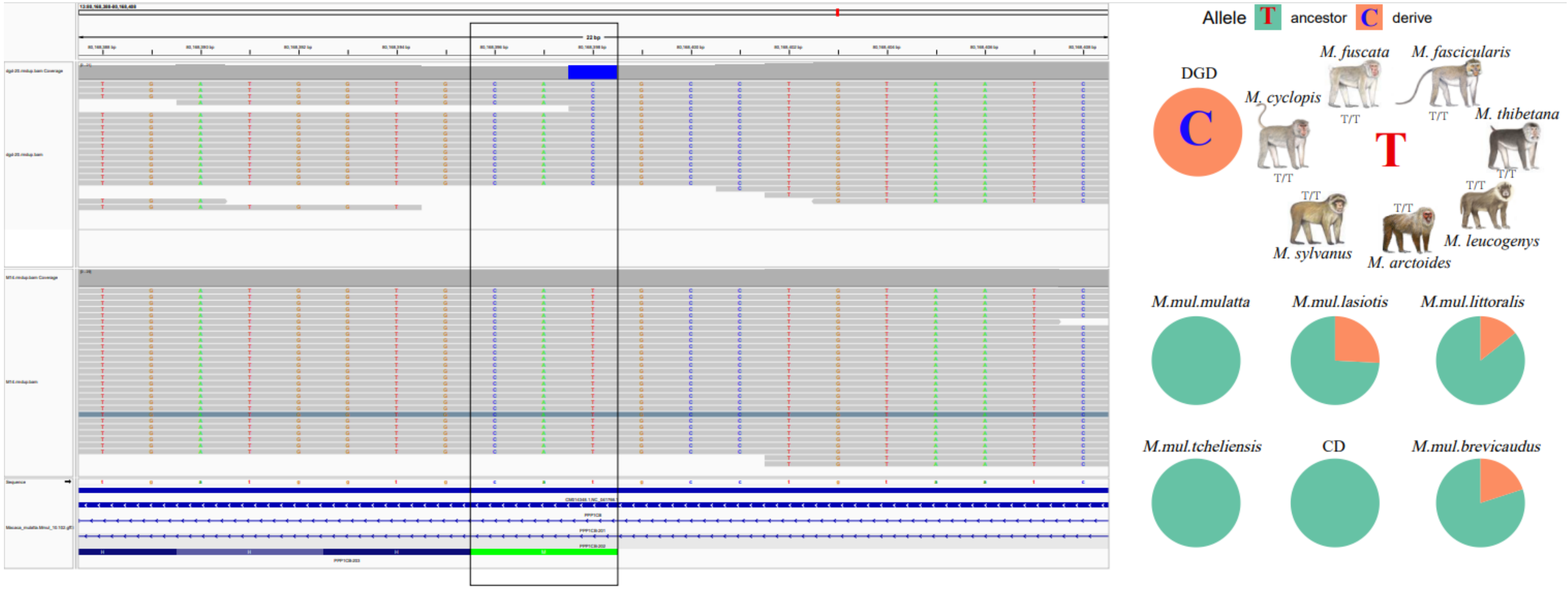


**Supplementary Figure 33.** PPP1CB: A putatively ultra-deleterious fixed start lost loss-of-function (LoF) gene. The protein encoded by this gene is one of the three catalytic subunits of protein phosphatase 1 (PP1). PP1 is a serine/threonine specific protein phosphatase known to be involved in the regulation of a variety of cellular processes, such as cell division, glycogen metabolism, muscle contractility, protein synthesis, and HIV-1 viral transcription[55].

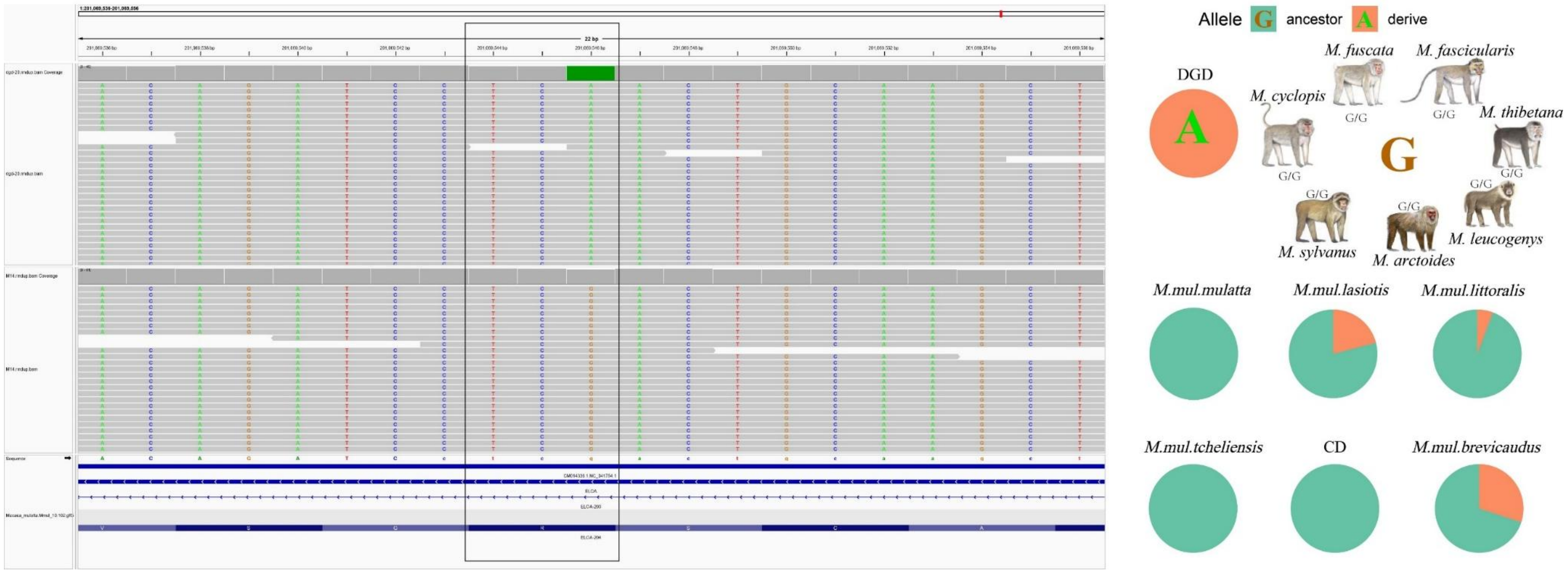


**Supplementary Figure 34.** ELOA: A putatively ultra-deleterious fixed stop gained loss-of-function (LoF) gene. This gene encodes the protein elongin A, which is a subunit of the transcription factor B (SIII) complex. The SIII complex is composed of elongins A/A2, B and C. It activates elongation by RNA polymerase II by suppressing transient pausing of the polymerase at many sites within transcription units. Diseases associated with ELOA include Developmental And Epileptic Encephalopathy 69 and Cockayne Syndrome B[56].

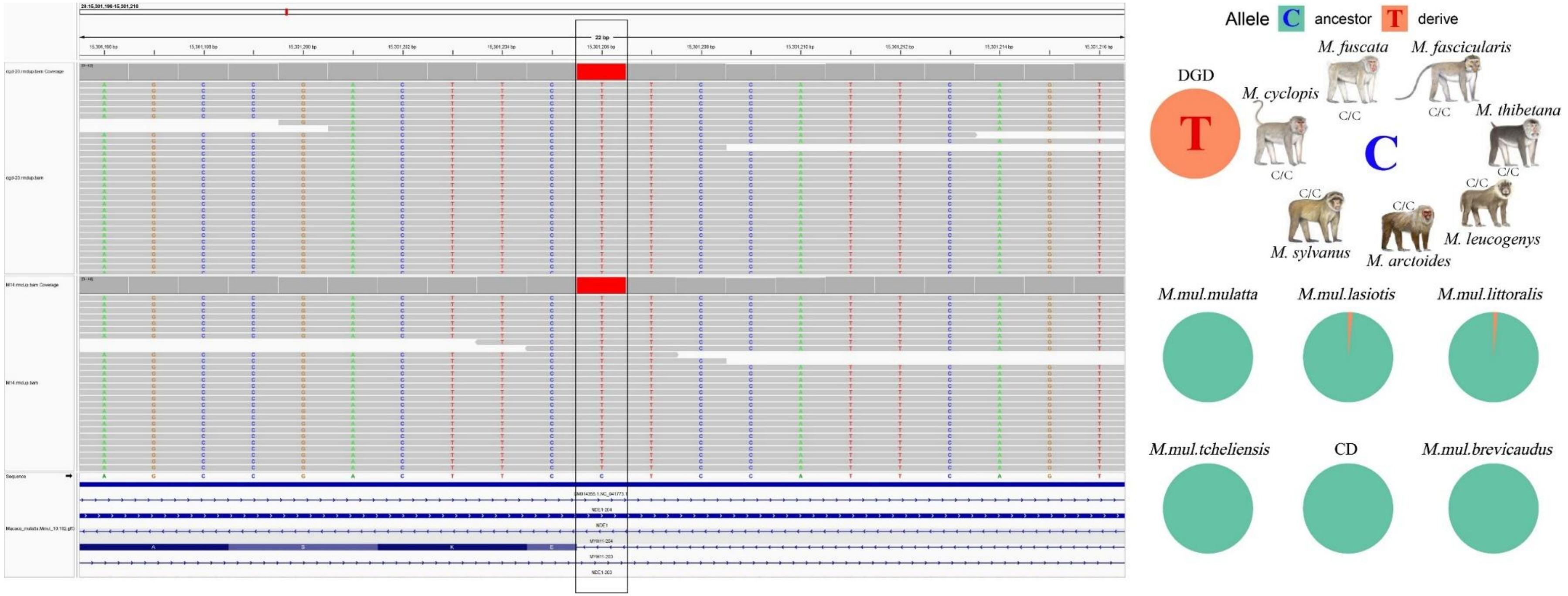


**Supplementary Figure 35.** NDE1: A putatively ultra-deleterious fixed splice acceptor loss-of-function (LoF) gene. This gene encodes a member of the nuclear distribution E (NudE) family of proteins. The encoded protein is localized at the centrosome and interacts with other centrosome components as part of a multiprotein complex that regulates dynein function. This protein plays an essential role in microtubule organization, mitosis and neuronal migration[57].

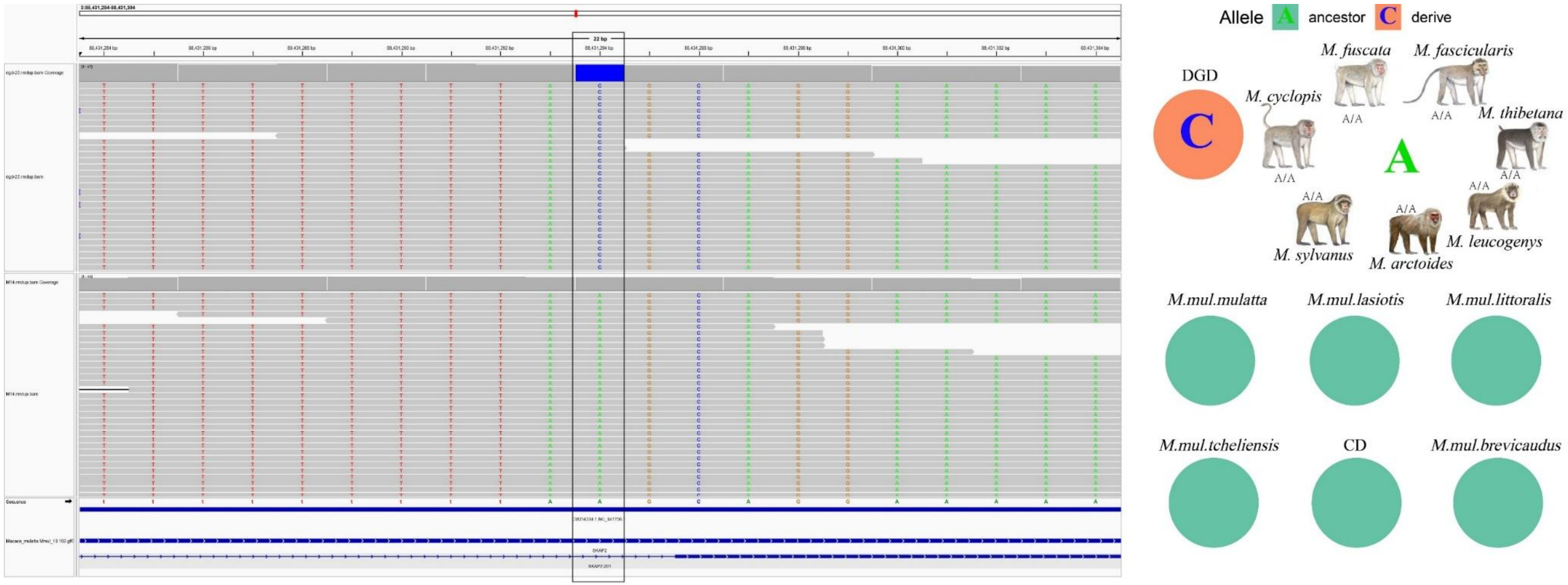


**Supplementary Figure 36.** SKAP2: A putatively ultra-deleterious fixed splice acceptor loss-of-function (LoF) gene. The protein encoded by this gene shares homology with Src kinase-associated phosphoprotein 1, and is a substrate of Src family kinases. It is an adaptor protein that is thought to play an essential role in the Src signaling pathway, and in regulating proper activation of the immune system[58].

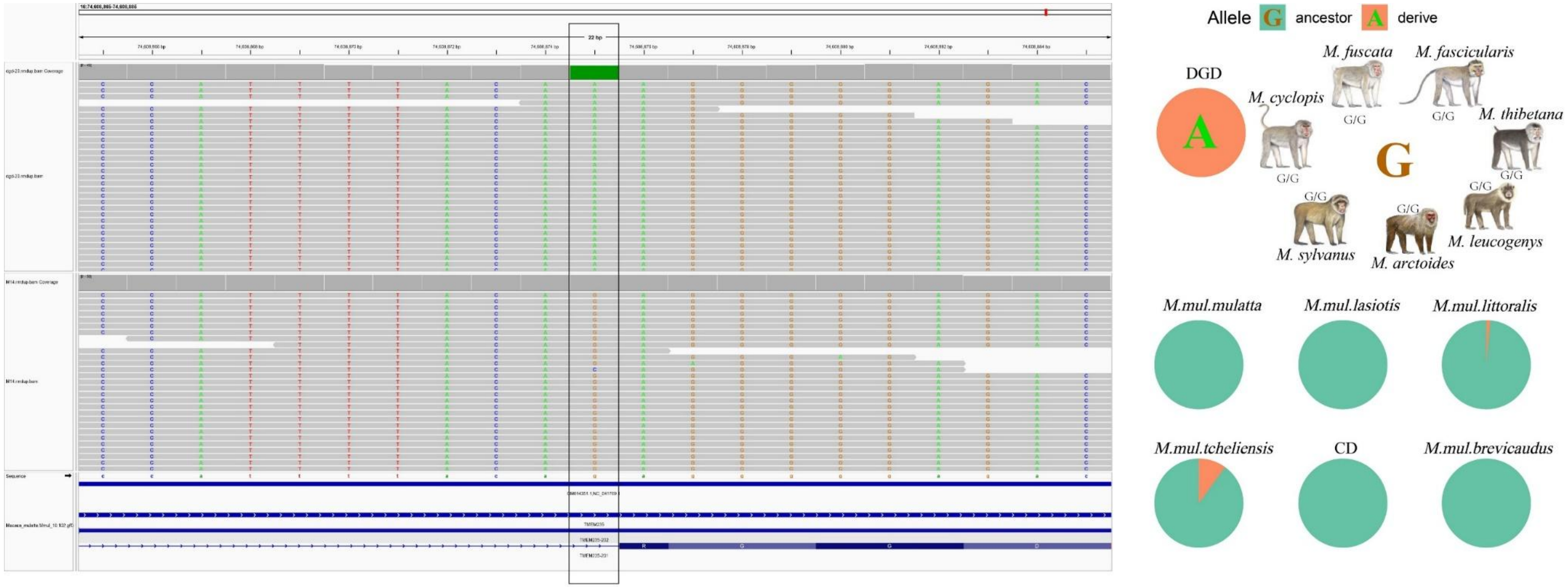


**Supplementary Figure 37.** *TMEM235:* A putatively ultra-deleterious fixed splice acceptor loss-of-function (LoF) gene. Diseases associated with *TMEM235* include cataract. RT-PCR analysis detected three variants of human *TMEM235* expressed in developing eye and central nervous system[59].

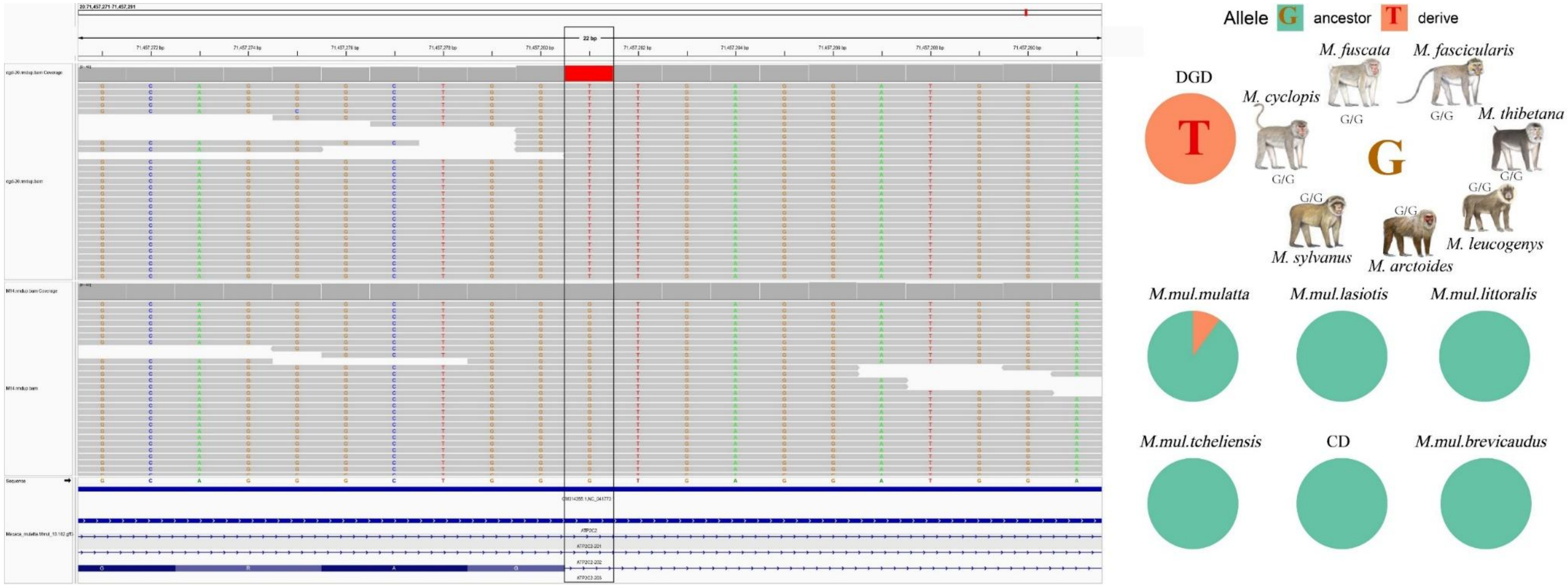


**Supplementary Figure 38.** ATP2C2: A putatively ultra-deleterious fixed splice doner loss-of-function (LoF) gene. Enables P-type calcium transporter activity and P-type manganese transporter activity. Predicted to be involved in calcium ion transmembrane transport; intracellular calcium ion homeostasis; and manganese ion transport. Diseases associated with ATP2C2 include Autism Spectrum Disorder and Dyslexia[60].

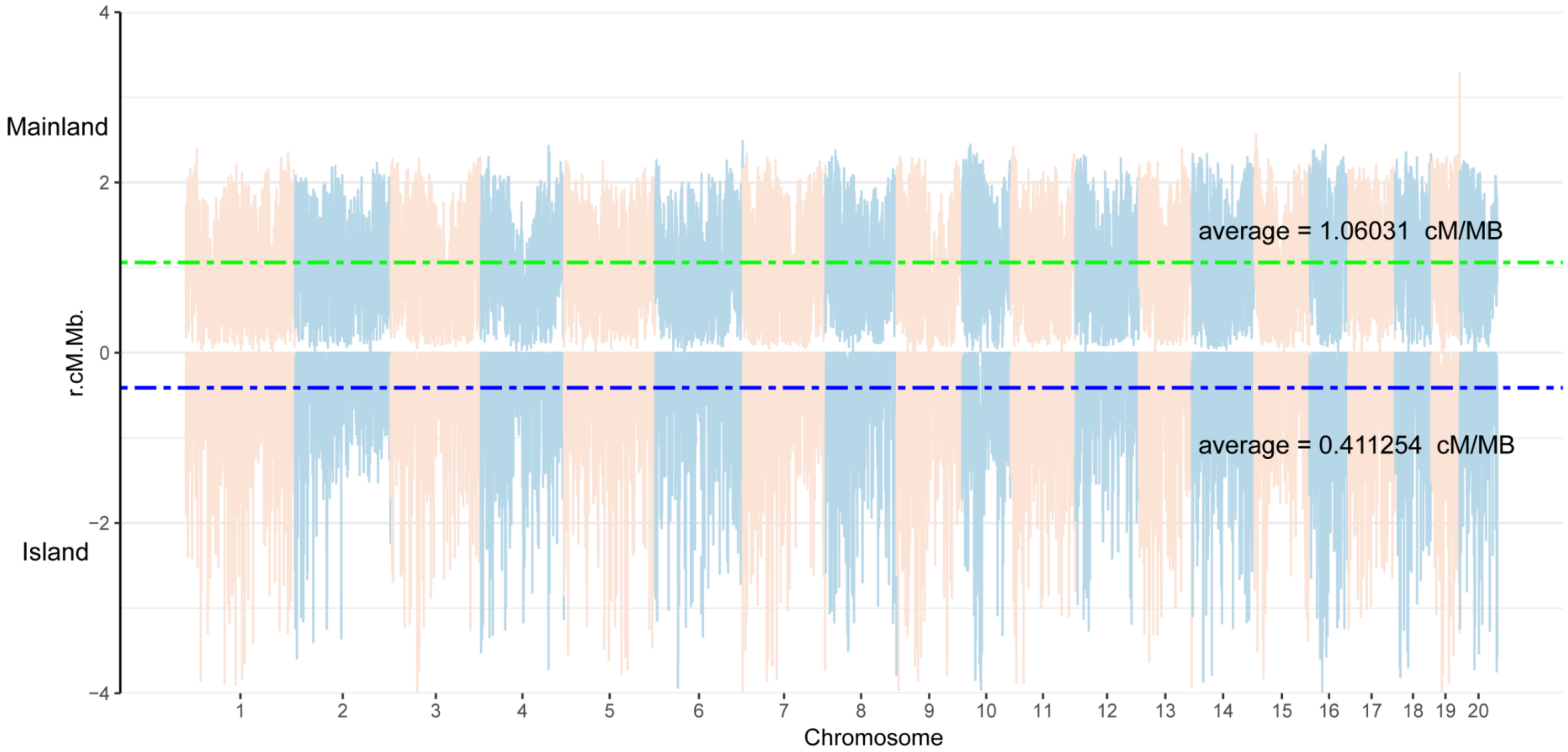


**Supplementary Figure 39.** Comparison of recombination rate maps for continents and islands calculated from FastEPRR. The recombination rate on the mainland is about 2.5 times that of the islands.

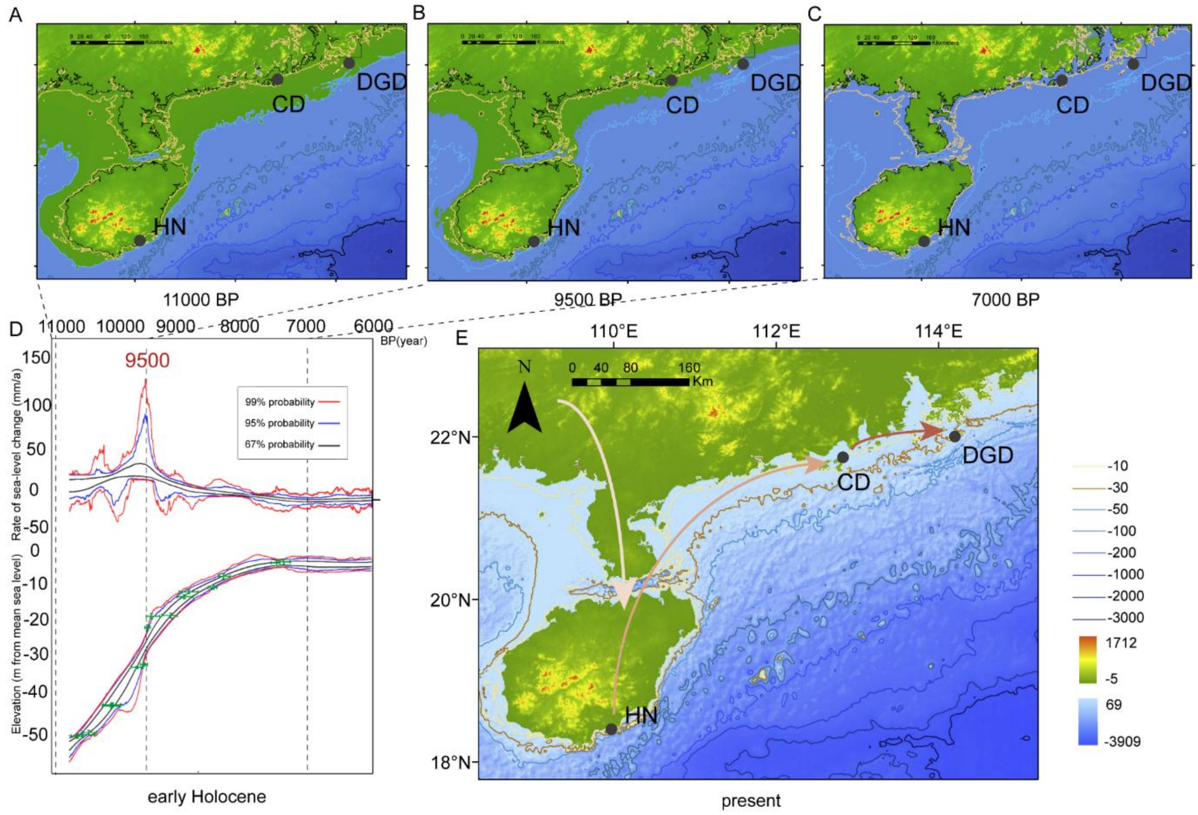


**Supplementary Figure 40.** Paleogeographic map of the Northwest Pacific continental shelf depicting reconstructed sea level contours from 11,000 to 7,000 years ago and present-day bathymetry/topography. Labeled paleocoastlines and sea level positions: (A) 11,000 cal yr BP, (B) 9,500 cal yr BP, (C) 7,000 cal yr BP, (D) Adapted from source, showing rates of sea-level change derived from Monte Carlo simulations using new and pre-existing sea-level index points from the Pearl River deltaic basin[61]. Adapted from source, showing elevations of new sea-level index points after correction for tectonic movement and sediment compaction. Superimposed sea-level probability intervals are from Monte Carlo simulations[61]. (E) Present day. Arrows indicate proposed dispersal routes of macaques. Initial dispersal occurred widely along the southern paleocoastline prior to the mid-Holocene highstand. The rapid sea-level rise ~9,500 cal yr BP flooded land bridges, cutting off dispersal routes and isolating the macaque population on DGD Island, providing natural conditions for speciation.

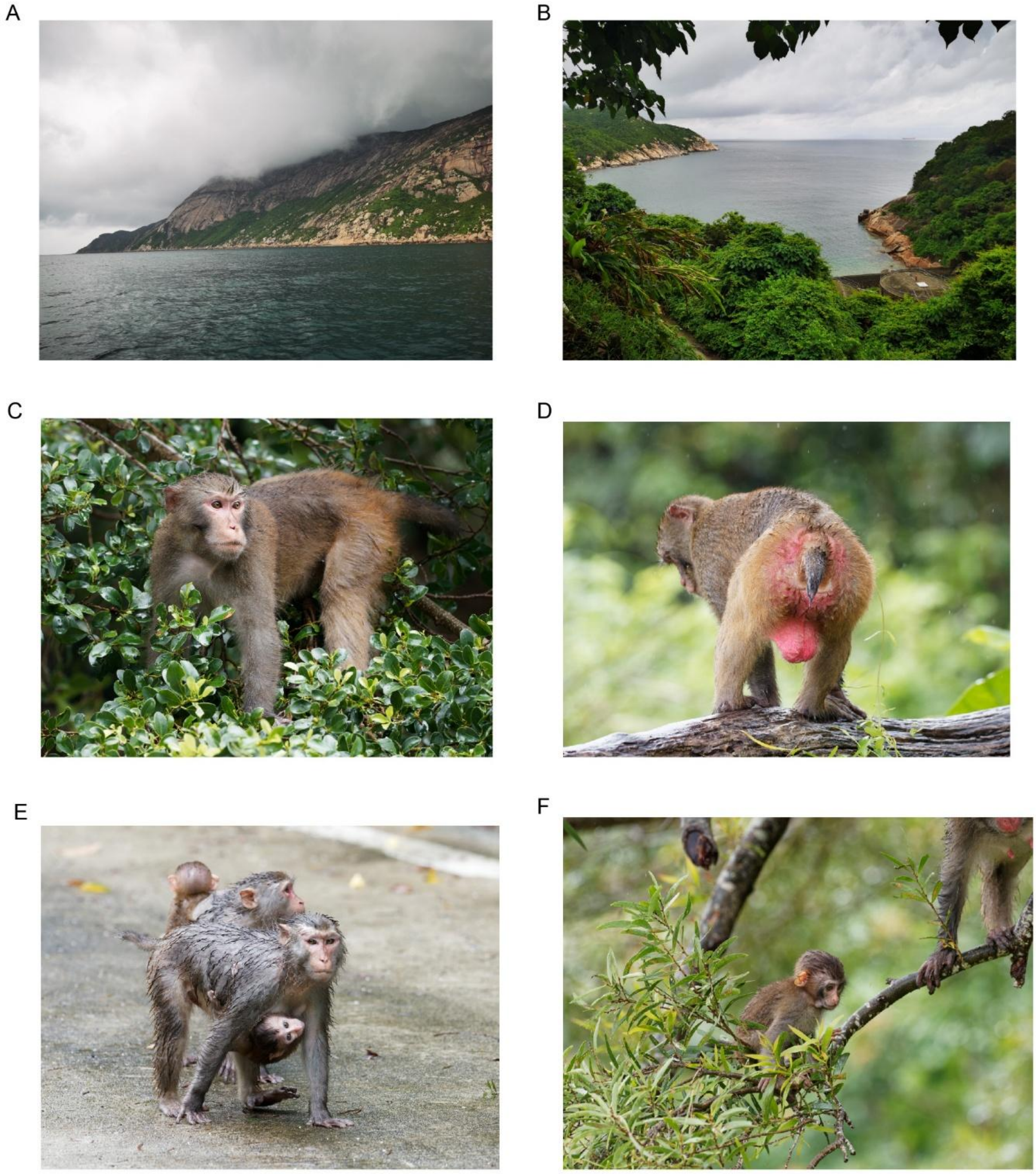


**Supplementary Figure 41.** Habitat and photographs of DGD macaques, South China Sea. A–B, Island habitat characterized by dense subtropical evergreen broadleaf forest (> 96% coverage). Dangan Island (13.4 km² total area) sustains perennial freshwater streams and hosts ~200 permanent human residents. C–F, Morphological documentation of free-ranging

DGD macaques. (C) Adult male foraging; (D) Juvenile with characteristic short tail (1/3 body length) resting on wood outcrop; (E) Family troop crossing road; (F) Female-infant dyad displaying pelage uniformity under monsoon forest understory.
